# Tumor mutational landscape is a record of the pre-malignant state

**DOI:** 10.1101/517565

**Authors:** Kirsten Kübler, Rosa Karlić, Nicholas J. Haradhvala, Kyungsik Ha, Jaegil Kim, Maja Kuzman, Wei Jiao, Sitanshu Gakkhar, Kent W. Mouw, Lior Z. Braunstein, Olivier Elemento, Andrew V. Biankin, Ilse Rooman, Mendy Miller, Wouter R. Karthaus, Christopher D. Nogiec, Edouard Juvenson, Edward Curry, Mari Mino- Kenudson, Leif W. Ellisen, Robert Brown, Alexander Gusev, Cristian Tomasetti, Martijn P. Lolkema, Neeltje Steeghs, Carla van Herpen, Hong-Gee Kim, Hwajin Lee, Kristian Vlahoviček, Bradley E. Bernstein, Charles L. Sawyers, Katherine A. Hoadley, Edwin Cuppen, Amnon Koren, Peter F. Arndt, David N. Louis, Lincoln D. Stein, William D. Foulkes, Paz Polak, Gad Getz, on behalf of the PCAWG Pathology and Clinical Correlates Working Group, and the ICGC/TCGA Pan-Cancer Analysis of Whole Genomes Network

**Affiliations:** Broad Institute of MIT and Harvard, Cambridge, Massachusetts, USA; Center for Cancer Research, Massachusetts General Hospital, Charlestown, Massachusetts, USA; Harvard Medical School, Boston, Massachusetts, USA; Bioinformatics Group, Division of Molecular Biology, Department of Biology, Faculty of Science, University of Zagreb, Zagreb, Croatia; Biomedical Knowledge Engineering Laboratory, Seoul National University, Seoul, South Korea; Interdisciplinary Program of Medical Informatics, College of Medicine, Seoul National University, Seoul, South Korea; Adaptive Oncology Initiative, Ontario Institute for Cancer Research, Toronto, Ontario, Canada; Canada’s Michael Smith Genome Sciences Centre, BC Cancer Agency, Vancouver, British Columbia, Canada; Department of Radiation Oncology, Brigham & Women’s Hospital and Dana Farber Cancer Institute, Boston, Massachusetts, USA; Department of Radiation Oncology, Memorial Sloan Kettering Cancer Center, New York, New York, USA; Institute for Computational Biomedicine, Weill Cornell Medical College, New York, New York, USA; Caryl and Israel Englander Institute for Precision Medicine, Weill Cornell Medical College, New York, New York, USA; Sandra and Edward Meyer Cancer Center, Weill Cornell Medical College, New York, New York, USA; Wolfson Wohl Cancer Research Centre, Institute of Cancer Sciences, University of Glasgow, Garscube Estate, Switchback Road, Bearsden, Glasgow, UK; Laboratory of Medical and Molecular Oncology, Oncology Research Center, Vrije Universiteit Brussel, Laarbeeklaan, Brussels, Belgium; Human Oncology and Pathology Program, Memorial Sloan Kettering Cancer Center, New York, New York, USA; Department of Oncological Sciences, Tisch Cancer Institute, Icahn School of Medicine at Mount Sinai, New York, New York, USA; McGill University Health Centre, McGill University, Montreal, Quebec; Department Surgery and Cancer, Imperial College London, London, United Kingdom; Department of Pathology, Massachusetts General Hospital, Boston, Massachusetts, USA; Institute of Cancer Research, Sutton, United Kingdom; Division of Genetics, Department of Medicine, Brigham and Women’s Hospital, Boston, Massachusetts, USA; Dana Farber Cancer Institute, Harvard Medical School, Boston, Massachusetts, USA; Sidney Kimmel Cancer Center at Johns Hopkins, The Johns Hopkins Medical Institutions, Baltimore, Maryland, USA; Division of Biostatistics and Bioinformatics, Department of Oncology, The Johns Hopkins Medical Institutions, Baltimore, Maryland, USA; Department of Biostatistics, The Johns Hopkins Bloomberg School of Public Health, Baltimore, Maryland, USA; Department of Medical Oncology, Erasmus MC Cancer Institute, Erasmus University Medical Center, Rotterdam, The Netherlands; Department of Medical Oncology, The Netherlands Cancer Institute, Antoni van Leeuwenhoek, Amsterdam, The Netherlands; Radboud University Medical Center, Nijmegen, the Netherlands; Dental Research Institute, School of Dentistry, Seoul National University, Seoul, South Korea; Howard Hughes Medical Institute, Chevy Chase, Maryland, USA; Department of Genetics, Lineberger Comprehensive Cancer Center, The University of North Carolina at Chapel Hill, Chapel Hill, North Carolina, USA; Hartwig Medical Foundation, Science Park 408, Amsterdam, The Netherlands; Center for Molecular Medicine and Oncode Institute, University Medical Center Utrecht, Heidelberglaan 100, Utrecht, The Netherlands; Department of Molecular Biology and Genetics, Cornell University, Ithaca, New York, USA; Department of Computational Molecular Biology, Max Planck Institute for Molecular Genetics, Ihnestr. 63/73, 14195 Berlin, Germany; Department of Molecular Genetics, University of Toronto, Toronto, Ontario, Canada; Department of Human Genetics, McGill University, Montreal, Quebec, Canada; Lady Davis Institute for Medical Research and Research Institute McGill University Health Centre, McGill University, Montreal, Quebec, Canada; Department of Genetics and Genomic Sciences, Icahn School of Medicine at Mount Sinai, New York, New York, USA; Department of Pathology, Tisch Cancer Institute, Icahn School of Medicine at Mount Sinai, New York, New York, USA; Department of Medicine, Hematology and Medical Oncology, Tisch Cancer Institute, Icahn School of Medicine at Mount Sinai, New York, New York, USA

## Abstract

Chromatin structure has a major influence on the cell-specific density of somatic mutations along the cancer genome. Here, we present a pan-cancer study in which we searched for the putative cancer cell-of-origin of 2,550 whole genomes, representing 32 cancer types by matching their mutational landscape to the regional patterns of chromatin modifications ascertained in 104 normal tissue types. We found that, in almost all cancer types, the cell-of-origin can be predicted solely from their DNA sequences. Our analysis validated the hypothesis that high-grade serous ovarian cancer originates in the fallopian tube and identified distinct origins of breast cancer subtypes. We also demonstrated that the technique is equally capable of identifying the cell-of-origin for a series of 2,044 metastatic samples from 22 of the tumor types available as primaries. Moreover, cancer drivers, whether inherited or acquired, reside in active chromatin regions in the respective cell-of-origin. Taken together, our findings highlight that many somatic mutations accumulate while the chromatin structure of the cell-of-origin is maintained and that this historical record, captured in the DNA, can be used to identify the often elusive cancer cell-of-origin.

## INTRODUCTION

One important, but largely unanswered, question in cancer biology is the identity of the normal cell (i.e., cell-of-origin, COO) from which the tumor is derived. While tumor morphology usually bears some resemblance to the originating tissue, histological similarity is often too broad a parameter to distinguish between molecularly and clinically distinct cancer subtypes^1^. Accordingly, knowledge of the precise nature of the cancer cell type of origin can help better understand the potential of certain normal cell types to transform and initiate cancer, as well as the association of the COO with tumor subtypes and treatment sensitivities. Current knowledge of the cancer COO is mostly based on mouse models^2,3^. However, studies that use human tissue are particularly valuable since they directly capture the neoplastic process and overcome limitations introduced by interspecies differences^4^.

It is well established that neoplastic transformation is driven by somatic mutations^5^. A subset of mutations is present at the time of initial cancer growth and common to all cancer cells (‘clonal’). Alterations in genes (‘drivers’) critical to the development of cancer arise on a background of random mutations (‘passengers’) that accumulate over time. It has been found that the set of driver genes vary substantially across tumor types^6^. Likewise the phenotypic effects of defects in most inherited cancer susceptibility genes (e.g., *BRCA1/2* genes) are limited to specific tissue types^7^. Both these observations suggest a cell type-specific vulnerability to mutations.

One major determinant of a cell phenotype is its chromatin structure, which differs considerably across tissue types and cell differentiation stages^8,9^. The chromatin structure, in turn, is governed by epigenetic processes, including DNA and histone modifications, which greatly influence the rate at which background mutations accumulate in the cell.

In our previous proof-of-concept study^10^, we developed a framework for understanding how different epigenetic features are associated with mutagenesis in a cell type-specific manner. This work described a method that quantifies the ability to predict the mutational density along the cancer genome from the profile of epigenetic modifications in normal cell types. Two main observations were used to guide our discovery: (i) mutations are not evenly distributed along chromosomes and across tumor types^11,12^; and (ii) mutation densities are associated with regional histone modifications, DNA accessibility and DNA replication timing^11,13–15^.

In the present study, we applied our method to identify the likely COO across multiple cancer types and asked whether the characterization of cancer origins from the distribution of somatic mutations is universally applicable to all common cancer types and subtypes. The availability of the large compendium of tumor mutation and normal chromatin data enabled us to extend our previous investigation in three key aspects: (i) to quadruple the number of cancer types from 8 to 32 and increase more than tenfold the number of individual samples analyzed from 173 to 2,550; (ii) to include tumors known to develop along a metaplasia-carcinoma sequence (i.e., conversion of one differentiated cell type into another differentiated cell type before becoming a *bona fide* cancer); and (iii) to study cancers that arise in the same organ but manifest as distinct subtypes. In addition, we demonstrate that the chromatin structure of the COO can also be detected by analyzing the mutational distribution in metastases, a result that can have clinical implications. Finally, we show that there is an association between regions of open chromatin in the relevant COO and genomic loci of cancer risk alleles and somatic driver genes.

## RESULTS

To search for the COO across a broad panel of tumor types, we utilized the ICGC/TCGA Pan-cancer Analysis of Whole Genomes (PCAWG) mutation data^16^ across 2,550 samples from 32 cancer types (**Fig. 1 left**, **Extended Data Table 1**). Furthermore, we gathered 98 complete chromatin profiles of normal tissue types from the Roadmap Epigenomics Consortium^17^, the Encyclopedia of DNA Elements (ENCODE^18^) and the International Human Epigenome Consortium (IHEC^19^) as well as 6 partial profiles from additional publications^20,21^. Overall, ChIP-sequencing (ChIP-seq) data were available for 104 normal tissue types including distinct cellular differentiation stages from the brain, blood, breast and prostate (**Fig. 1 right**, **Extended Data Fig. 1**, **Extended Data Table 2**). For our main analysis we chose six histone modifications due to their broad availability across almost all tissue types (see **Methods** for details).

**Figure 1.**
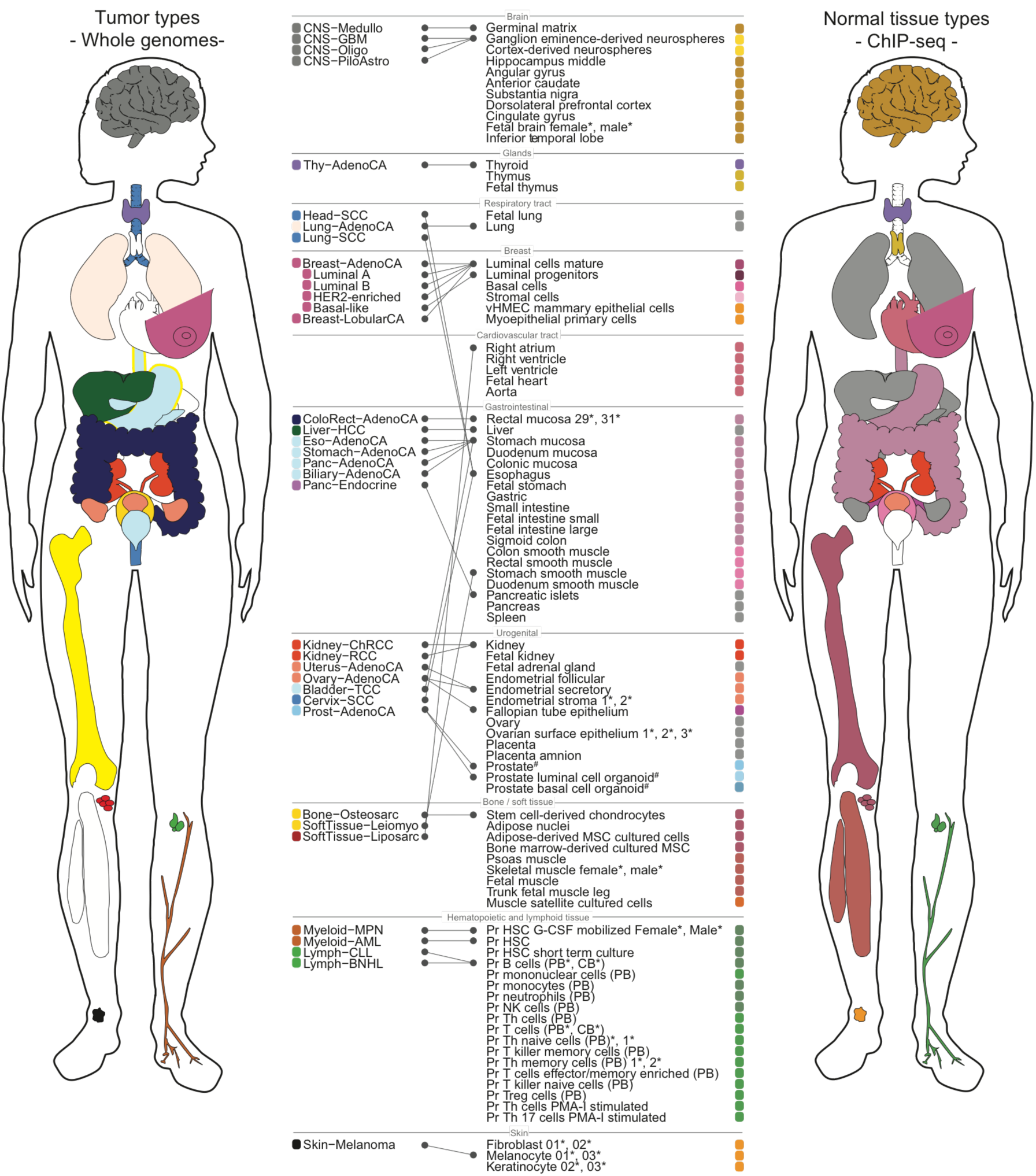
Overview of tumor and normal tissue types. Schematic illustrations depict the anatomical sites of 32 cancer types (left) and 104 normal tissue types (right), color-coded according to **Extended Data Tables 1, 2**. Blank spaces throughout the cartoons depict unavailability of appropriate data. Cases with biological replicates are indicated by an asterisk with numbers depicting each individual replicate; organs for which epigenetic features are available of the respective tumor type are marked with a hashtag. The best-matched normal tissue type for each tumor type as detected by Random Forest regression is indicated by joined lines. Random Forest regression was additionally performed on a subset of tissues for high-grade serous Ovary-AdenoCA and Prost-AdenoCA identifying a best match on a more detailed level. Abbreviations: vHMEC, variant human mammary epithelial cells; Pr, primary; Th cells, T helper cells; PB, peripheral blood; CB; cord blood; HSC, hematopoietic stem cells; MSC, mesenchymal stem cells; PMA, phorbol myristate acetate; NK cells, natural killer cells; Treg cells, regulatory T cells.

The number of individual tumors varied across cancer types, from a minimum of 10 samples for acute myeloid leukemia (Myeloid−AML) to a maximum of 314 samples for hepatocellular carcinoma (Liver-HCC). Likewise, the mutational burden varied across cancer types from a high mutational burden in melanoma to a low mutational burden in non-diffuse glioma. To account for these differences in sampling and mutational burden between cancer types, we aggregated the somatic mutation data across all samples from a given cancer type and generated an ‘aggregated tumor profile’ per tumor type. However, to quantify certain aspects of our analysis, we also used individual tumors when appropriate.

Based on our earlier work^10^, we used random forest regression to model the distribution of somatic mutations in 1Mb windows based on the density of ChIP-seq reads derived from each of the normal tissues. To quantify the accuracy of our models, we measured the correlation between the number of predicted and observed mutations (R^2^ or % variance explained). While previously we trained one multivariate model with the chromatin marks from all tissue types and identified the most significant coefficient, here we trained, for each tumor type, 98 different models using the chromatin profiles of each normal tissue type. We then determined the candidate COO by identifying the model with the best predictive power. To train the models, we used the same six core ChIP-seq tracks available for all Roadmap and IHEC tissues to ensure comparability between models and cancer types. This modified approach allowed us to add additional normal tissue types and easily identify the best-matching tissue. For example, the two best-matching chromatin profiles for uterine adenocarcinoma (Uterus-AdenoCA) are derived from endometrial cells (follicular and secretory), and the best-matching normal tissue for pancreatic ductal adenocarcinoma (Panc-AdenoCA) is stomach mucosa (**Fig. 2**).

**Figure 2.**
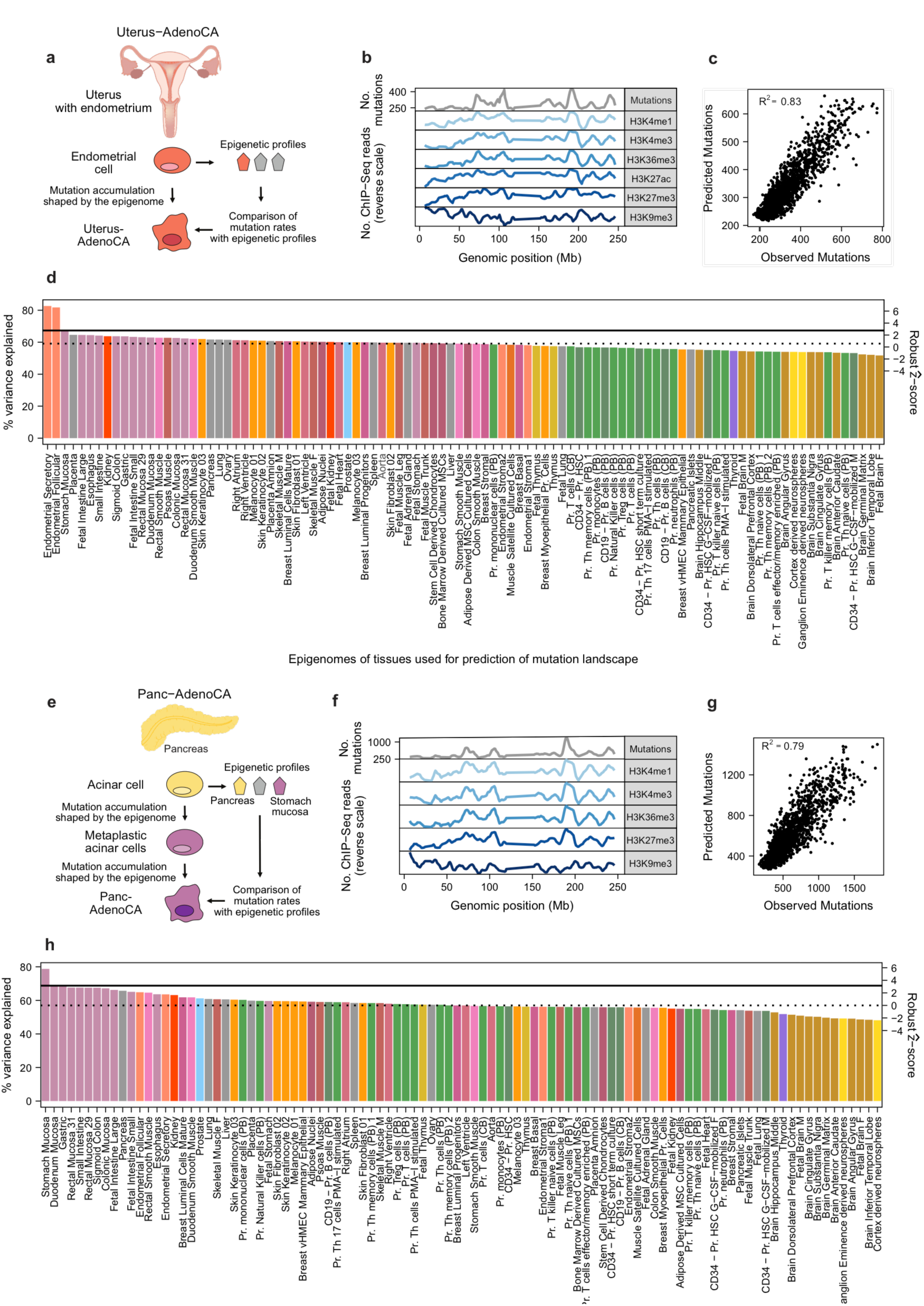
The chromatin composition of the COO is a major determinant of cancer mutation distribution and provides insights regarding tumor origins. (**a,e**) Cartoons depict our strategy using normal chromatin profiles and cancer mutation profiles in Uterus-AdenoCA and Panc-AdenoCA. (**b,f**) Mutation counts of aggregated tumor profiles in 1Mb windows are shown alongside six histone modification profiles derived from the respective best match (in normalized reverse scale; peaks correspond to less accessible chromatin and vice versa). (**c,g**) Random Forest regression was used to predict the tumor mutation distribution from chromatin marks of normal cells. Scatterplots show the correlation between observed mutation counts and those predicted from chromatin marks. The overall prediction accuracy is reported by the average R^2^ value between the predicted and observed mutation profiles across the 10-fold cross validation. (**d,h**) For each cancer type, individual models were trained on chromatin marks from 98 cell types. The cell type that fit the model with the highest prediction accuracy was defined as the best match. Solid horizontal lines indicate performance of the second-best model based on chromatin marks from histologically unrelated cells; dashed horizontal lines depict the variance explained based on chromatin marks that represent the median tissue. Bars are colored according to **Extended Data Table 2**. The right-hand (secondary) axis shows the robust ẑ-score.

### The best-matching cell type for most cancer types is the expected normal cellular counterpart

By applying our methodology, we identified the best-matching normal cell type (of the 98) to each of the aggregated mutational profiles of 32 tumor types. In 23 of the 32 tumor types, we matched the chromatin profiles of their expected normal cellular counterparts (e.g., melanoma best matched melanocytes) We found that the best-matched normal cellular counterpart explained significantly more variance than the next-best histologically unrelated cell type in 20 out of the 23 tumor types (all *p*-values < 0.03, WMW test; solid lines in **Fig. 3a**, **Extended Data Fig. 2**, **Extended Data Table 1**).

**Figure 3.**
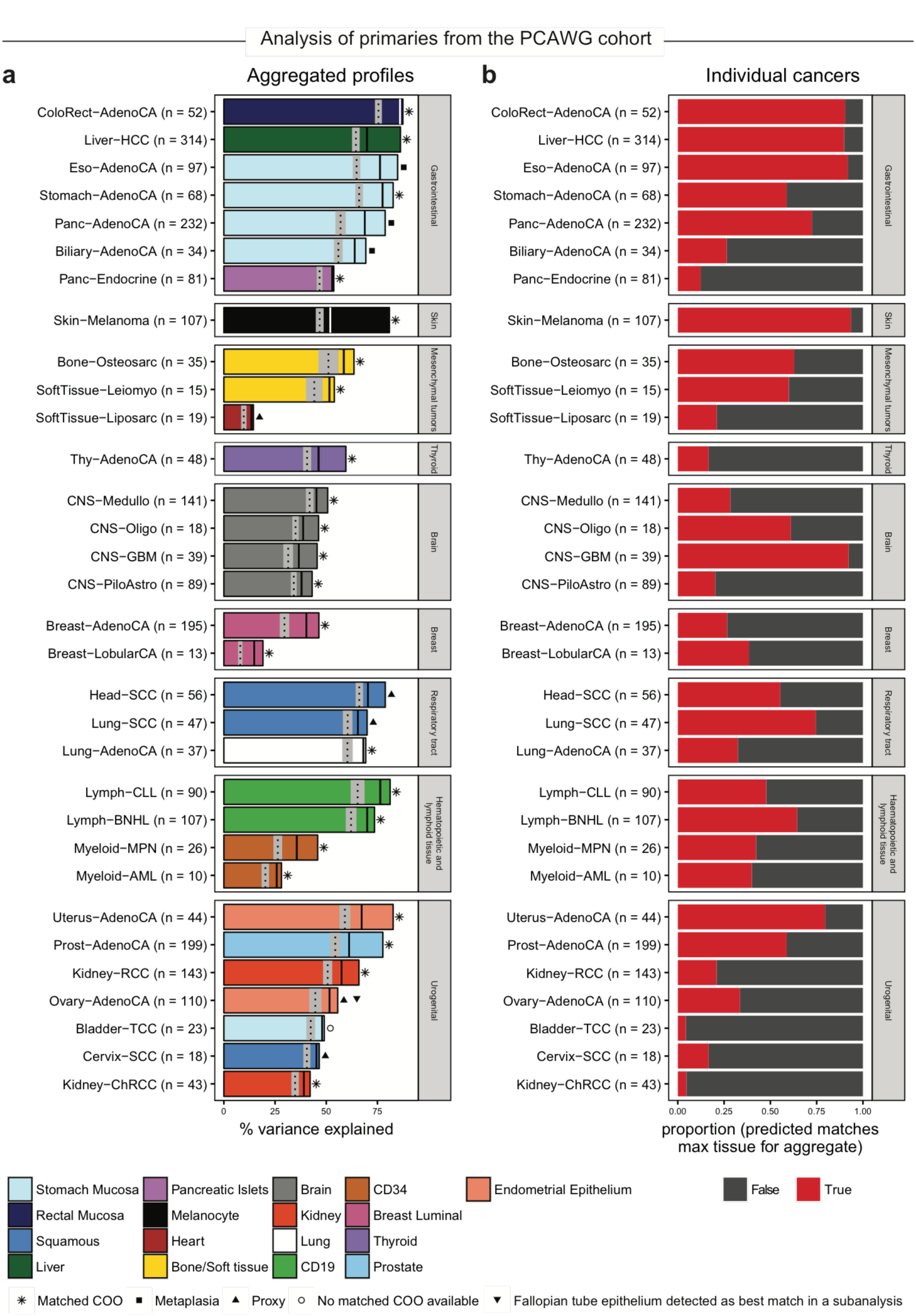
Chromatin marks predict tumor origins for aggregated and individual tumor mutation profiles. (**a**) For each cancer type, we trained models on histone modifications from 98 normal cell types to predict the mutation density in 1Mb windows of aggregated tumor profiles. Bar plots show the variance explained by the best performing model, identified via ten-fold cross validation. Solid vertical lines indicate performance of the second-best model based on chromatin marks from histologically unrelated cells; dashed vertical lines depict the variance explained based on chromatin marks that represent the median tissue. The grey area indicates ± Median Absolute Deviation (MAD). Bars are colored according to the best match. Symbols identify types of best matches. (**b**) Barplots show the proportion of individual tumors (red) in which the prediction on an individual level matches the prediction based on aggregated tumor profiles. In some cases, chromatin profiles were grouped (see **Methods**, **Extended Data Fig. 1**)

Not all best-matched models (among the 23 tumor types) had the same predictive power, with variance explained ranging between 19 and 87% (median = 69.6%, 95% CI: 51-69%). Consistent with our previous work^10^ the overall mutation frequency in the aggregated profile greatly influenced the ability to build an accurate model. Aggregated profiles with above the median mutation frequency (≥ 184/Mb) typically had above the median variance explained (≥62%; *p* = 0.01, Fisher's exact test). We next checked whether the clonal status of mutations had an influence on the variance explained. First, we compared predictions based only on the clonal mutations^22^ (on average 70.8% of somatic mutations) to predictions based on all the mutations (clonal and sub-clonal) and found similar values of variance explained (**Extended Data Fig. 3a**). However, when we contrasted predictions based on clonal mutations to predictions based on sub-clonal mutations (controlling for the number of mutations), the variance explained by the models that used clonal mutations was typically significantly higher than those based on sub-clonal mutations, suggesting that earlier somatic mutations are that ones that capture the information about the COO (**Extended Data Fig. 3b**).

After analyzing tumor types based on their aggregated tumor profiles, we tested whether the COO discovered above for the 23 tumor types could also be identified using individual tumors, even in the setting of far fewer mutations available for modeling (**Fig. 3b**). Gratifyingly, the COO determined by the aggregated tumor profile was among the top three best matches for more than 50% of individual tumors in 15/23 cancer types (**Extended Data Fig. 4**). As expected, these 15 tumor types were enriched with tumors containing higher mutation frequencies (*p* = 1.2×10^−26^, WMW test; **Extended Data Fig. 5**). Likewise, a larger number of individual tumors matched the expected normal cell in tumor types with higher variance explained when using the aggregated tumor profiles (r = 0.59, *p* = 4×10^−4^, Pearson correlation coefficient; **Extended Data Fig. 6**). Taken together, our data indicate that the memory of historical cell lineage is retained by the tumor and can be discovered from the somatic mutation density of tumors identifying the often elusive cancer COO.

**Figure 4.**
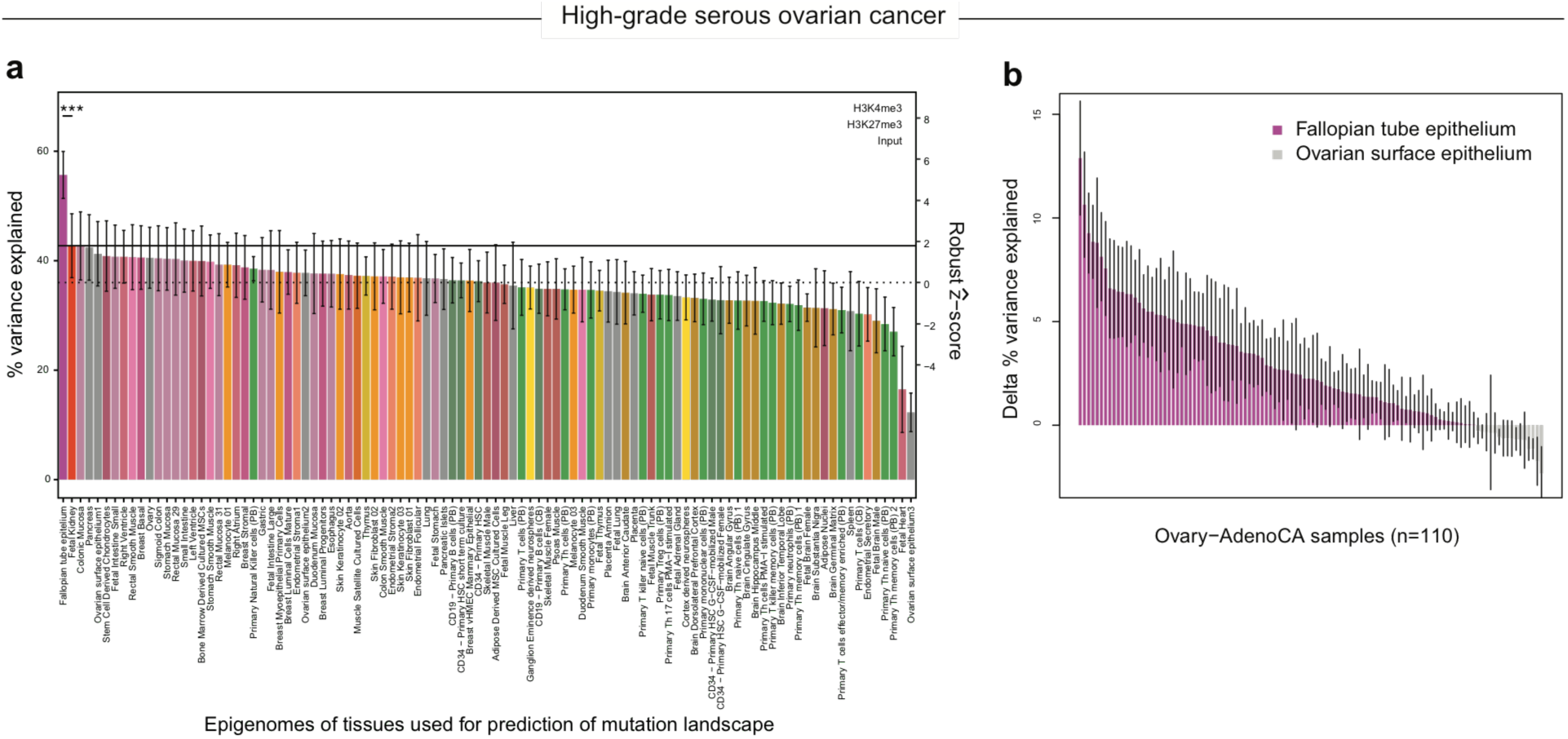
High-grade serous ovarian cancer best matches fallopian tube epithelium. (**a**) Random Forest regression models for the prediction of mutation density in 1Mb windows of aggregated tumor profiles of high-grade serous Ovary-AdenoCA were trained on an extended set of 99 tissues for which three ChIP-seq experiments were available. The best match was identified corresponding to the model with the highest prediction accuracy; *p*-values were obtained using the paired Wilcoxon-Mann-Whitney test for the comparison of R^2^ values from the 10-fold cross validation (***, *p* < 0.001; dots represent ten-fold cross-validation values). (**b**) Random Forest regression models for individual high-grade serous Ovary-AdenoCA tumors were trained on fallopian tube epithelium and three different ovarian surface epithelia. Color indicates the best match. Depicted is the difference of the means of the R squared values from the 10-fold cross validation and the error of this difference calculated from the standard error of the means.

**Figure 5.**
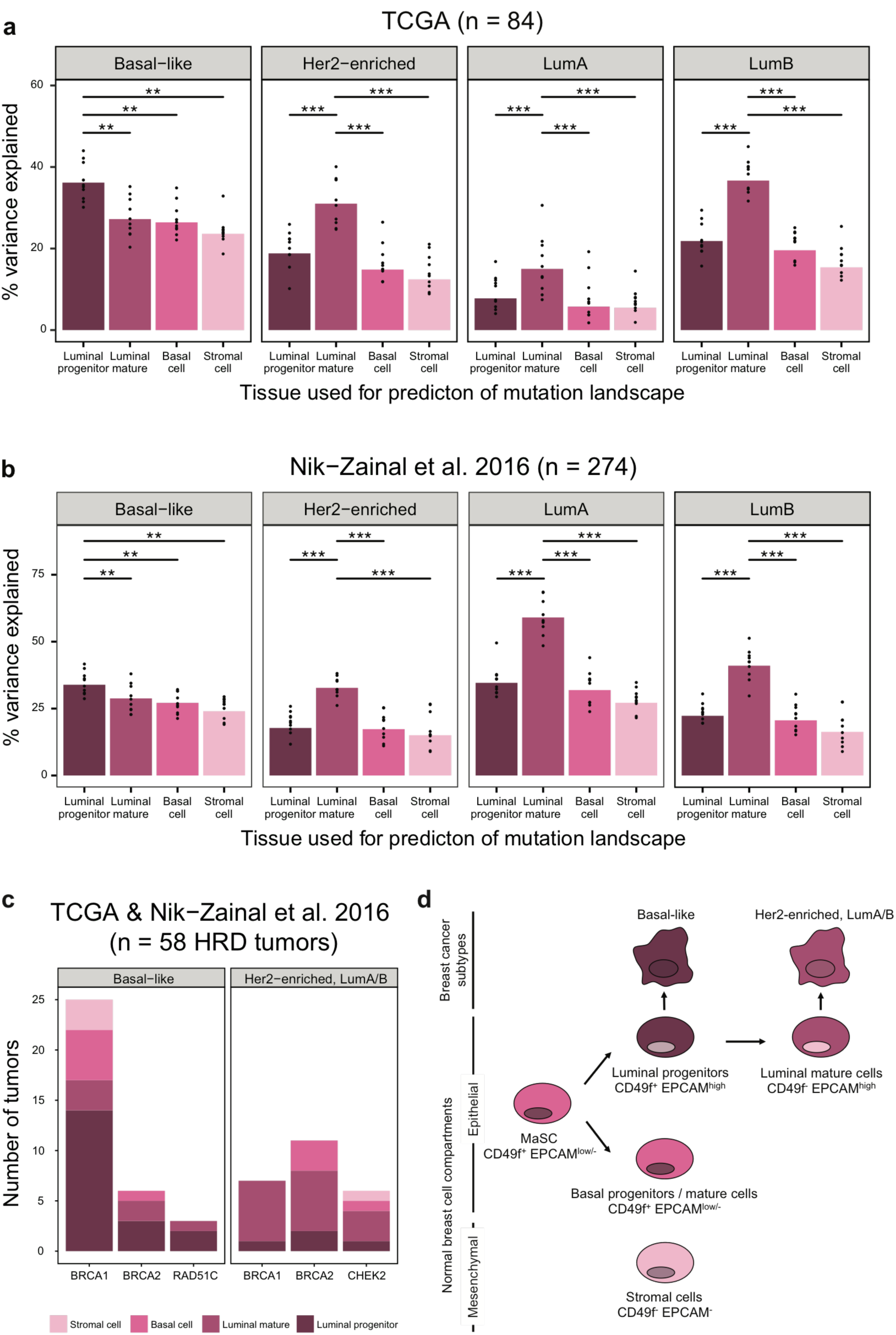
Breast cancer subtypes arise in different mammary cell types. (**a**, **b**) Random Forest regression models for the prediction of mutation density in 1Mb windows of aggregated tumor profiles of distinct molecular breast cancer subtypes were trained on chromatin data from four normal mammary cell types. Using two independent data sets, the best match for each subtype was determined, corresponding to the model with the highest prediction accuracy; *p*values were obtained using the paired Wilcoxon-Mann-Whitney test for the comparison of R^2^ values from the 10-fold cross validation (**, *p* < 0.01; ***, *p* < 0.001; dots represent ten-fold cross-validation values). (**c**) A subset of 58 breast cancer samples with HRD was analyzed for their best matching normal cell using chromatin profiles from four normal breast cells. Colors indicate the number of tumors that best match one of the mammary cell subtypes. (**d**) Schematic representation of the mammary epithelial cell hierarchy and its association with distinct breast cell types. Only breast cancer subtypes and normal mammary cells available in our cohort are depicted. The expression of selected cell surface markers represents the markers used for isolating normal breast cell subpopulations.

**Figure 6.**
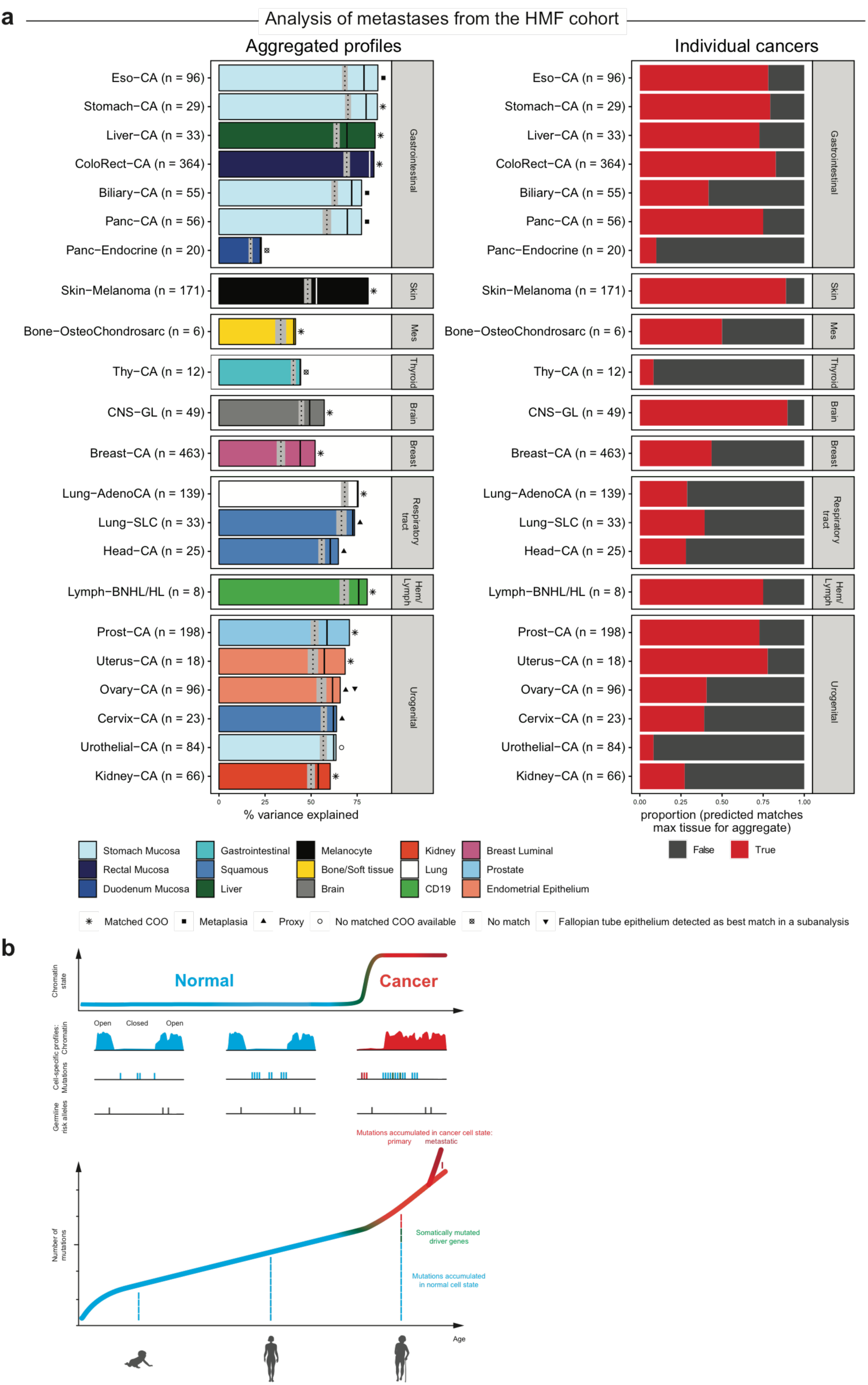
The cell type of origin can be inferred from metastatic cancer samples. (**a, left**) Models were trained on histone modifications from 98 normal cell types to predict the mutation density aggregated metastatic cancer profiles. Bar plots show the variance explained by the best-performing model, identified via ten-fold cross validation. Solid vertical lines indicate performance of the second-best model based on chromatin marks from histologically unrelated cells; dashed vertical lines depict the variance explained based on chromatin marks that represent the median tissue. The grey area indicates ± Median Absolute Deviation (MAD). Bars are colored according to the best match. Symbols identify types of best matches. Abbreviations: Mes, Mesenchymal tumors; Hem/Lymph, Hematopoietic and lymphoid tissue. (**a, right**) Bar plots depict the proportion of individual metastatic samples (red) in which the prediction on an individual level matches the prediction of the aggregated metastatic cancer profile. In some cases, chromatin profiles were grouped (see **Methods**, **Extended Data Fig. 1**). (**b**) Schematic illustrations depict the qualitative change of the cell type-specific chromatin state from normal to cancer (top), cell type-specific chromatin and mutation profiles alongside germline risk alleles along one genomic region (middle) and the accumulation of somatic mutations throughout the human lifespan (bottom). Blue color indicates a normal cell state, red color indicates a cancerous cell state.

### Histologically related tissue types serve as a proxy for the cell-of-origin

Next, we sought to understand and define the cellular origins of the remaining 9 tumor types (out of the aggregated 32) that did not match their expected normal cellular counterpart. In 5 of the 9 tumor types, our analysis indicated that the best match was a histologically related (or ‘proxy’) tissue type (**Fig. 3a**), consistent with the previous observation of similar chromatin profiles^17^. The three squamous differentiated tumor types cervix (Cervix−SCC), head and neck (Head-SCC) and lung (Lung-SCC) do not have a direct normal tissue counterpart Roadmap, ENCODE or IHEC projects. However, their best match is another squamous-lined tissue, the esophagus that we did have epigenetic profiles for (**Extended Data Fig. 2**). This convergence is consistent with previous reports of similar molecular profiles including expression, copy number changes and mutations, for these tumor types and the observation that more molecular features are shared across squamous tumors than with adenocarcinomas that arise in their respective organs^23,24^. Likewise, the mesenchymal tumor type liposarcoma (SoftTissue-LipoSarc), which not have a direct counterpart (adipose tissue) in our data set, best matched the right atrium, another mesenchymal cell type. Thus, we are able to identify a normal cell type that is related to the expected COO in cases wherein the exact normal cell type did not exist in our data set of normal tissues, but a closely related tissue was available.

### Fallopian tube best matches high-grade serous ovarian cancer

Of particular interest was our finding in high-grade serous ovarian cancer (Ovary-AdenoCA). When using the original 98 tissue types, the best match was endometrial epithelium (next-best unrelated tissue: right atrium; *p* = 0.007, WMW test; **Fig. 3a**). This finding piqued our interest because, from an embryologic standpoint, the endometrium and fallopian tubes both arise from the Müllerian ducts, while the ovaries develop separately from mesodermal epithelium on the urogenital ridges. To further probe this observation we applied our methodology to epigenetic data recently generated for ovarian surface and fallopian tube epithelial cells^20^. Since only a restricted number of ChIP-seq experiments was available for these tissue types, we ran our model using the three histone marks available for 99 tissue types. Using the expanded set of normal tissues, fallopian tube epithelium became the best match to high-grade serous Ovary-AdenoCA, and explained significantly more variance than the second-best match (fetal kidney; *p* = 0.00098; **Fig. 4a**. In addition, we interrogated individual high-grade serous ovarian cancer tumors and found that the majority (95 of 110 samples, 86%) best matched fallopian tube epithelium (variance explained: median, 9%; range, 1-37%). As few as 16 tumors best-matched the epithelium of the ovary (variance explained: median, 1.7%; range, 0.6-23%). However, the difference between the best match and the second-best match was more pronounced in tumors in which the fallopian tube epithelium was the predicted COO (**Fig. 4b**). Our finding supports the unconfirmed hypothesis that secretory epithelial cells of the distal fallopian tube represent the COO of high-grade serous Ovary-AdenoCA^25-27^ and not the epithelium of the ovaries^28^. In contrast to previous work in this area, our approach can associate high-grade serous Ovary-AdenoCA with the epithelium of the fallopian tube without specifying pre-invasive lesions. The true identity of the precursors in the fallopian tube have been the source of a persisting paradox in that pre-invasive lesions occur with high frequency early on in the course of the disease but with low frequency in advanced stages, prompting the concept of ‘precursor escape’ in which tumors do not arise from pathologically suspect pre-invasive lesions, but from normal-looking early serous precursors arising within the fallopian tube that are shed from the fallopian tube early. In light of these observations, our finding, directly pointing to the fallopian tube epithelium, provide additional orthogonal support.

### Gastric mucosa is the best-matched cell type of origin for gastrointestinal tumors that develop through a metaplastic intermediate

Finally, in the 4 remaining cancer types (out of the aggregated 32), the COO matched neither the normal cellular counterpart nor any histologically related cell. In three of these types, the COO supports the idea of the tumor arising from an intermediate step of metaplasia, a process in which mature cells from a given tissue are replaced by mature cells from another, typically adjacent, tissue (**Fig. 3a**). In the case of esophageal adenocarcinoma (Eso-AdenoCA), a cancer of epithelial origin, the best match was stomach mucosa and not the potentially more related esophageal tissue type (*p* = 0.00098) or any other gastrointestinal mucosa (best hit: rectal mucosa; *p* = 0.00098; **Extended Data Fig. 2**). This was observed for the aggregated tumor profiles for Eso-AdenoCA and almost all individual Eso-AdenoCA tumors (88 of 97 tumors; **Fig. 3b**).

Our finding is consistent with the accepted model that Eso-AdenoCA development involves an intermediate step of metaplasia, known as Barrett’s esophagus (BE), in which adult esophageal squamous epithelium changes to columnar epithelium (similar in histological appearance to the lining of the stomach)^29^. Accordingly, the epithelium of BE has been found to exhibit a gastric differentiation program^30,31^. To further test whether the distribution of somatic mutations in BE reflect the COO, we analyzed 23 patient-matched pairs of BE and Eso-AdenoCA^32^. In agreement with previous similar analysis^33^, stomach mucosa was the best match for both BE and Eso-AdenoCA in 21 of the 23 pairs. Typically, the variance explained in the Eso-AdenoCA samples was higher than in the matching BE cases, likely due to the increased number of mutations.

However, we saw a significant correlation between the variance explained in both samples of the pairs (r = 0.44, *p* = 0.037, Pearson correlation coefficient; **Extended Data Fig. 7**). Together, these findings highlight a close relationship between BE and Eso-AdenoCA and suggest that Eso-AdenoCA may acquire most of its mutations during the metaplastic state of BE.

We identified 2 additional gastrointestinal tumor types (Panc–AdenoCA and biliary adenocarcinoma [Biliary–AdenoCA]) in which the best-matched cell type was stomach mucosa suggesting a metaplastic intermediate step. In Panc-AdenoCA, the variance explained was significantly higher in models of stomach mucosa than pancreatic tissue (enriched for acinar cells; *p* = 0.00098, WMW test) and to the second-best gastrointestinal tissue (duodenum mucosa; *p* = 0.00098). These data support the current thinking that ‘acinar-to-ductal’ metaplasia (ADM) is the initial morphologic change towards Panc-AdenoCA and that this loss of acinar differentiation induces stomach-specific gene expression^34,35^. Similar results were found in Biliary-AdenoCA, where the variance explained was significantly higher in models of stomach mucosa than the second-best gastrointestinal tissue (duodenum mucosa; *p* = 0.00098).

We are left with one tumor type, bladder transitional cell carcinoma (Bladder-TCC), whose best match was also stomach mucosa. Our chromatin data set did not include all potentially relevant normal cell types across the 4 cancer types (i.e., enriched cell subtypes such as ductal pancreatic cells and esophageal mucosa as well as the direct normal cell counterparts cholangiocytes and bladder transitional cells). Despite these limitations, we observed a significant difference between stomach mucosa and the second-best tissue for all gastrointestinal tumor types but not Bladder-TCC (second best: rectal mucosa; *p* = 0.54; **Extended Data Fig. 8**). Likewise, a considerable fraction of individual gastrointestinal tumors best matched stomach mucosa – Eso-AdenoCA (88 out of 97, 91%) and Panc-AdenoCA (165 out of 232, 71%) – while only 1 out of 23 individual Bladder-TCC samples (4.3%) best matched stomach mucosa. These observations suggest that the gastrointestinal cancer types are different from Bladder-TCC in that they may have undergone metaplasia. In the case of Bladder-TCC however we may simply be lacking the chromatin data from the most appropriate normal tissue type.

Collectively, our results reveal that most mutations across the broad landscape of different tumor types occur during a time when the chromatin organization of the cells is still similar to the COO. Of note, we found that using an alternative regression method leads to quantitatively and qualitatively similar results (see **Extended Data Information** for details; **Extended Data Fig. 9a,b; Extended Data Table 1**).

### Chromatin differences between cell differentiation states in a single tissue help to decipher the precise cell subtype of origin

With the COO identified for almost all (31/32) aggregated cancer types, we next tested whether our method was powerful enough to delineate the COO on a finer scale by matching the COO to different tumor subtypes of the same lineage. Here, we focused on brain, blood, breast and prostate cancers.

Within the brain, different cell types, including neuronal and glia cells, have been identified at different differentiation stages. Our analysis nominated neurospheres, but not mature brain regions, as the best match across all gliomas (glioblastoma, CNS-GBM; oligodendroglioma, CNS-Oligo; pilocytic astrocytoma, CNS-PiloAstro). In contrast, medulloblastoma (CNS-Medullo) best matched the germinal matrix, a fetal brain region that gives rise to neuronal cells but not neurospheres (*p* = 0.0046; **Extended Data Fig. 10**). Neurospheres currently represent the best model of neural stem cells (NSCs)^36^ and our findings are in line with the current hypothesis that undifferentiated NSCs and more committed progenitor cells give rise to all above-mentioned gliomas^37-40^. Of note, our findings also recapitulate the historical understanding of the distinct COOs for gliomas and CNS-Medullo wherein gliomas were thought to derive from glia cells, while CNS-Medullo was believed to derive from neuronal-type cells^41,42^. Further support for our findings comes from single-cell transcriptomics of the developing mouse brain showing that CNS-Medullo tumors mirror primitive cells in the neuronal lineage that are found in the germinal matrix but are all-but-gone at birth^43^.

Similar to the nervous system, the differentiation cascade of the hematopoietic system is particularly well-understood, giving rise to distinct cancer subtypes. Accordingly, we next searched for the COO of different hematopoietic neoplasms, including tumors originating from B cells (chronic lymphocytic leukemia, CLL; B cell Non-Hodgkin’s Lymphoma, BNHL) and myeloid cells (acute myeloid leukemia, AML; myeloproliferative neoplasm, MPN). Aggregated tumor profiles of myeloid tumors best matched CD34^+^ hematopoietic stem cells (HSCs) while B cell tumors best matched CD19^+^ B cells (compared to next-best non-hematopoietic tissue: AML, *p* = 0.006; MPN, *p* = 0.001; CLL, *p* = 0.00098; BHNL, *p* = 0.001; **Extended Data Fig. 11a**). Consistently, most individual samples best matched the same cell type as the aggregated profiles (**Extended Data Fig. 11b**). This is in accordance with the understanding that B cell tumors develop late during B cell differentiation (i.e., in cells that already express CD19^44^). In contrast myeloid leukemic transformation primarily occurs in multipotent or granulocyte macrophage progenitors, both of which express CD34, the marker for HSCs^45^.

Breast cancer also has several different subtypes that arise from within the same tissue^46^. Four main ‘intrinsic’ molecular subtypes have been identified by gene expression profiling: luminal A, luminal B, HER2-enriched and basal-like^47^. Epithelial cell types from the duct-lobular unit of the mammary gland include luminal and basal cells, which both derive from mammary stem cells (MaSCs) via different progressively committed progenitors^48^. We used chromatin profiles generated from four subsets of normal breast cells, isolated via two cell-surface markers: (i) one subset that includes MaSCs, basal progenitors and basal mature cells (CD49f^+^ EPCAM^low/-^); (ii) luminal progenitors (CD49f^+^ EPCAM^high^); (iii) luminal mature cells (CD49f^−^ EPCAM^high^); and (iv) stromal cells (CD49f^−^ EPCAM^−^)^49^. Aggregated tumor profiles of Breast-AdenoCA best matched mature luminal cells (**Fig. 3a**). Similarly, the analysis of pre-invasive disease found mature luminal cells to be the best match of DCIS (ductal carcinoma in situ; 10% variance explained), further confirming our findings (data not shown; n = 3). Principal component analysis (PCA) of the mutation profiles significantly separated basal-like and non-basal subtypes into two distinct clusters, suggesting that the origin of each of these clusters is different (*p* = 1×10^−67^, Fisher's exact test; **Extended Data Fig. 12a**). Accordingly, we discovered that mutation profiles of basal-like breast tumors matched best to breast luminal progenitors (all *p*-values < 0.003, WMW test) while luminal A, luminal B and HER2-enriched subtypes matched best to mature luminal cells (all *p*-values < 0. 001; **Fig. 5a**). We confirmed our results using an additional independent data set of 274 breast cancer whole-genome sequences (**Fig. 5b**)^50^. Our finding held true even when we divided patients according to subtype and ancestry, albeit with lower variance explained in the smaller sub-cohorts compared to the aggregated breast cancer analysis (**Extended Data Fig. 12b,c**). African-American females have been found to exhibit a higher prevalence of basal-like breast tumors^51^. This higher susceptibility to basal-like subtypes has been suggested to be influenced by a higher basal to luminal cell ratio in African-American women^52^. In contrast to this hypothesis, our results suggest that basal-like breast tumors originate from luminal progenitors independent of ethnicity.

Next, we asked whether our COO results of breast cancer depend on specific genetic events. We therefore aggregated tumors according to their type of homologous recombination deficiency (HRD; see **Extended Data Information**). All 34 HRD-associated basal-like breast cancers best matched luminal progenitors, regardless of *BRCA1*, *BRCA2* or *RAD51C*^53^ inactivation (**Extended Data Fig. 13**). Likewise all 26 HRD-associated luminal A/B and HER-2 enriched subtypes best matched mature luminal cells, again irrespective of the inactivated HR gene. We found similar results when analyzing individual tumors: most basal-like tumors (19/34, 55.9%) best matched luminal progenitors while the majority of LumA/B and HER-2 enriched tumors (15/24, 62.5%) best-matched luminal mature cells (**Fig. 5c**). Although different breast cancer subtypes are associated with different HRD mechanisms, we found that the HRD event was not influenced by the COO.

In addition to our analysis of ductal carcinoma, we also sought to understand the origins of lobular cancer (Breast-LobularCA). Both entities are assumed to derive from the mammary duct-lobular unit but are distinct in terms of their molecular and histological features^54^. Until now no conclusive reports on the COO of Breast-LobularCA have been published. We found that Breast-LobularCA matched best to mature luminal cells (**Extended Data Fig. 12d**), a finding that concurs with the observation that up to 72% of lobular tumors are classified as the molecular subtypes luminal A/B^55^.

Like the breast, the prostate is a hormone-regulated, two-layered glandular organ comprising luminal and basal cells. We differentiated between these two main cell types using ChIP-seq data from 3D organoid luminal and basal cell cultures, isolated via two cell surface markers from normal tissue: CD26 to enrich for luminal cells, and CD49f to enrich for basal cells^21,56^. Only two histone modifications were evaluated for the organoids, and we ran our model on 13 tissue types for which those marks were available. The best match for prostate adenocarcinoma (Prost-AdenoCA) was prostate bulk representing a primary tissue. Luminal cells explained less variance than bulk primary tissue but not significantly lower. In contrast, basal cells explained significantly less variance (*p*-value = 0.02, WMW test; **Extended Data Fig. 14a**) suggesting that luminal cells are the COO of Prost-AdenoCA (**Extended Data Fig. 14b**). Our finding of the lack of a significant difference also points out that organoid-derived cells maintain a chromatin structure highly comparable to primary tissue which is not the case for cell lines^57^. Furthermore, we used epigenetic data from two additional prostate cancer organoids^58^. One was derived from an androgen receptor (AR)-independent basal/mesenchymal prostate cancer phenotype (PCa1) and one from an AR-dependent luminal phenotype (PCa2). However, we found that they both explained less variance than normal prostate cells. This suggests that the chromatin structure evolves significantly during tumorigenesis and that most of the somatic mutations occurred before this change.

Overall, our findings in cancer subtypes demonstrate the power of our method for inferring COOs from distinct differentiation states along the same cell hierarchy.

### The cell-of-origin captures important features of cancer biology

To further support our findings of the COO and to highlight the biological relevance to tumorigenesis we performed three types of orthogonal analyses. First, we investigated GWAS hits in estrogen receptor negative (ERneg) and positive (ERpos) breast cancer subtypes to estimate the influence of the cellular context on the effect size of single-nucleotide polymorphisms (SNPs) to heritability. Second, we used metastatic samples to determine whether they still preserve molecular echoes of the COO. Lastly, we evaluated driver mutations to study whether the COO can explain differences in the oncogene and tumor suppressor composition across cancer types.

### Germline risk alleles for breast cancer reside in active chromatin regions of the cell-of-origin

Genome-wide association studies (GWAS) for breast cancer risk have identified 107 independent loci^59^. Since we have epigenetic data for different breast cell subtypes (from progenitors to mature cells), we wanted to assess additional evidence from risk GWAS for the best matching cell type of origin. To this end, we explored whether GWAS heritability for ERneg and ERpos breast cancer were enriched in active enhancer/transcription regions (peaks of H3K27ac) of the different cell types. H3K27ac peaks were selected for two reasons: (i) they are generally enriched across complex disease^60^; and (ii) highly enriched with GWAS hits for prostate cancer^61^. Indeed, we found the largest enrichment of ERneg breast cancer heritability for luminal progenitors followed by luminal mature cells (9.8× and 9.5×; all *p*-values < 0.02, Z-test; **Extended Data Fig. 15** and **Extended Data Table 3**). In ERpos breast cancer, luminal breast cell types also showed a high enrichment of heritability (progenitor, 13.3×; mature cell, 12.6×; all *p*-values < 0.0005), though not the highest of all cell types tested. These differences may reflect the fact that high-penetrance mutations are associated with ERneg breast cancer while most of the susceptibility loci show a stronger association with ERpos disease^62^. Of note, in both ERneg and ERpos tumors, the basal and the stromal breast cell types explained considerably less heritability than the luminal cell types (**Extended Data Fig. 15**). Previous analysis of GWAS heritability based on H3K4me1 regions of a different breast cell type (myoepithelium) achieved a poorer enrichment of 6.7-fold compared to ~13-fold using our suggested COO^63^. Overall, our findings indicate that active enhancer/transcription regions of the COO are highly enriched with GWAS heritability, thereby highlighting the tissue specificity of those genes and the significance of the cellular context. Such effects could help in localizing GWAS associations to the most likely cell type-specific regulatory features.

### Metastatic samples allow the identification of their originating tissue types

With the exception of Skin-Melanoma samples, which are largely metastatic in origin, almost all of the tumors in the PCAWG data set are primaries (**Extended Data Table 1**). To extend our method further we profiled 2,044 metastatic samples (HMF metastases)^64^ from 22 of the tumor types available as primaries in the PCAWG data set (**Extended Data Table 4**).

Fascinatingly, we found that the power of our method for inferring COOs is similar across primary and metastatic disease (**Fig. 6a**). Analogous to our analysis of primaries the same COO was detected in almost all types of metastatic disease. In only two endocrine tumor types (Panc-Endocrine and Thy-AdenoCa) did unrelated cell types show the highest variance explained. This is most likely due to a smaller sample size leading to a reduced power to detect in cancer types with an already low mutation frequency. Similar to our findings in primaries, fallopian tube epithelium showed the highest prediction accuracy in high-grade serous Ovary-AdenoCA giving additional evidence that the fallopian tube serves as the COO (next best: colonic mucosa; *p* = 0.00098, WMW test; **Extended Fig. 16a**). Metastatic breast cancer subtypes were classified according to their expression of hormone receptors. Analogous to our results when using Prediction Analysis of Microarray 50 (PAM50)-grouped primaries, we observed that triple negative breast cancer best matched luminal progenitors (all *p*-values *p* < 0.02; **Extended Fig. 16b**), while all other subtypes best matched luminal mature cells (all *p*-values < 0.03). Overall, the analysis of metastases provided independent validation for our findings in primaries. Although the epigenetic landscape changes after oncogenic transformation^65^ our data suggest that new mutational events and processes that may evolve during late tumorigenesis do not override the original information reflecting the normal tissue (**Fig. 6b**).

### The cellular context of driver genes highlights tissue specificity

In our main analysis, we found that the chromatin structure of the COO is highly associated with the acquisition of somatic mutations. In parallel to the overall mutational landscape, alterations in cancer genes have been found to be tissue specific. While some driver genes are altered across tumor types, most are mutated in only a restricted set of tumor types. Alterations in driver genes are expected to be positively selected in cells in which they are transcribed and hence have open chromatin. We therefore hypothesized that the COO chromatin landscape offers a cell type-specific fertile ground at sites of high transcriptional activity for the acquisition of cancer type-specific driver mutations.

We analyzed 78 individual genes that were mutated in >2% of patients and located in the 1Mb windows not removed due to mappability (median 8%, range 2-86%) and identified either as significantly mutated in PCAWG^66^ or previously shown to be drivers^67^. These 78 genes were mutated in 24 tumor types, with most of them (42 genes) only in one tumor type. Across all tumor types, we observed 225 alterations in these genes (**Extended Data Fig. 17**).

Next, we inferred the regional transcriptional activity in a 1Mb window around the driver genes by counting ChIP-seq reads of three activating chromatin marks (H3K4me1, H3K4me3 and H3K36me3) for each tissue type. For each chromatin mark, we then compared the number of reads in the COO to the other tissue types and determined whether it is an outlier (see **Methods**, **Extended Data Information**). Altogether, we found 105 regions with outlier activity in the COO associated with each of the tumor types.

In the next step, we searched for co-occurrence of tumor type-specific driver gene and COO outlier activity. Supporting our hypothesis, we observed a significant enrichment of driver genes in chromatin regions with outlier activity in their COO: 29 pairs affecting 23 genes (*p* = 6.9×10^−6^, Fisher's exact test; **Extended Data Fig. 17**). Even when controlling for the total number of outliers in each tissue and each gene, we found that the overlap between outliers and drivers was significantly larger than expected by chance (29 vs. an average of 19, *p* = 0.0004, permutation test).

To further understand this observation, we gathered functional and biological annotations from the literature for 18 of the 23 (78%) drivers and found 4 of them to contribute to tissue homeostatic processes and 14 to be crucial for development and differentiation of their corresponding COO (see **Extended Data Information** for details). This suggests that a subset of the drivers not only confers a selective growth advantage in their mutant form but also have major cell type-specific regulatory functions in the COO.

In addition, we observed that the COOs of B cell-derived tumors (Lymph-BNHL, Lymph-CLL) and Breast-AdenoCA showed an exceptionally high number of regions with outlier chromatin activity (74 of the 105 outlier regions in these three tumor types vs. 31 in the others). This significant enrichment of outlier activity (*p* = 2.2×10^−16^, Fisher's exact test) paralleled a significant enrichment of driver genes in these outlier regions (11 of the 29 pairs in these three vs. 18 in others; *p* = 0.0004). B cells and mammary cells have been reported to have a high turnover rate^68,69^, suggesting that cell proliferation of the COO could influence the risk of acquiring driver mutations. Two previous studies support this hypothesis: (i) mathematical modeling correlated the number of cell divisions with the overall cancer risk^70^; and (ii) the frequency of actively cycling normal breast cells was reported to be associated with higher breast cancer risk^71^.

Overall, our data suggest that COO-specific chromatin sites of high transcriptional activity are associated with mutations in driver genes in a cell context-dependent manner. Despite an overall lower abundance of somatic mutations in these regions, we assume the enrichment of drivers due to the positive selection of genes that play an important functional and physiological role. By differentiating subclasses of driver genes, our approach helps to better understand the potential contribution of the COO to tumor initiation. In addition, our approach of focusing on active regions in the COO could guide the search for new tumor-specific drivers, including non-coding genes in enhancer/promoter regions.

## DISCUSSION

Understanding the dynamics of tumor formation by studying the genomics of advanced tumors at diagnosis is a challenge due to the acquired plasticity that typically masks features of the originating tissue. However, we found footprints of early tumorigenesis in the COO that are preserved in the tumor genome. We took advantage of the correlation between chromatin marks and mutational profiles, which reflect transcriptional activity. Typically, decreased mutation frequencies are observed in open, transcriptionally active chromatin, likely due to more efficient DNA repair or fewer errors in earlier replication regions^72,73^.

One key finding in this study is that the mutational landscape is significantly influenced by the normal cellular context of the COO. This observation is in accordance with previous results in single neurons showing that the density of somatic mutations varies according to the originating brain region^74^. Our data imply that most somatic mutations detected in tumors arise at a time when the chromatin state still resembles the normal cell (either in normal cells or early in tumor progression). This observation is supported by a previous study estimating that at least half of a tumor’s mutations occurred before the onset of neoplasia^75^. The concept that most mutations happen before tumor initiation is further reinforced by the higher prediction accuracy of clonal compared to sub-clonal mutations. Since the epigenetic landscape of tumors is distinct from that of their originating normal cells^65,76,77^ mutations that were acquired earlier can be better explained by the chromatin structure of the COO. This is not the case for copy number changes, as these typically occur synchronously with oncogenic transformation.

Our findings help to better understand the biology of cancer. The identification of the fallopian tube epithelium as the COO for almost all high-grade serous Ovary-AdenoCA tumors without the use of precursor lesions serves as additional evidence in the debate regarding the origin of ovarian cancer. Generally, by matching the cancer genome directly to normal chromatin profiles, we bypassed the necessity of pre-invasive disease for identifying cancer origins. Moreover, in cases in which we analyzed mutational profiles in precursor lesions (DCIS and BE) we found that they matched the same COO as their invasive counterparts. While it is difficult to causally connect pre-invasive lesions to the later appearing invasive cancer, our findings point towards a common origin and thereby support a multistep model of cancer development. Our result that Prost-AdenoCA matches the chromatin state of organoids derived from normal cells equally well as primary prostate tissue suggests that these model systems provide good approximations of the COO and might accurately reflect the characteristics of cells *in vivo*.

Another novel finding from the present work is the discovery that stomach-like mucosa serves as the preferred representative proxy for different metaplasia phenotypes in the gastrointestinal tract. Our findings are particularly important to understand the debated origin of Panc-AdenoCA^78^. The pancreas consists of three functionally distinct, but anatomically interwoven cell populations: islet or endocrine cells (~2.4% of pancreatic area), exocrine acinar cells (~86%), and exocrine ductal cells (~1.1%)^79^. Although debated, the current hypothesis is that Panc-AdenoCA develops from acinar cells through a sequence of acinar-to-ductal metaplasia (ADM) followed by pancreatic intraepithelial neoplasia (PanIN)^34^. There is a large body of evidence that both metaplastic cell types acquire gastric cell features^35,80,81^. Additional support comes from the embryonic development of the gastrointestinal tract. The pancreas forms because Hedgehog signaling is suppressed; when not suppressed, Hedgehog drives the endoderm to intestine transition. Hedgehog signaling is upregulated in Panc-AdenoCA and points towards a close relationship between gastric epithelium and the COO of Panc-AdenoCA^82^. Our results are consistent with these observations and indicate that most mutations in Panc-AdenoCA occur in a cell state that resembles stomach mucosa (most likely a metaplastic state) and not in acinar cells (represented by native pancreatic tissue, and only the 10^th^ best match).

One main conclusion from this study is that cancers from the same organ but of distinct subtypes can be matched to different COOs. In particular, we study the COO of breast cancer subtypes. There are three hypotheses about the origins of breast cancer: (i) the MaSC is the COO of all subtypes^83^; (ii) basal stem/progenitor cells give rise to basal-like cancers and luminal progenitors to luminal tumors^84^; and (iii) all breast cancer subtypes derive from cell types along the luminal differentiation hierarchy^83^. Since previous conclusions were drawn from murine experiments, our analysis of human tumors yields valuable insights given the morphogenetic differences between the mouse and human mammary glands^85^. Here, using our method of integrating chromatin state with mutational density, we found data that support the hypothesis that basal-like tumors originate from luminal progenitors and all other subtypes from mature luminal cells, a finding consistent with previous gene expression profiles^86^. We also found that the association of breast cancer subtypes with their corresponding COOs is not influenced by the specific HR-inactivation event. This result is supported by previous data showing that functionally disabling *BRCA1* in luminal progenitors, but not in mature luminal or basal cells, gives rise to basal-like breast cancer^87-89^. Together, our data are in accordance with the concept that the COO helps in dictating the cancer type that eventually occurs even in the presence of the same genetic event.

While almost all aggregated tumor mutation profiles matched their direct cellular counterpart or a close proxy, many individual tumors (depending on the cancer type) did not have enough mutations to distinguish the COO from other cell types. In some tumor types, such as blood cancers, the chromatin profiles of the different COOs are highly similar to each other; hence, a larger number of mutations is required to distinguish between individual cancers from the same lineage. Furthermore, chromatin modifications were primarily derived from bulk normal tissues that are heterogeneous. Chromatin marks were available for enriched cell populations only in a few cases (e.g., breast and prostate cell subtypes). Despite the heterogeneity of bulk normal tissue that can mask the features of the COO, most cancer types matched their expected normal cell counterpart. This suggests that an enrichment of the COO in the normal tissues is sufficient for being identified by our approach. One limitation of our study is that the current data do not provide a 1:1 relationship between all cancer types and normal cell types. In cases in which we did not have chromatin data of the direct normal cellular counterpart, an appropriate COO could often only be reached as a proxy (e.g., histologically related cells). Accordingly, analyses such as these will benefit from additional cell type-specific chromatin data, especially when derived from single cells and/or enriched cell populations.

Our results may have unique clinical implications. We show, for the first time, that most somatic mutations in metastases reflect the chromatin state of the COO. This finding can help elucidate the origins of a metastatic lesion, an approach that can be applied to identify the COO of cancers of unknown primary (CUPs). A precise characterization of the tumor COO is particularly relevant given recent ‘basket’ trial data, suggesting that the tumor’s response to treatment depends not only on the oncogenic mutations but also on the specific cell context of the tumor and/or the COO^90^. Thus, our findings may supplement clinical decision-making and improve trial design.

Collectively, our results will help (i) inform the development of animal and cellular models to understand tumor initiation and progression in greater detail; (ii) enable the evaluation and development of treatment options that take into account the COO; and (iii) focus on the relevant cell type for early detection or prevention of cancer.

## MATERIALS and METHODS

### Genomic data

As described previously^10^, we divided the human genome (hg19) into 1Mb windows. We excluded regions overlapping centromeres and telomeres as well as regions with a low fraction of uniquely mappable bases (<92% of bases within uniquely mapped 36-mers). This approach resulted in 2,128 1Mb windows, corresponding to ~2.1Gb of DNA.

We obtained whole-genome mutation data for 2,550 cancer genomes, belonging to 32 different tumor types, from PCAWG. Clinical annotations of tumor samples, the generation of sequencing data, and their analysis through a series of pipelines have been described in detail^16^. The **Extended Data Table 1** provides an overview of tumor samples characteristics relevant for our approach.

We included all cancer types with ≥ 10 individual samples; we excluded all non-malignant bone tumors and the category ‘epithelioid bone neoplasms’ that summarizes three cancer types of distinct origins. For each individual cancer sample we calculated the total number of mutations in each window. Aggregated tumor profiles for each tumor type were calculated by summing the total number of mutations across all tumors from that tumor type in each window. Mutation clonality annotation was obtained from PCAWG^22^.

In addition, we obtained whole-genome mutation data from a metastatic data set based on the Hartwig Medical Foundation cohort (HMF metastases) ^64^. Altogether we analyzed 2,044 metastatic cancer genomes, belonging to 22 tumor types (**Extended Data Table 4**). We concentrated our analysis on tumor types present in the PCAWG data set and included all cancer types with ≥ 6 individual samples. In cases with multiple samples per patient only one tumor (‘A’) was used. Samples lacking mutation counts or appropriate histological annotations were excluded. We included a limited number of primary tumors that were not previously surgically removed when the sampling of the metastatic lesion was not feasible or safe (see **Extended Data Table 4** for details).

### Chromatin data

We downloaded read alignment information from eight ChIP-Seq experiments against six active^91^ histone modifications (H3K27ac, histone H3 lysine 27 acetylation; H3K27me3, histone H3 lysine 27 trimethylation; H3K36me3, histone H3 lysine 36 trimethylation; H3K4me1, histone H3 lysine 4 monomethylation; H3K4me3, histone H3 lysine 4 trimethylation; H3K9ac, histone H3 lysine 9 acetylation) and one repressive histone modification (H3K9me3, histone H3 lysine 9 trimethylation) as well as the background sample ‘Input’ (**Extended Data Table 2**). We obtained data (human primary cell cultures, enriched cells and bulk tissues) for 87 cell types from the Roadmap Epigenomics Consortium (release 9)^17^, for kidney, thyroid and prostate from ENCODE^18^ and for four breast cell subtypes as well as four endometrial cell types from IHEC^19^ (**Extended Data Table 2**). In addition, four chromatin profiles for fallopian tube and ovarian surface epithelium as well as four profiles for prostate organoids were gathered from publications^20,21^. Similar to the mutation data, for each of the ChIP-Seq data sets, we calculated its profile, i.e. the number of reads in the same 1Mb windows as defined above.

Chromatin modifications were grouped when they were biological replicates or derived from histologically related tissues to evaluate the best match of individual tumors (**Extended Data Fig. 1**). For the following cell types biological replicates were available and we considered them as one: fibroblast primary cells, melanocyte primary cells, keratinocytes primary cells, T helper memory cells, T helper naive cells, rectal mucosa, and endometrial stroma. Likewise, 10 groups were formed from the following histologically related cells: (i) the ‘brain group’ consists of cells from fetal brain, adult brain regions and neurospheres; (ii) the ‘bone/soft tissue group’ consists of all mesenchymal cells throughout the body including muscle and other connective tissues; (iii) the ‘squamous group’ consists of squamous esophagus and skin keratinocytes; (iv) the ‘endometrial epithelium group’ consists of EndoFollicular and EndoSecretory cells; (v) the ‘breast luminal group’ consists of luminal progenitors and mature luminal cells; (vi) the ‘breast other group’ consists of basal, breast myoepithelial, and breast variant human mammary epithelial cells; (vii) the ‘T cell group’ consists of all T cells; (viii) the ‘B cell group’ consists of all B cells; (ix) the ‘HSC cell group’ consists of all HSCs; (x) the ‘blood cell group other’ consists of myeloid cells. In addition, we combined fetal and adult tissues (lung, thymus and kidney) as well as gastric and stomach mucosa profiles to represent stomach mucosa.

### Random Forest regression analysis

For each of the cancers and the aggregated tumor profiles, we used Random Forest regression (with 1,000 trees) to predict the mutation density profile using the chromatin profiles. For each mutation density profile, we trained 98 separate regression models using the complete chromatin profiles associated with each of the tissue types. For the cancer-specific aggregated mutation profiles, we calculated the performance using 10-fold cross validation (i.e., we divided the 2,128 windows into 10 non-overlapping sets, trained the model on 9 sets and predicted the number of mutations in the remaining windows). We additionally identified the best-matching model for each individual cancer by training 98 regression models, using all windows, and finding the one with highest prediction accuracy. For each cancer type, we then calculated the proportion of individual tumor samples in which the best-matching model was either the same as the presumed COO of the aggregated tumor profile or, in some tumor types, belonged to a group of histologically related tissue types. We reported the overall prediction accuracy by calculating the average R^2^ between the predicted and observed profiles across the 10 sets of windows. The analysis was run using the *caret* and *ranger* packages in ‘R’. To demonstrate the robustness of Random Forest regression, we here used different seeds for random numbers to control for the noise introduced into the model. Using this approach we did not find qualitatively significant differences in the best matches (data not shown).

To compare models based on two different chromatin profiles, we used the paired Wilcoxon-Mann-Whitney (WMW) test between the R^2^ values from ten-fold cross validation tests. Correlations were analyzed using the Pearson correlation coefficient. Mutation frequency is given as overall mutation frequency and was determined by summing up all mutations of all samples of a given tumor type in 1Mb windows.

We compared the prediction accuracy for clonal and sub-clonal mutations across cancer types. To assure that differences in prediction accuracy in our models are not driven by a higher number of clonal mutations, we simulated clonal mutation profiles. Monte Carlo simulation was used to generate profiles with a total number of clonal mutations that equals the total number of sub-clonal mutations in each cancer type. In detail, simulated profiles were generated by a Poisson number generator using rates that are equal to the normalized profile of clonal mutations multiplied by the total number of sub-clonal mutations. In addition, the tissue that represents the median across all tissues and the Median Absolute Deviation (MAD) were determined. For the comparison of models trained on histone marks from different cell types, the robust ẑ-score was computed^92^.

### Analysis of the chromatin environment of candidate driver genes

Lists of significantly recurrently mutated coding driver genes and previously described driver genes were provided by the PCAWG Drivers and Functional Interpretation Group^66,67^. Of those, 78 drivers were used in our analysis, which are located in the 1Mb windows that were not removed due to mappability; they resided in 24 tumor types.

To analyze the chromatin context of different driver genes, we used, for each driver, the number of reads of H3K4me1, H3K4me3 and H3K36me3 in the 1Mb window in which the gene is located across the core 98 tissue types. For cell types with biological replicates and groups of histologically related cells (see above), the number of ChIP-Seq reads was calculated as the median number of reads over all replicates. Square-root transformation of data was used. We then compared the number of reads in the window for the best-matched tissue (i.e., the identified COO) to the other 97 tissues and considered the gene to have a tissue-specific chromatin environment if it was an outlier (i.e., above the 1.5*interquartile range from the 75^th^ percentile of the 97 tissues). In addition, we used the top 1%, top 2.5% and top 5% to define outlier status. By comparing the same 1Mb window across tissue types, we accounted for possible variations in the number of binding sites or ChIP affinities^93^. We performed a permutation test with 1,000,000 permutations using the ‘Curveball algorithm’^94^.

### Clustering of individual breast cancer genomes

We performed principal component analysis (PCA) on the mutation profiles of all individual breast cancer genomes from PCAWG and Nik-Zainal et al.^50^. Each tumor was classified to one of the breast cancer subtypes using RNA expression levels of 50 genes (PAM50, Prediction Analysis of Microarray 50) taken from the corresponding literature^50,54^. We used *k*-means clustering (with *k*=2 and 1-Peason correlation as the distance metric) to cluster the tumors based on their first two principal coordinates from mutation counts along 1Mb windows.

### Analysis of GWAS heritability

We used stratified linkage disequilibrium score regression (S-LDSC)^60^ to quantify the enrichment of GWAS heritability in epigenetically active regions (annotations). Briefly, S-LDSC evaluates the full distribution of GWAS associations (not restricted to significant SNPs) and infers heritability parameters from the relationship between the effect-size and the LD of each SNP. Annotations that are in LD with higher effect-size SNPs will be assigned higher heritability and the converse for low effect-size SNPs. Further details are discussed elsewhere^60,61^.

GWAS summary statistics were downloaded from recent studies of ER negative (N=127,442) and ER positive (N=175,475) breast cancer risk^59^. Each study was restricted to ~1M HapMap3 SNPs that are typically well-imputed across all GWAS platforms and have been shown to perform well in heritability analyses. We then included each epigenetic annotation in turn in the S-LDSC model together with the standard “baseline model” that captures potential confounding factors (generic features such as coding, promoter and intronic). Enrichment for each annotation was computed as the % of heritability accounted for by the annotation, divided by the % of SNPs contained in the annotation, wherein an enrichment of 1.0 is expected under the null. Statistical significance was assessed by the block jackknife as implemented in S-LDSC. For context, we evaluated H3K27ac ChIP-seq calls from ROADMAP^17^ and breast^49^ cell types using the imputed, narrow peaks call-set. To test statistical significance, *p*-values were generated from Z-scores.

## ACKNOWLEDGEMENTS

P.P. and K.K. were partly funded by the startup funds of G.G. at Massachusetts General Hospital. K.K. and G.G. are supported by a CDMRP award (W81XWH-17-1-0084). G.G. is funded by the Broad/IBM Cancer Resistance Research Project, by National Institutes of Health grants R01DE022087 and P01CA206978-01 as well as the Paul C. Zamecnik Chair in Oncology at the Massachusetts General Hospital Cancer Center. P.P was supported by the Schneider-Lesser fellowship and the V-Scholar award by the V-Foundation. W.D.F. is funded by the Canadian Institute for Health Research (FDN-148390). R.K., M.K. and K.V. were supported by the European Structural and Investment Funds grant for the Croatian National Centre of Research Excellence in Personalized Healthcare (contract #KK.01.1.1.01.0010), Croatian National Centre of Research Excellence for Data Science and Advanced Cooperative Systems (contract KK.01.1.1.01.0009), the European Commission Seventh Framework Program (Integra-Life; grant 315997) and Croatian Science Foundation (grant IP-2014-09-6400). H.-G.K. and K.H. were funded by the Basic Science Research Program through the National Research Foundation of Korea (NRF) under the Ministry of Science, ICT and future Planning (NRF-2017R1A2B2008729). I.R. is supported by the FWO Odysseus programme (Research Foundation Flanders). A.G. is supported by the Claudia Adams Barr Award. R.B. is supported by a Cancer Research UK grant (A13086) and the Ovarian Cancer Action. C.L.S. is an investigator of the HHMI and is supported by National Institutes of Health grants CA155169, CA193837, CA224079, CA092629 and CA16000. W.R.K. is supported by the Dutch Cancer Foundation (KWF BUIT-2015– 7545) and a Prostate Cancer Foundation Young Investigator Award (17YOUN10). L.D.S. and W.J. were supported by funding from the Province of Ontario. We thank Els Witteveen, Haiko Bloemendal, Henk Verheul and Laurens Beerepoot for their efforts to collect material and data for the set of HMF metastases. We thank Shawn M. Gillespie for his support in generating the ChIP-seq data from prostate organoids. We are grateful to Esther Rheinbay and the PCAWG steering committee for helpful feedback on the manuscript. We thank Brett S. Carver for helping in accessing the prostate organoid data set. This publication and the underlying study have been made possible partly on the basis of the data that Hartwig Medical Foundation and the Center of Personalized Cancer Treatment (CPCT) have made available to the study.

## AUTHOR CONTRIBUTIONS

This work was carried out a subgroup of the PCAWG Pathology and Clinical Correlates Working Group based on data from the ICGC/TCGA Pan-Cancer Analysis of Whole Genomes Network. K.K. and R.K. conceived the study, performed analyses and wrote the manuscript. N.J.H., E.Cur. and M.K. performed analyses. J.K. contributed analysis tools and advice. C.L.S., B.E.B., N.S. and C.H. provided data. E.Cup. provided data and performed analyses. K.H., H.-G.K., A.G., P.F.A. and H.L. performed analyses and helped edit the manuscript. S.G., M.P.L., W.R.K. and R.B. provided data, scientific insight and/or helped edit the manuscript. W.J., K.W.M., L.Z.B., O.E., A.V.B., I.R., C.D.N., M.M.-K., L.W.E., C.T., K.V., M.M., E.J. and K.A.H. provided scientific insight and/or helped edit the manuscript. L.D.S. helped edit the manuscript and shape the discussion section. D.N.L and A.K. wrote, provided scientific insight and/or helped edit the manuscript. P.P. conceived the work, performed and oversaw the analyses, and wrote the manuscript. W.D.F. and G.G. conceived the work, oversaw the analyses, and wrote the manuscript.

