## Extended Data: Information, Figures and Legends for "Tumor mutational landscape is a record of the pre-malignant state"

### Extended Data Figure 1

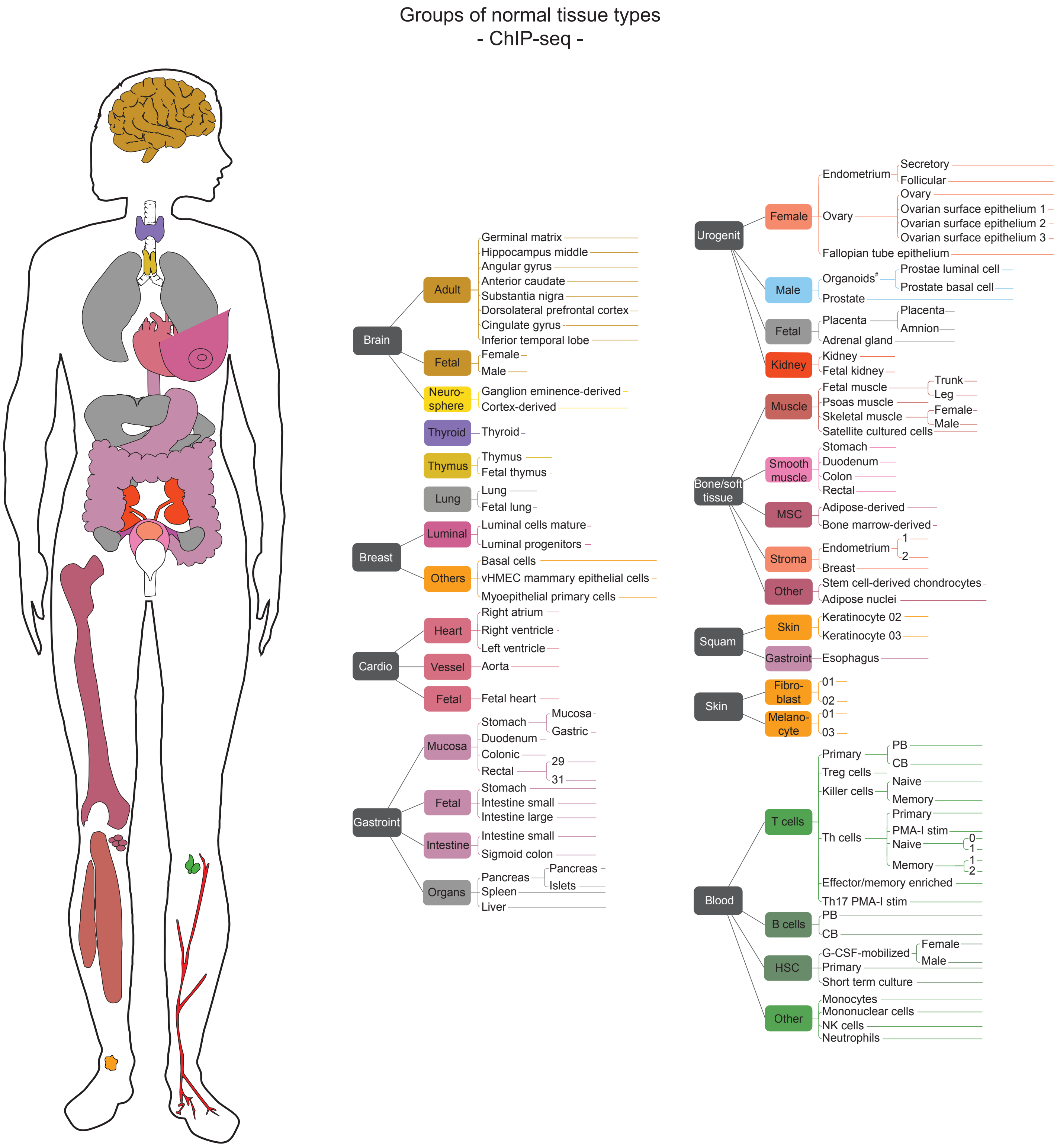

### Extended Data Figure 2

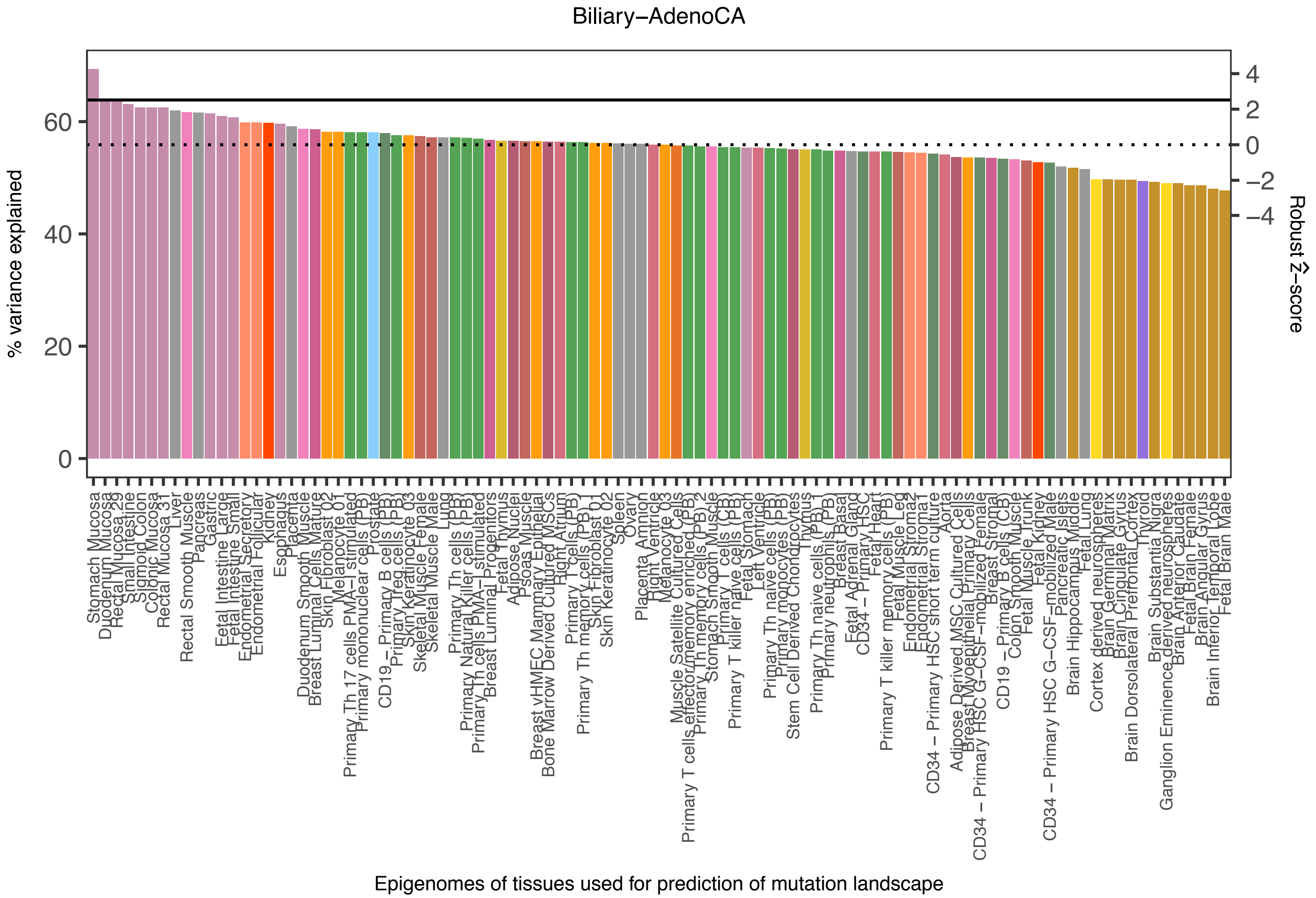

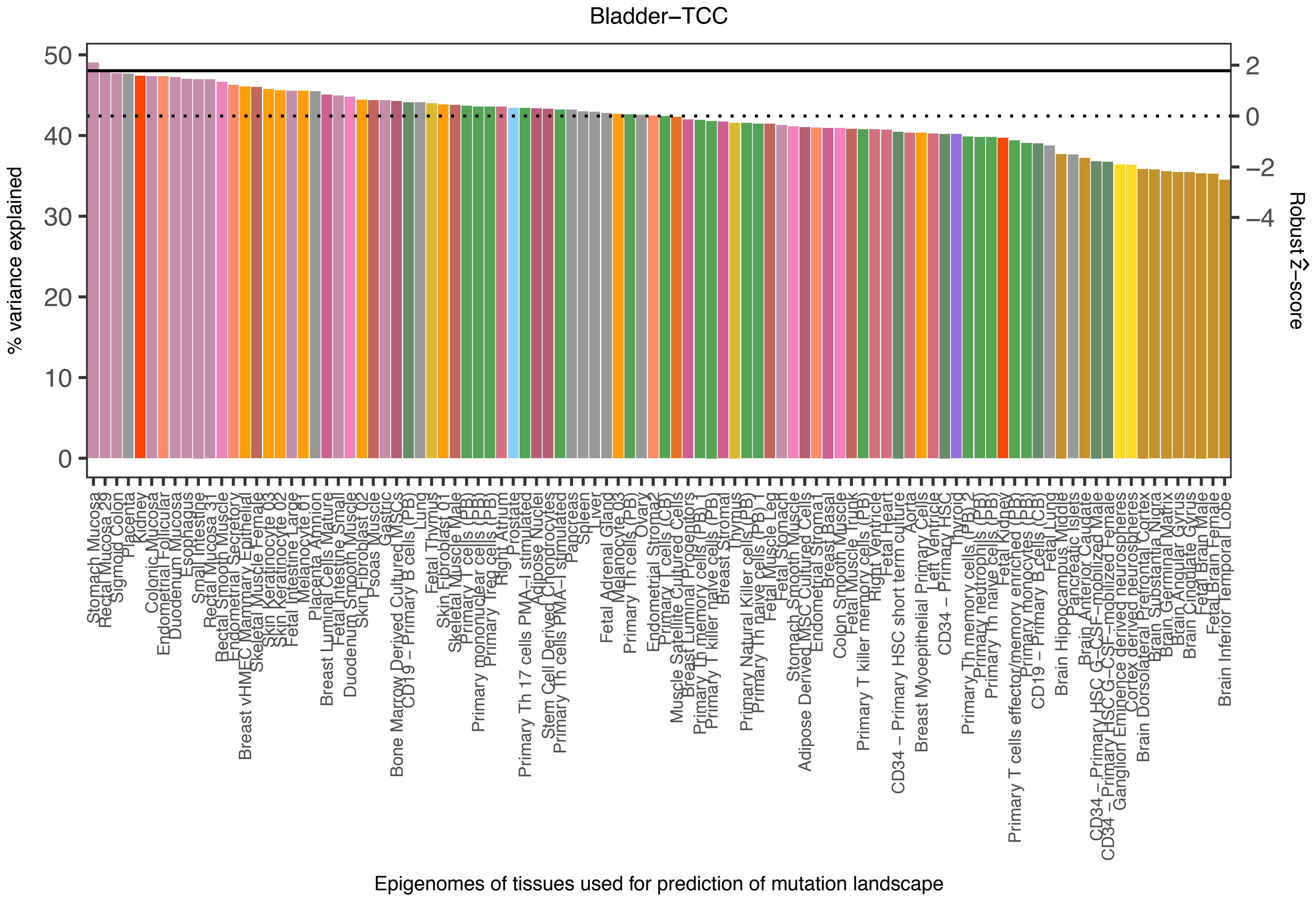

Bladder−TCC
% variance explained
Robust Ẑ−score
Epigenomes of tissues used for prediction of mutation landscape
0
10
20
30
40
50
2
0
−2
−4

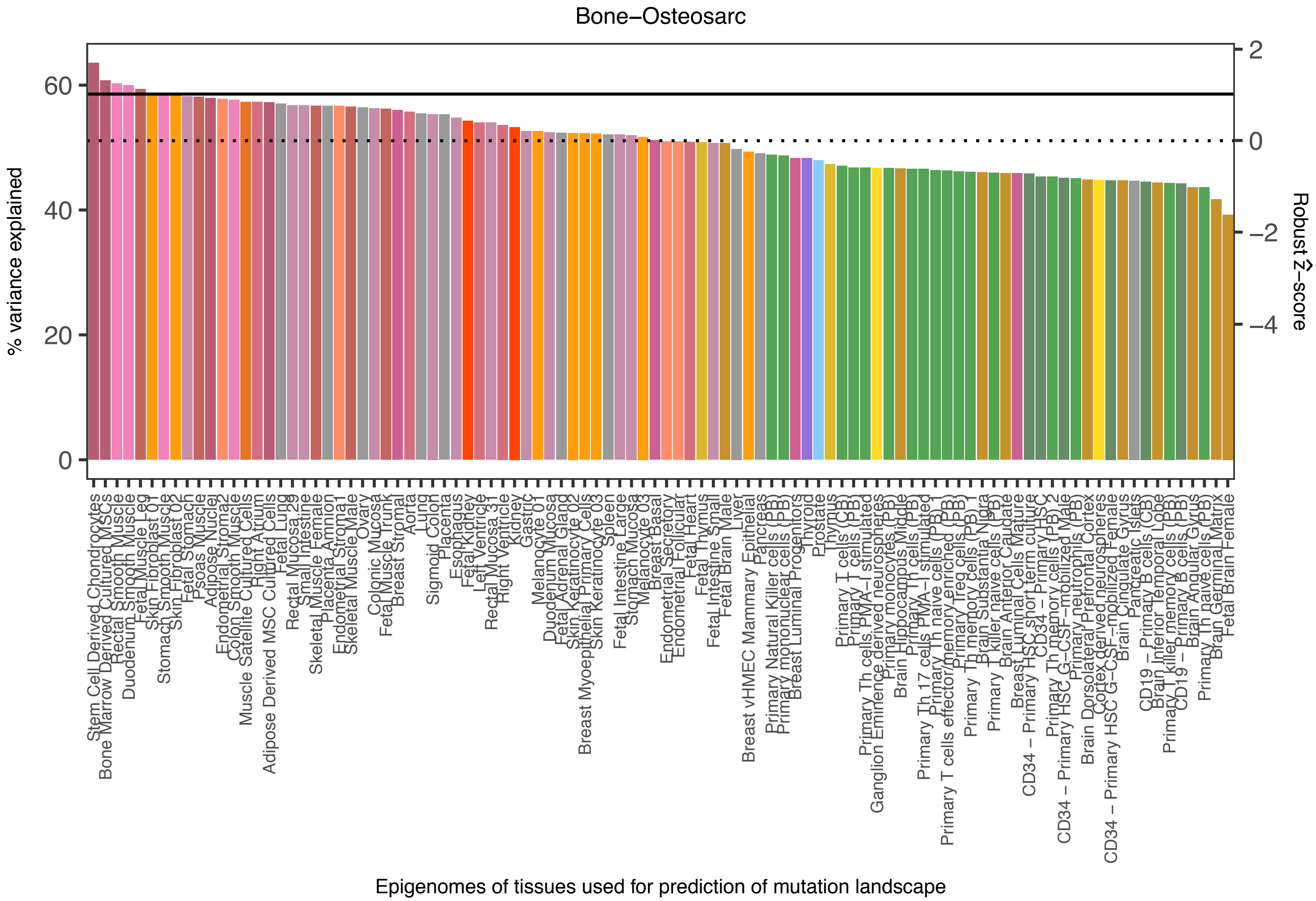
Bone−Osteosarc
% variance explained
Robust Ẑ−score
Epigenomes of tissues used for prediction of mutation landscape
0
20
40
60
2
0
−2
−4
Stem Cell Derived Chondrocytes
Bone Marrow Derived Cultured MSCs
Rectal Smooth Muscle
Duodenum Smooth Muscle
Fetal Muscle Leg
Skin Fibroblast 01
Stomach Smooth Muscle
Skin Fibroblast 02
Fetal Stomach
Psoas Muscle
Adipose Nuclei
Endometrial Stroma2
Colon Smooth Muscle
Muscle Satellite Cultured Cells
Right Atrium
Adipose Derived MSC Cultured Cells
Fetal Lung
Rectal Mucosa 29
Small Intestine
Skeletal Muscle Female
Placenta Amnion
Endometrial Stroma1
Skeletal Muscle Male
Ovary
Colonic Mucosa
Fetal Muscle Trunk
Breast Stromal
Aorta
Lung
Sigmoid Colon
Placenta
Esophagus
Fetal Kidney
Left Ventricle
Rectal Mucosa 31
Right Ventricle
Kidney
Gastric
Melanocyte 01
Duodenum Mucosa
Fetal Adrenal Gland
Skin Keratinocyte 02
Breast Myoepithelial Primary Cells
Skin Keratinocyte 03
Spleen
Fetal Intestine Large
Stomach Mucosa
Melanocyte 03
Breast Basal
Endometrial Secretory
Endometrial Follicular
Fetal Heart
Fetal Thymus
Fetal Intestine Small
Fetal Brain Male
Liver
Breast vHMEC Mammary Epithelial
Pancreas
Primary Natural Killer cells (PB)
Primary mononuclear cells (PB)
Breast Luminal Progenitors
Thyroid
Prostate
Thymus
Primary T cells (PB)
Primary T cells
Primary Th cells PMA−I stimulated
Ganglion Eminence derived neurospheres
Primary monocytes (PB)
Brain Hippocampus Middle
Primary Th cells (PB)
Primary Th 17 cells PMA−I stimulated
Primary Th naive cells (PB)
Primary T cells effector/memory enriched (PB)
Primary Treg cells (PB)
Primary Th memory cells (PB) 1
Brain Substantia Nigra
Primary T killer naive cells (PB)
Brain Anterior Caudate
Breast Luminal Cells Mature
CD34 − Primary HSC short term culture
CD34 − Primary HSC
Primary Th memory cells (PB) 2
CD34 − Primary HSC G−CSF−mobilized Male
Primary neutrophils (PB)
Brain Dorsolateral Prefrontal Cortex
Cortex derived neurospheres
CD34 − Primary HSC G−CSF−mobilized Female
Brain Cingulate Gyrus
Pancreatic Islets
CD19 − Primary B cells (CB)
Brain Inferior Temporal Lobe
Primary T killer memory cells (PB)
CD19 − Primary B cells (PB)
Brain Angular Gyrus
Primary Th naive cells (PB)
Brain Germinal Matrix
Fetal Brain Female

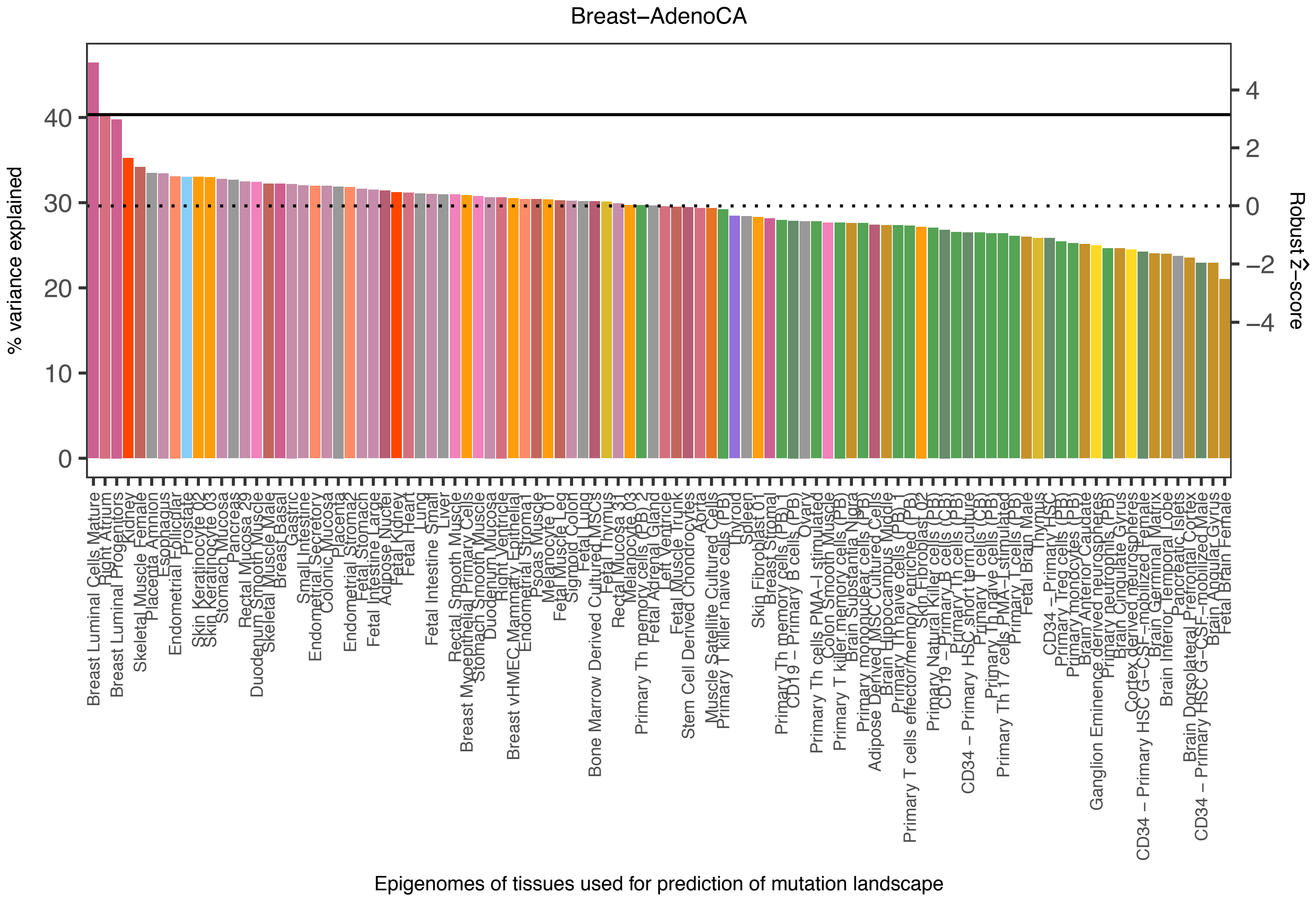
Breast−AdenoCA
% variance explained
Robust Ẑ−score
Epigenomes of tissues used for prediction of mutation landscape
Breast Luminal Cells Mature
Right Atrium
Breast Luminal Progenitors
Kidney
Skeletal Muscle Female
Placenta Amnion
Esophagus
Endometrial Follicular
Prostate
Skin Keratinocyte 02
Skin Keratinocyte 03
Stomach Mucosa
Pancreas
Rectal Mucosa 29
Duodenum Smooth Muscle
Skeletal Muscle Male
Breast Basal
Gastric
Small Intestine
Endometrial Secretory
Colonic Mucosa
Placenta
Endometrial Stromal
Fetal Stomach
Fetal Intestine Large
Adipose Nuclei
Fetal Kidney
Fetal Heart
Lung
Fetal Intestine Small
Liver
Rectal Smooth Muscle
Breast Myoepithelial Primary Cells
Stomach Smooth Muscle
Duodenum Mucosa
Right Ventricle
Breast vHMEC Mammary Epithelial
Endometrial Stromal
Psoas Muscle
Melanocyte 01
Fetal Muscle Leg
Sigmoid Colon
Fetal Lung
Bone Marrow Derived Cultured MSCs
Fetal Thymus
Rectal Mucosa 31
Melanocyte 03
Primary Th memory cells (PB) 2
Fetal Adrenal Gland
Left Ventricle
Fetal Muscle Trunk
Stem Cell Derived Chondrocytes
Aorta
Muscle Satellite Cultured Cells
Primary T killer naive cells (PB)
Thyroid
Spleen
Skin Fibroblast 01
Breast Stromal
Primary Th memory cells (PB)
CD19 − Primary B cells (PB)
Ovary
Primary Th cells PMA−I stimulated
Colon Smooth Muscle
Primary T killer memory cells (PB)
Brain Substantia Nigra
Primary mononuclear cells (PB)
Adipose Derived MSC Cultured Cells
Brain Hippocampus Middle
Primary Th naive cells (PB) 1
Primary T cells effector/memory enriched (PB)
Skin Fibroblast 02
Primary Natural Killer cells (PB)
CD19 − Primary B cells (CB)
Primary Th cells (PB)
CD34 − Primary HSC short term culture
Primary T cells (CB)
Primary Th naive cells (PB)
Primary Th 17 cells PMA−I stimulated
Primary T cells (PB)
Fetal Brain Male
Thymus
CD34 − Primary HSC
Primary Treg cells (PB)
Primary monocytes (PB)
Brain Anterior Caudate
Ganglion Eminence derived neurospheres
Primary neutrophils (PB)
Brain Cingulate Gyrus
Cortex derived neurospheres
CD34 − Primary HSC G−CSF−mobilized Female
Brain Germinal Matrix
Brain Inferior Temporal Lobe
Pancreatic Islets
Brain Dorsolateral Prefrontal Cortex
CD34 − Primary HSC G−CSF−mobilized Male
Brain Angular Gyrus
Fetal Brain Female

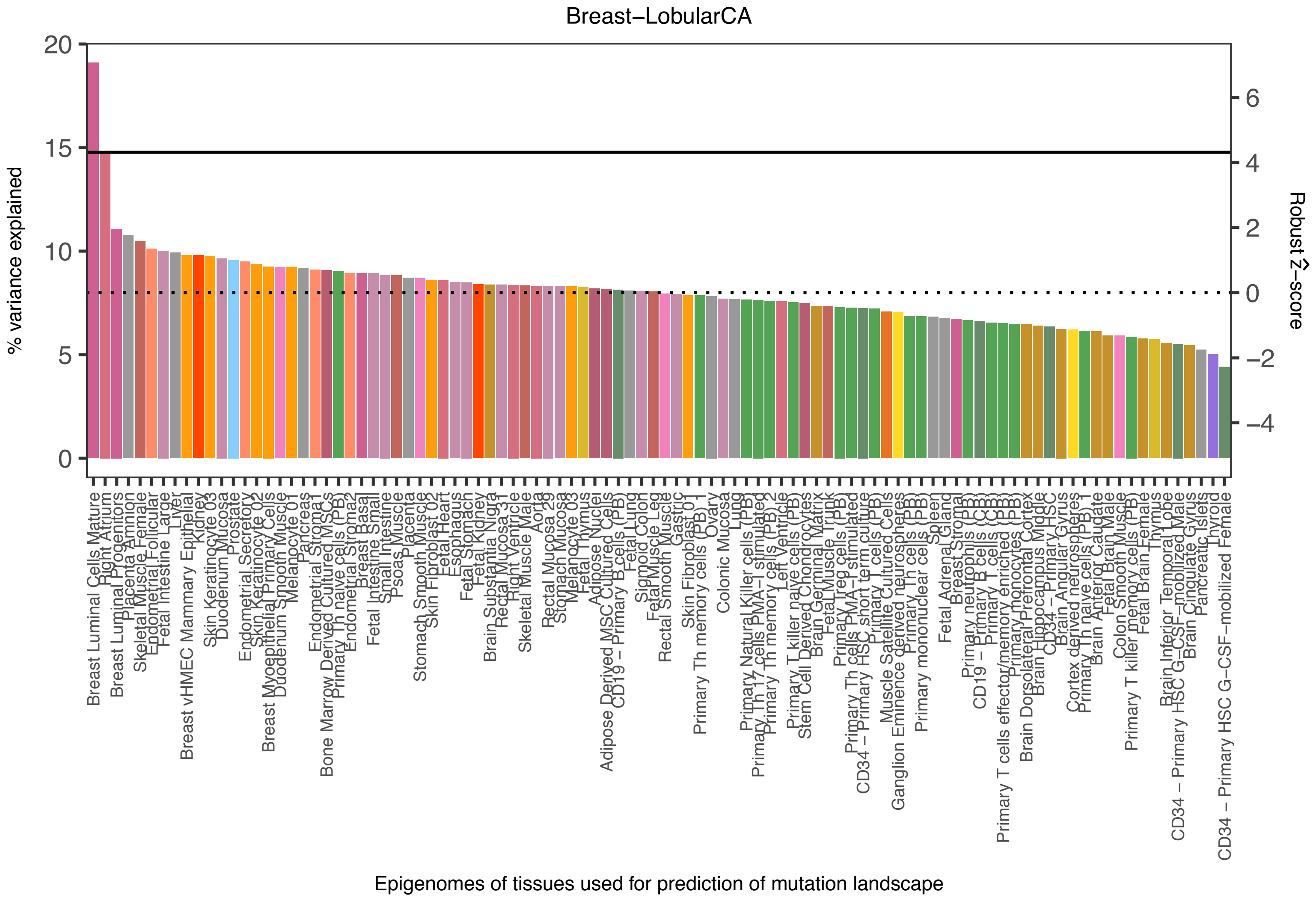
Breast−LobularCA
% variance explained
Robust Ẑ−score
20
15
10
5
0
6
4
2
0
−2
−4
Breast Luminal Cells Mature
Right Atrium
Breast Luminal Progenitors
Placenta Amnion
Skeletal Muscle Female
Endometrial Follicular
Fetal Intestine Large
Liver
Breast vHMEC Mammary Epithelial
Kidney
Skin Keratinocyte 03
Duodenum Mucosa
Prostate
Endometrial Secretory
Skin Keratinocyte 02
Breast Myoepithelial Primary Cells
Duodenum Smooth Muscle
Melanocyte 01
Pancreas
Endometrial Stromal
Bone Marrow Derived Cultured MSCs
Primary Th naive cells (PB)
Endometrial Stromal2
Breast Basal
Fetal Intestine Small
Small Intestine
Psoas Muscle
Placenta
Stomach Smooth Muscle
Skin Fibroblast 02
Fetal Heart
Esophagus
Fetal Stomach
Fetal Kidney
Brain Substantia Nigra
Rectal Mucosa 31
Right Ventricle
Skeletal Muscle Male
Aorta
Rectal Mucosa 29
Stomach Mucosa
Melanocyte 03
Fetal Thymus
Adipose Nuclei
Adipose Derived MSC Cultured Cells
CD19 − Primary B cells (PB)
Fetal Lung
Sigmoid Colon
Fetal Muscle Leg
Rectal Smooth Muscle
Gastric
Skin Fibroblast 01
Primary Th memory cells (PB) 1
Ovary
Colonic Mucosa
Lung
Primary Natural Killer cells (PB)
Primary Th17 cells PMA−I stimulated
Primary Th memory cells (PB) 2
Left Ventricle
Primary T killer naive cells (PB)
Stem Cell Derived Chondrocytes
Brain Germinal Matrix
Fetal Muscle Trunk
Primary Treg cells (PB)
Primary Th cells PMA−I stimulated
CD34 − Primary HSC short term culture
Primary T cells (PB)
Muscle Satellite Cultured Cells
Ganglion Eminence derived neurospheres
Primary Th cells (PB)
Primary mononuclear cells (PB)
Spleen
Fetal Adrenal Gland
Breast Stromal
Primary neutrophils (PB)
CD19 − Primary B cells (CB)
Primary T cells (CB)
Primary T cells effector/memory enriched (PB)
Primary monocytes (PB)
Brain Dorsolateral Prefrontal Cortex
Brain Hippocampus Middle
CD34 − Primary HSC
Brain Angular Gyrus
Cortex derived neurospheres
Primary Th naive cells (PB) 1
Brain Anterior Caudate
Fetal Brain Male
Colon Smooth Muscle
Primary T killer memory cells (PB)
Fetal Brain Female
Thymus
Brain Inferior Temporal Lobe
CD34 − Primary HSC G−CSF−mobilized Male
Brain Cingulate Gyrus
Pancreatic Islets
Thyroid
CD34 − Primary HSC G−CSF−mobilized Female
Epigenomes of tissues used for prediction of mutation landscape

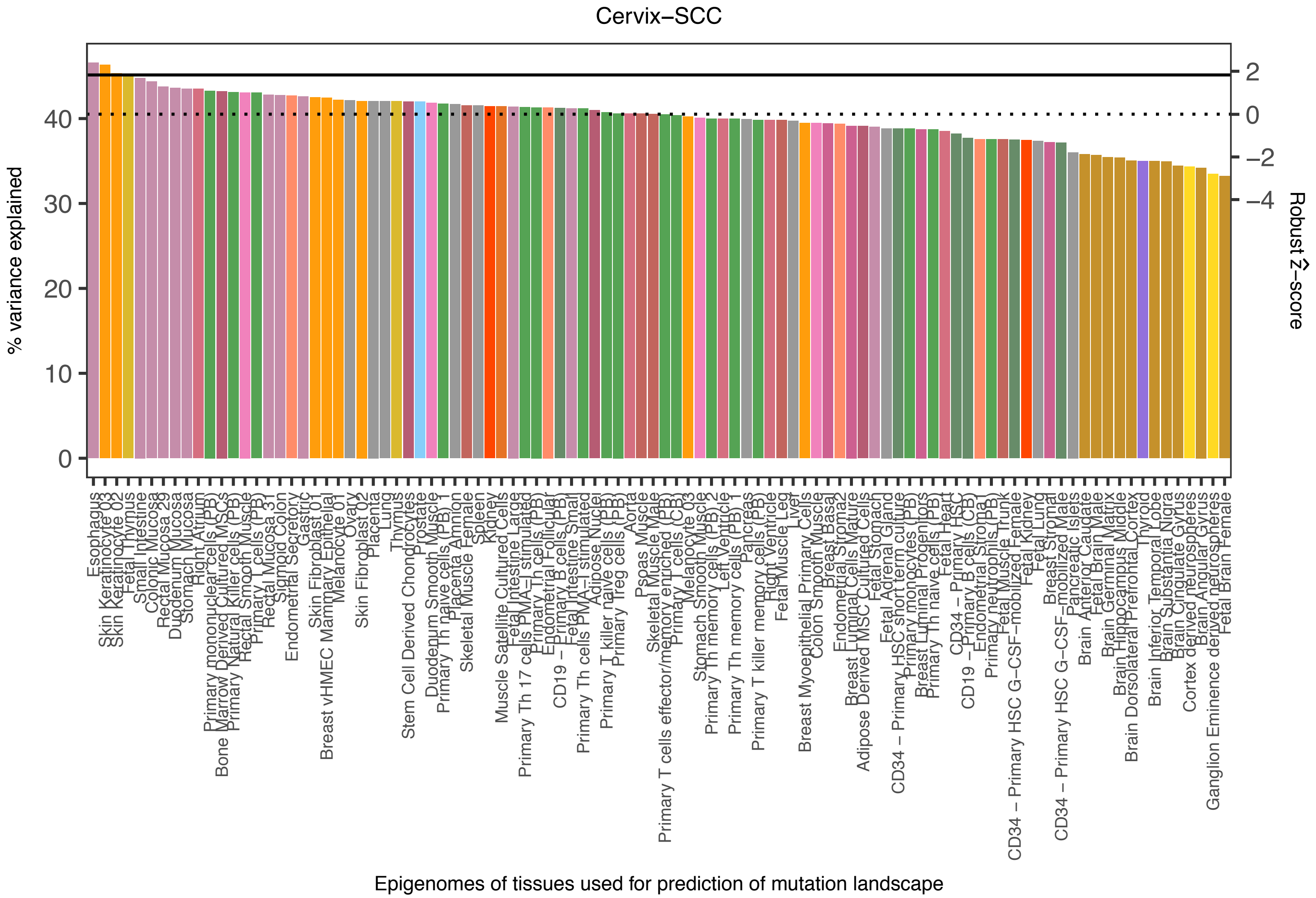

Cervix−SCC
% variance explained
0
10
20
30
40
Robust Ẑ−score
2
0
−2
−4
Epigenomes of tissues used for prediction of mutation landscape

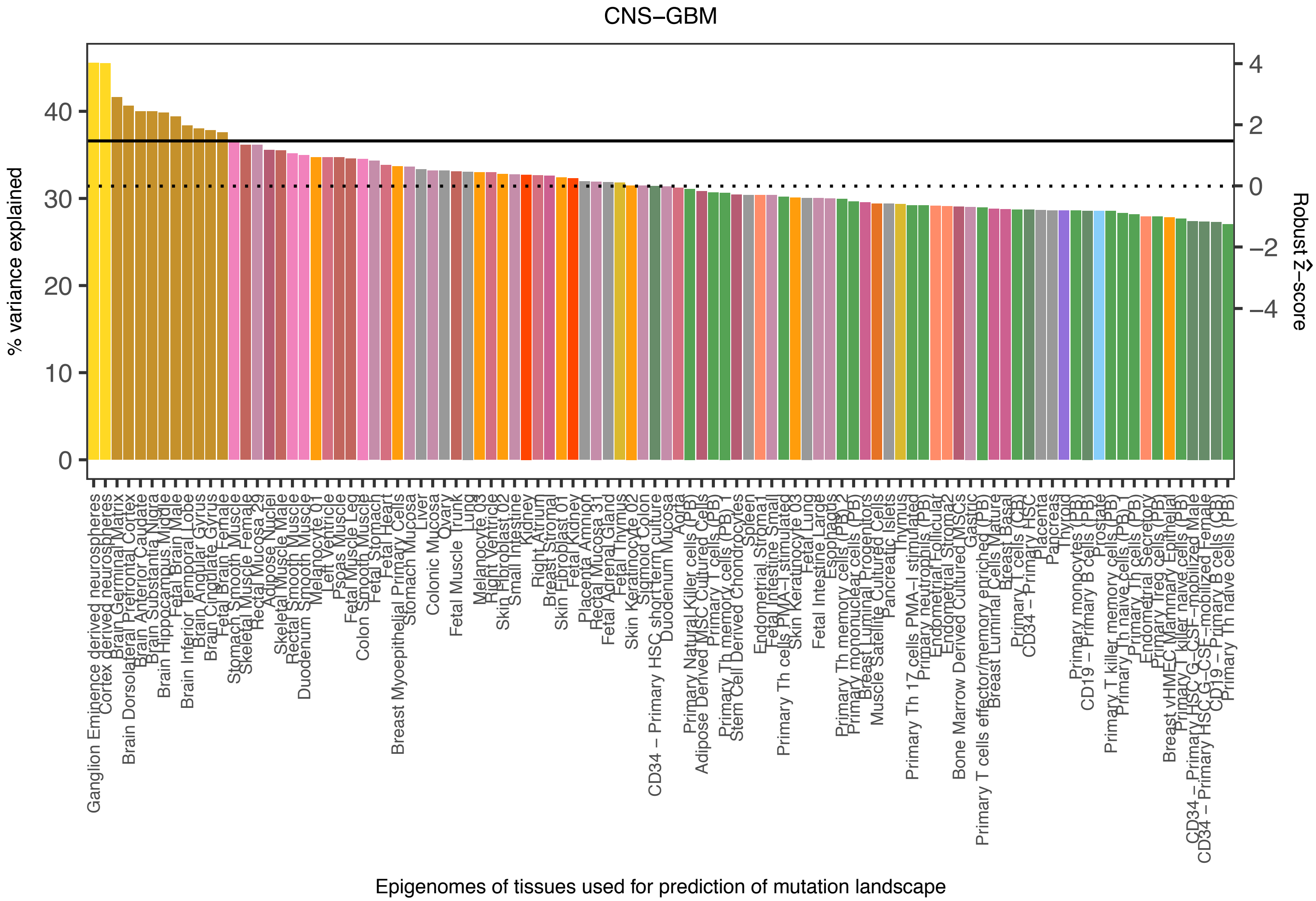

CNS−GBM
% variance explained
Robust Ẑ−score
0
10
20
30
40
4
2
0
−2
−4
Ganglion Eminence derived neurospheres
Cortex derived neurospheres
Brain Germinal Matrix
Brain Dorsolateral Prefrontal Cortex
Brain Anterior Caudate
Brain Substantia Nigra
Brain Hippocampus Middle
Fetal Brain Male
Brain Inferior Temporal Lobe
Brain Angular Gyrus
Brain Cingulate Gyrus
Fetal Brain Female
Stomach Smooth Muscle
Skeletal Muscle Female
Rectal Mucosa 29
Adipose Nuclei
Skeletal Muscle Male
Rectal Smooth Muscle
Duodenum Smooth Muscle
Melanocyte 01
Left Ventricle
Psoas Muscle
Fetal Muscle Leg
Colon Smooth Muscle
Fetal Stomach
Fetal Heart
Breast Myoepithelial Primary Cells
Stomach Mucosa
Liver
Colonic Mucosa
Ovary
Fetal Muscle Trunk
Lung
Melanocyte 03
Right Ventricle
Skin Fibroblast 02
Small Intestine
Kidney
Right Atrium
Breast Stromal
Skin Fibroblast 01
Fetal Kidney
Placenta Amnion
Rectal Mucosa 31
Fetal Adrenal Gland
Fetal Thymus
Skin Keratinocyte 02
Sigmoid Colon
CD34 − Primary HSC short term culture
Duodenum Mucosa
Aorta
Primary Natural Killer cells (PB)
Adipose Derived MSC Cultured Cells
Primary T cells (PB)
Primary Th memory cells (PB) 1
Stem Cell Derived Chondrocytes
Spleen
Endometrial Stromal 1
Fetal Intestine Small
Primary Th cells PMA−I stimulated
Skin Keratinocyte 03
Fetal Lung
Fetal Intestine Large
Esophagus
Primary Th memory cells (PB) 2
Primary mononuclear cells (PB)
Breast Luminal Progenitors
Muscle Satellite Cultured Cells
Pancreatic Islets
Thymus
Primary Th 17 cells PMA−I stimulated
Primary neutrophils (PB)
Endometrial Follicular
Endometrial Stromal 2
Bone Marrow Derived Cultured MSCs
Gastric
Primary T cells effector/memory enriched (PB)
Breast Luminal Cells Mature
Breast Basal
Primary T cells (CB)
CD34 − Primary HSC
Placenta
Pancreas
Thyroid
Primary monocytes (PB)
CD19 − Primary B cells (PB)
Prostate
Primary T killer memory cells (PB)
Primary Th naive cells (PB) 1
Primary Th cells (PB)
Endometrial Secretory
Primary Treg cells (PB)
Breast vHMEC Mammary Epithelial
Primary T killer naive cells (PB)
CD34 − Primary HSC G−CSF−mobilized Male
CD34 − Primary HSC G−CSF−mobilized Female
CD19 − Primary B cells (CB)
Primary Th naive cells (PB)
Epigenomes of tissues used for prediction of mutation landscape

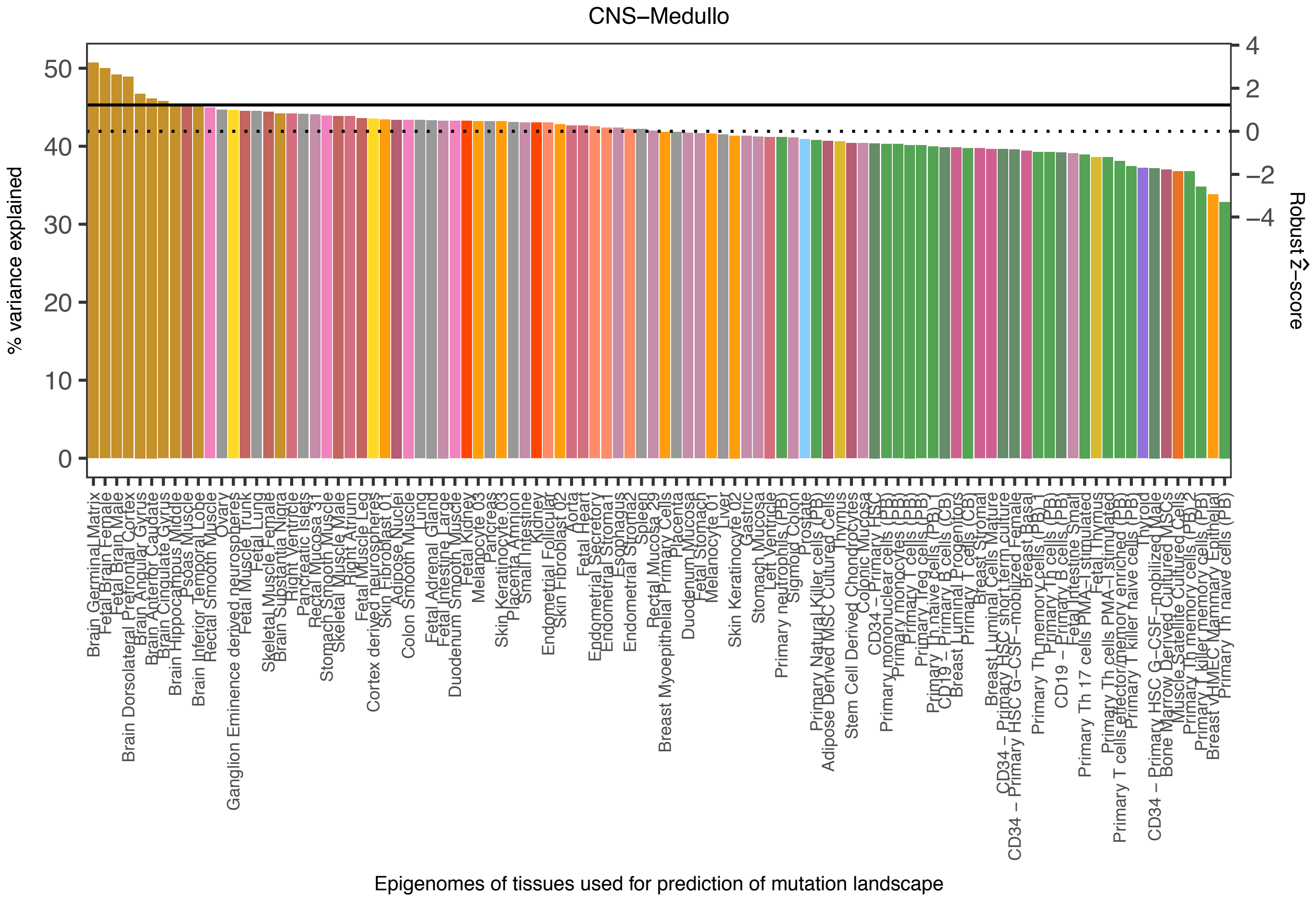
CNS−Medullo
% variance explained
0
10
20
30
40
50
Robust Ẑ−score
4
2
0
−2
−4
Brain Germinal Matrix
Fetal Brain Female
Fetal Brain Male
Brain Dorsolateral Prefrontal Cortex
Brain Angular Gyrus
Brain Anterior Caudate
Brain Cingulate Gyrus
Brain Hippocampus Middle
Psoas Muscle
Brain Inferior Temporal Lobe
Rectal Smooth Muscle
Ovary
Ganglion Eminence derived neurospheres
Fetal Muscle Trunk
Fetal Lung
Skeletal Muscle Female
Brain Substantia Nigra
Right Ventricle
Pancreatic Islets
Rectal Mucosa 31
Stomach Smooth Muscle
Skeletal Muscle Male
Right Atrium
Fetal Muscle Leg
Cortex derived neurospheres
Skin Fibroblast 01
Adipose Nuclei
Colon Smooth Muscle
Lung
Fetal Adrenal Gland
Fetal Intestine Large
Duodenum Smooth Muscle
Fetal Kidney
Melanocyte 03
Pancreas
Skin Keratinocyte 03
Placenta Amnion
Small Intestine
Kidney
Endometrial Follicular
Skin Fibroblast 02
Aorta
Fetal Heart
Endometrial Secretory
Endometrial Stroma1
Esophagus
Endometrial Stroma2
Spleen
Rectal Mucosa 29
Breast Myoepithelial Primary Cells
Placenta
Duodenum Mucosa
Fetal Stomach
Melanocyte 01
Liver
Skin Keratinocyte 02
Gastric
Stomach Mucosa
Left Ventricle
Primary neutrophils (PB)
Sigmoid Colon
Prostate
Primary Natural Killer cells (PB)
Adipose Derived MSC Cultured Cells
Thymus
Stem Cell Derived Chondrocytes
Colonic Mucosa
CD34 − Primary HSC
Primary mononuclear cells (PB)
Primary monocytes (PB)
Primary T cells (PB)
Primary Treg cells (PB)
Primary Th naive cells (PB)
CD19 − Primary B cells (CB)
Breast Luminal Progenitors
Primary T cells (CB)
Breast Stromal
Breast Luminal Cells Mature
CD34 − Primary HSC short term culture
CD34 − Primary HSC G−CSF−mobilized Female
Breast Basal
Primary Th memory cells (PB) 1
Primary Th cells (PB)
CD19 − Primary B cells (PB)
Fetal Intestine Small
Primary Th 17 cells PMA−I stimulated
Fetal Thymus
Primary Th cells PMA−I stimulated
Primary T cells effector/memory enriched (PB)
Primary T killer naive cells (PB)
Thyroid
CD34 − Primary HSC G−CSF−mobilized Male
Bone Marrow Derived Cultured MSCs
Muscle Satellite Cultured Cells
Primary Th memory cells (PB) 2
Primary T killer memory cells (PB)
Breast vHMEC Mammary Epithelial
Primary Th naive cells (PB)
Epigenomes of tissues used for prediction of mutation landscape

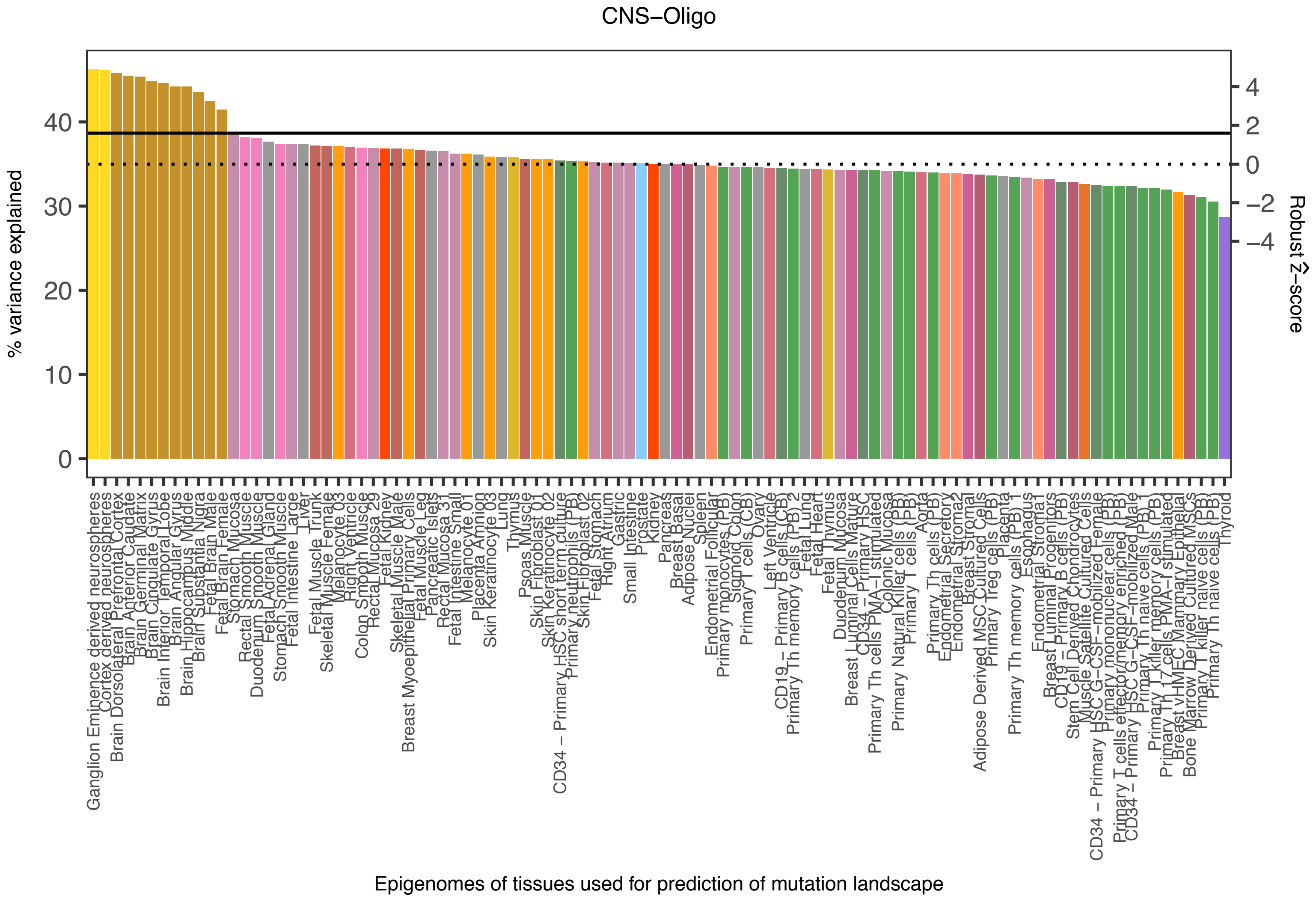
CNS−Oligo
% variance explained
0
10
20
30
40
Robust Ẑ−score
4
2
0
−2
−4
Epigenomes of tissues used for prediction of mutation landscape
Ganglion Eminence derived neurospheres
Cortex derived neurospheres
Brain Dorsolateral Prefrontal Cortex
Brain Anterior Caudate
Brain Germinal Matrix
Brain Cingulate Gyrus
Brain Inferior Temporal Lobe
Brain Angular Gyrus
Brain Hippocampus Middle
Brain Substantia Nigra
Fetal Brain Male
Fetal Brain Female
Stomach Mucosa
Rectal Smooth Muscle
Duodenum Smooth Muscle
Fetal Adrenal Gland
Stomach Smooth Muscle
Fetal Intestine Large
Liver
Fetal Muscle Trunk
Skeletal Muscle Female
Melanocyte 03
Right Ventricle
Colon Smooth Muscle
Rectal Mucosa 29
Fetal Kidney
Skeletal Muscle Male
Breast Myoepithelial Primary Cells
Fetal Muscle Leg
Pancreatic Islets
Rectal Mucosa 31
Fetal Intestine Small
Melanocyte 01
Placenta Amnion
Skin Keratinocyte 03
Lung
Thymus
Psoas Muscle
Skin Fibroblast 01
Skin Keratinocyte 02
CD34 − Primary HSC short term culture
Primary neutrophils (PB)
Skin Fibroblast 02
Fetal Stomach
Right Atrium
Gastric
Small Intestine
Prostate
Kidney
Pancreas
Breast Basal
Adipose Nuclei
Spleen
Endometrial Follicular
Primary monocytes (PB)
Sigmoid Colon
Primary T cells (CB)
Ovary
Left Ventricle
CD19 − Primary B cells (CB)
Primary Th memory cells (PB) 2
Fetal Lung
Fetal Heart
Fetal Thymus
Duodenum Mucosa
Breast Luminal Cells Mature
CD34 − Primary HSC
Primary Th cells PMA−I stimulated
Colonic Mucosa
Primary Natural Killer cells (PB)
Primary T cells (PB)
Aorta
Primary Th cells (PB)
Endometrial Secretory
Endometrial Stroma2
Breast Stromal
Adipose Derived MSC Cultured Cells
Primary Treg cells (PB)
Placenta
Primary Th memory cells (PB) 1
Esophagus
Endometrial Stroma1
Breast Luminal Progenitors
CD19 − Primary B cells (PB)
Stem Cell Derived Chondrocytes
Muscle Satellite Cultured Cells
CD34 − Primary HSC G−CSF−mobilized Female
Primary mononuclear cells (PB)
Primary T cells effector/memory enriched (PB)
CD34 − Primary HSC G−CSF−mobilized Male
Primary Th naive cells (PB) 1
Primary T killer memory cells (PB)
Primary Th 17 cells PMA−I stimulated
Breast vHMEC Mammary Epithelial
Bone Marrow Derived Cultured MSCs
Primary T killer naive cells (PB)
Primary Th naive cells (PB)
Thyroid

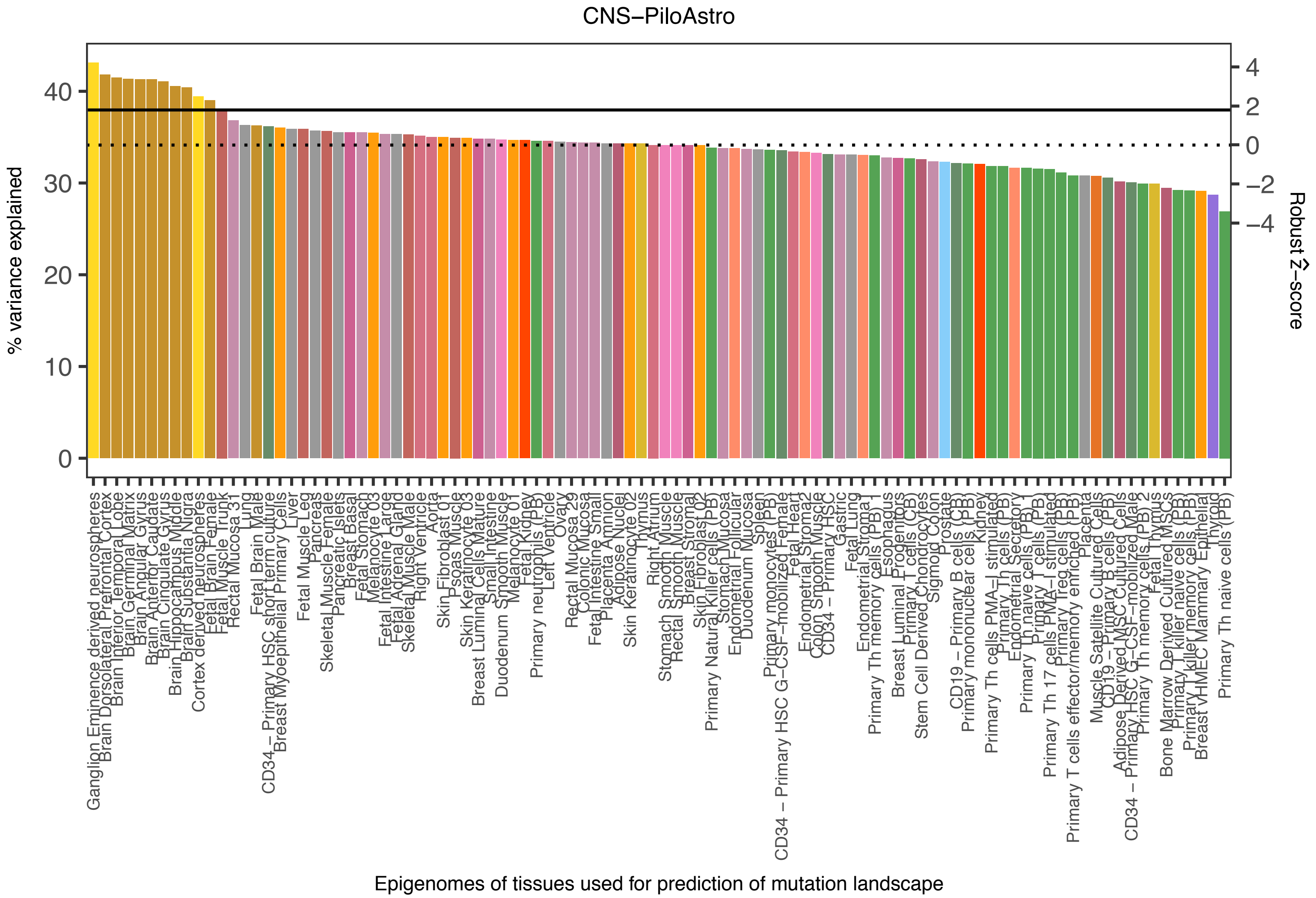
CNS−PiloAstro
% variance explained
Robust Z−score
Epigenomes of tissues used for prediction of mutation landscape

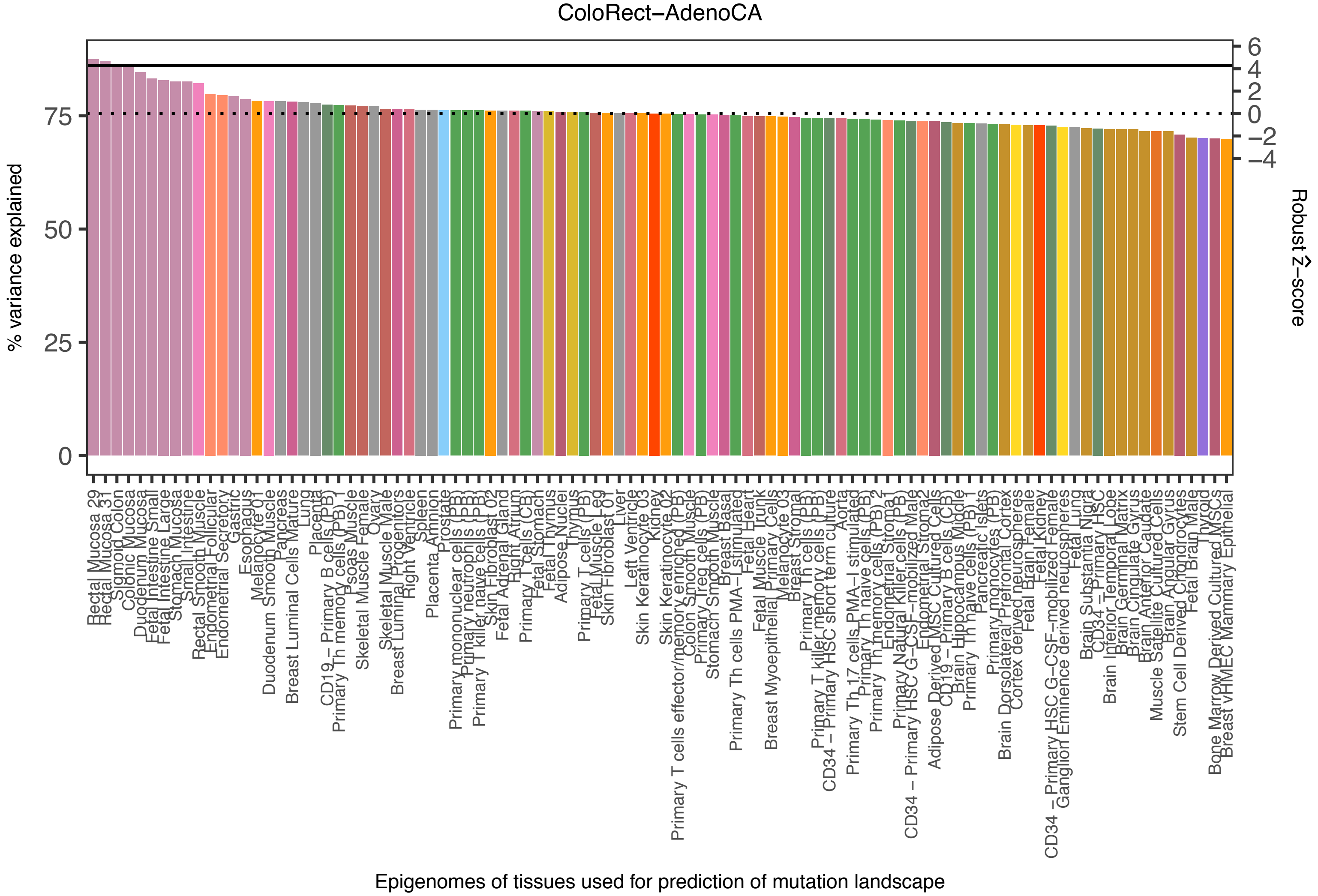
ColoRect−AdenoCA
% variance explained
Robust Ẑ−score
Epigenomes of tissues used for prediction of mutation landscape
0
25
50
75
6
4
2
0
−2
−4

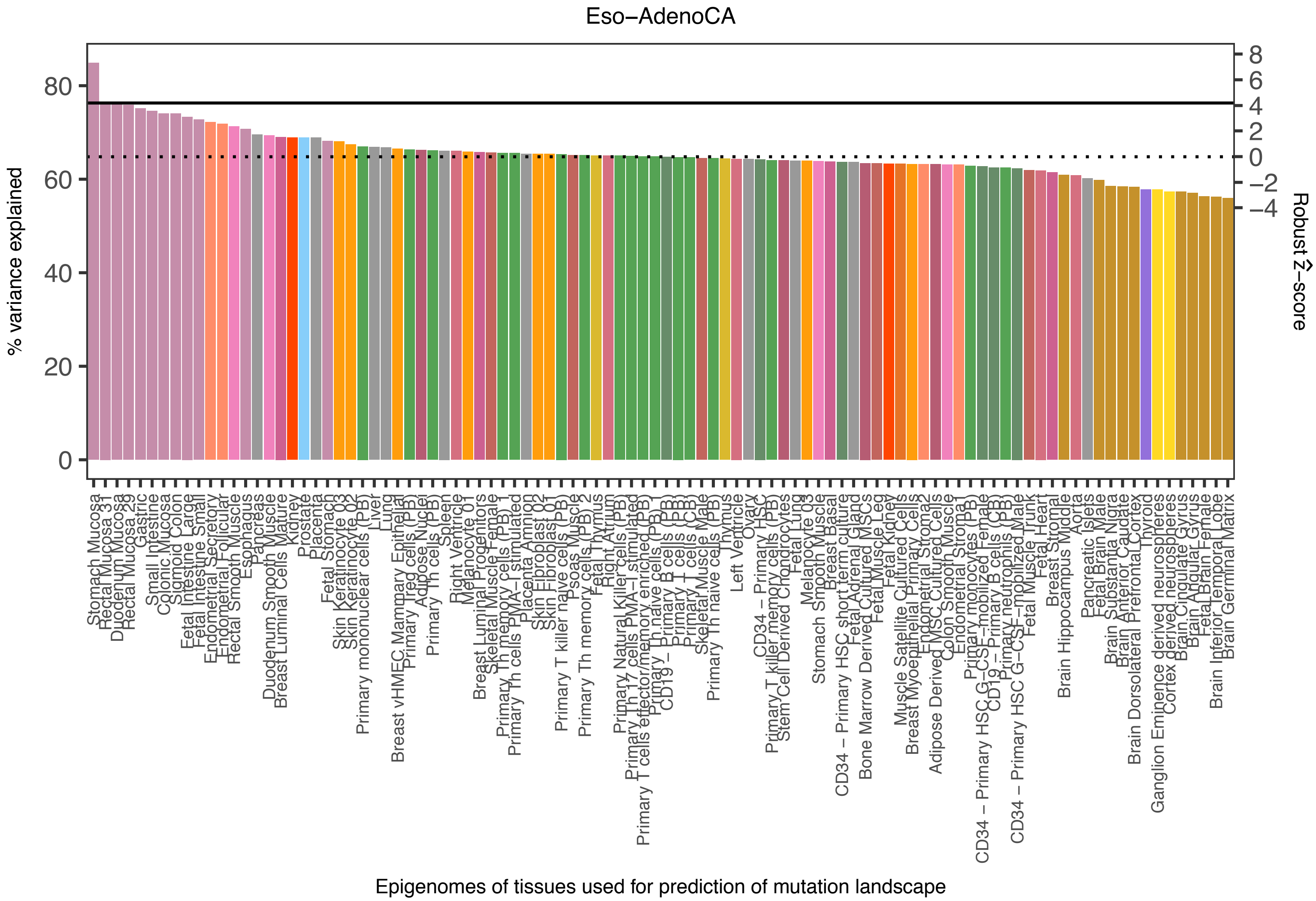

Eso−AdenoCA
% variance explained
0
20
40
60
80
Robust Ẑ−score
8
6
4
2
0
−2
−4
Stomach Mucosa
Rectal Mucosa 31
Duodenum Mucosa
Rectal Mucosa 29
Gastric
Small Intestine
Colonic Mucosa
Sigmoid Colon
Fetal Intestine Large
Fetal Intestine Small
Endometrial Secretory
Endometrial Follicular
Rectal Smooth Muscle
Esophagus
Pancreas
Duodenum Smooth Muscle
Breast Luminal Cells Mature
Kidney
Prostate
Placenta
Fetal Stomach
Skin Keratinocyte 03
Skin Keratinocyte 02
Primary mononuclear cells (PB)
Liver
Lung
Breast vHMEC Mammary Epithelial
Primary Treg cells (PB)
Adipose Nuclei
Primary Th cells (PB)
Spleen
Right Ventricle
Melanocyte 01
Breast Luminal Progenitors
Skeletal Muscle Female
Primary Th memory cells (PB)
Primary Th cells PMA−I stimulated
Placenta Amnion
Skin Fibroblast 02
Skin Fibroblast 01
Primary T killer naive cells (PB)
Psoas Muscle
Primary Th memory cells (PB) 2
Fetal Thymus
Right Atrium
Primary Natural Killer cells (PB)
Primary Th17 cells PMA−I stimulated
Primary T cells effector/memory enriched (PB)
Primary Th naive cells (PB)
CD19 − Primary B cells (PB)
Primary T cells (CB)
Primary T cells (PB)
Skeletal Muscle Male
Primary Th naive cells (CB)
Thymus
Left Ventricle
Ovary
CD34 − Primary HSC
Primary T killer memory cells (PB)
Stem Cell Derived Chondrocytes
Fetal Lung
Melanocyte 03
Stomach Smooth Muscle
Breast Basal
CD34 − Primary HSC short term culture
Fetal Adrenal Gland
Bone Marrow Derived Cultured MSCs
Fetal Muscle Leg
Fetal Kidney
Muscle Satellite Cultured Cells
Breast Myoepithelial Primary Cells
Endometrial Stroma2
Adipose Derived MSC Cultured Cells
Colon Smooth Muscle
Endometrial Stroma1
Primary monocytes (PB)
CD34 − Primary HSC G−CSF−mobilized Female
CD19 − Primary B cells (CB)
Primary neutrophils (PB)
CD34 − Primary HSC G−CSF−mobilized Male
Fetal Muscle Trunk
Fetal Heart
Breast Stromal
Brain Hippocampus Middle
Aorta
Pancreatic Islets
Fetal Brain Male
Brain Substantia Nigra
Brain Anterior Caudate
Brain Dorsolateral Prefrontal Cortex
Thyroid
Ganglion Eminence derived neurospheres
Cortex derived neurospheres
Brain Cingulate Gyrus
Brain Angular Gyrus
Fetal Brain Female
Brain Inferior Temporal Lobe
Brain Germinal Matrix
Epigenomes of tissues used for prediction of mutation landscape

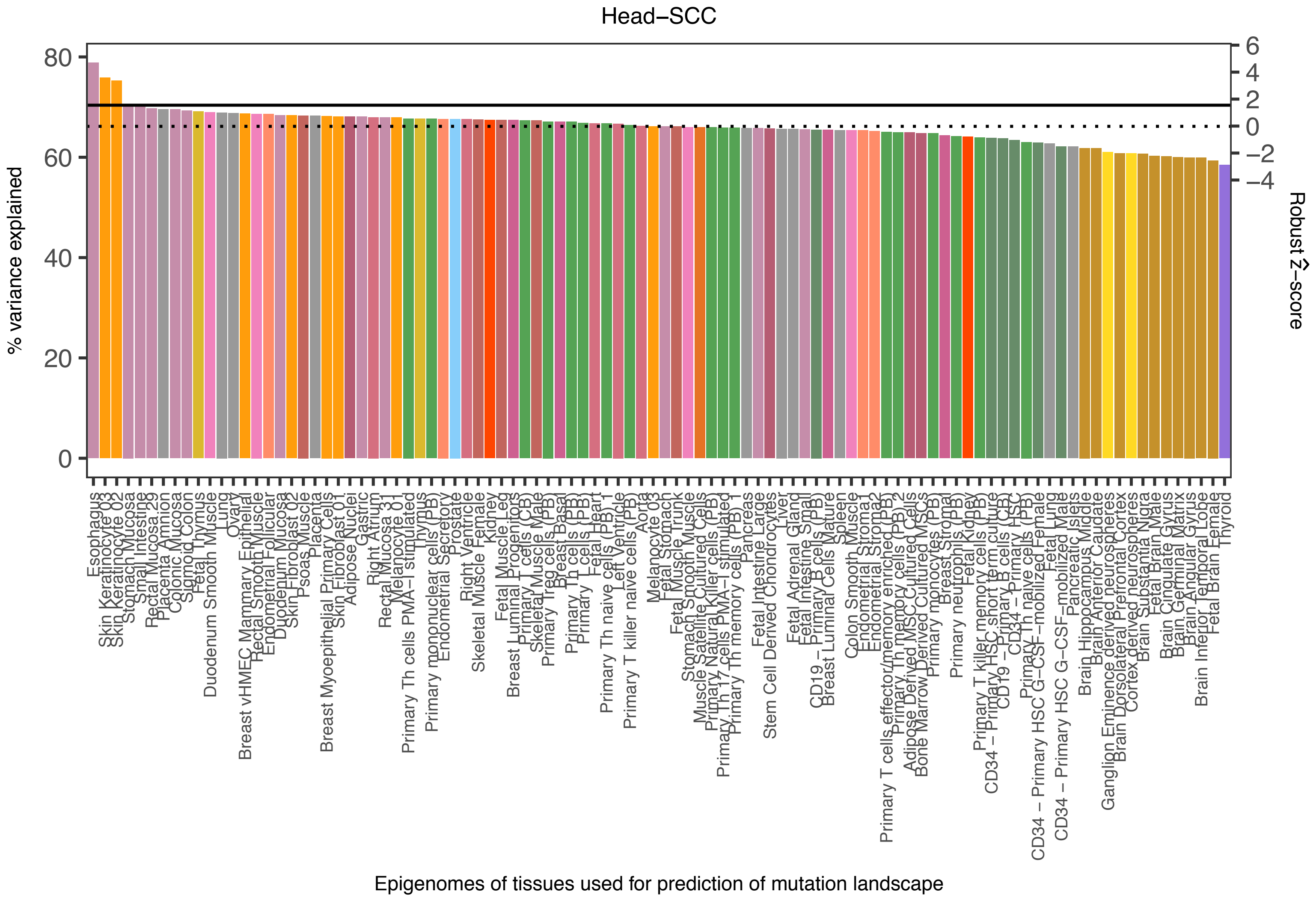
Head−SCC
% variance explained
80
60
40
20
0
Robust Ẑ−score
6
4
2
0
−2
−4
Epigenomes of tissues used for prediction of mutation landscape

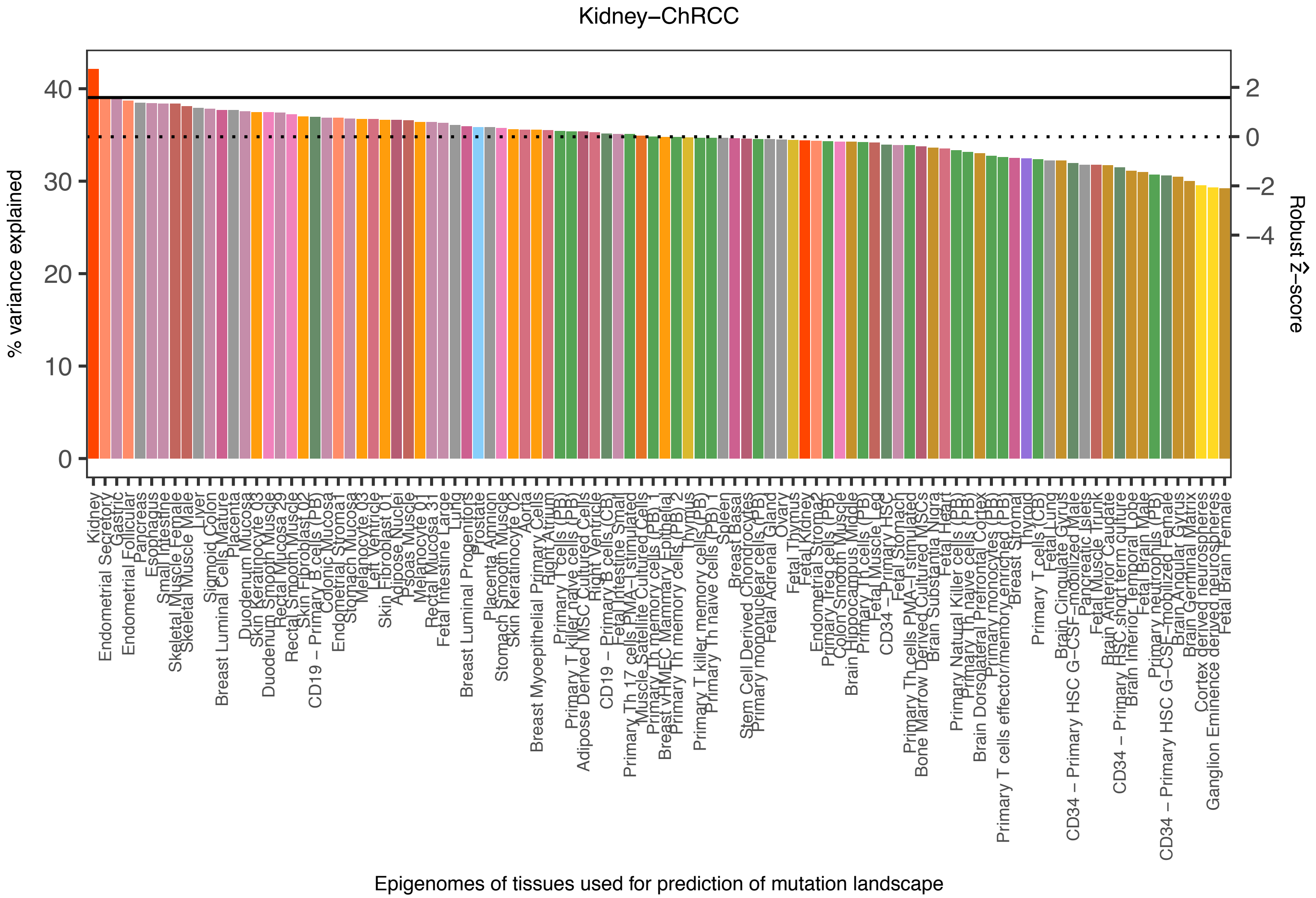

Kidney−ChRCC
% variance explained
0
10
20
30
40
Robust Ẑ−score
2
0
−2
−4
Kidney
Endometrial Secretory
Gastric
Endometrial Follicular
Pancreas
Esophagus
Small Intestine
Skeletal Muscle Female
Skeletal Muscle Male
Liver
Sigmoid Colon
Breast Luminal Cells Mature
Placenta
Duodenum Mucosa
Skin Keratinocyte 03
Duodenum Smooth Muscle
Rectal Mucosa 29
Rectal Smooth Muscle
Skin Fibroblast 02
CD19 − Primary B cells (PB)
Colonic Mucosa
Endometrial Stromal
Stomach Mucosa
Melanocyte 03
Left Ventricle
Skin Fibroblast 01
Adipose Nuclei
Psoas Muscle
Melanocyte 01
Rectal Mucosa 31
Fetal Intestine Large
Lung
Breast Luminal Progenitors
Prostate
Placenta Amnion
Stomach Smooth Muscle
Skin Keratinocyte 02
Aorta
Breast Myoepithelial Primary Cells
Right Atrium
Primary T cells (PB)
Primary T killer naive cells (PB)
Adipose Derived MSC Cultured Cells
Right Ventricle
CD19 − Primary B cells (CB)
Fetal Intestine Small
Primary Th 17 cells PMA−I stimulated
Muscle Satellite Cultured Cells
Primary Th memory cells (PB) 1
Breast vHMEC Mammary Epithelial
Primary Th memory cells (PB) 2
Thymus
Primary T killer memory cells (PB)
Primary Th naive cells (PB) 1
Spleen
Breast Basal
Stem Cell Derived Chondrocytes
Primary mononuclear cells (PB)
Fetal Adrenal Gland
Ovary
Fetal Thymus
Fetal Kidney
Endometrial Stroma2
Primary Treg cells (PB)
Colon Smooth Muscle
Brain Hippocampus Middle
Primary Th cells (PB)
Fetal Muscle Leg
CD34 − Primary HSC
Fetal Stomach
Primary Th cells PMA−I stimulated
Bone Marrow Derived Cultured MSCs
Brain Substantia Nigra
Fetal Heart
Primary Natural Killer cells (PB)
Primary Th naive cells (PB)
Brain Dorsolateral Prefrontal Cortex
Primary monocytes (PB)
Primary T cells effector/memory enriched (PB)
Breast Stromal
Thyroid
Primary T cells (CB)
Fetal Lung
Brain Cingulate Gyrus
CD34 − Primary HSC G−CSF−mobilized Male
Pancreatic Islets
Fetal Muscle Trunk
Brain Anterior Caudate
CD34 − Primary HSC short term culture
Brain Inferior Temporal Lobe
Fetal Brain Male
Primary neutrophils (PB)
CD34 − Primary HSC G−CSF−mobilized Female
Brain Angular Gyrus
Brain Germinal Matrix
Cortex derived neurospheres
Ganglion Eminence derived neurospheres
Fetal Brain Female
Epigenomes of tissues used for prediction of mutation landscape

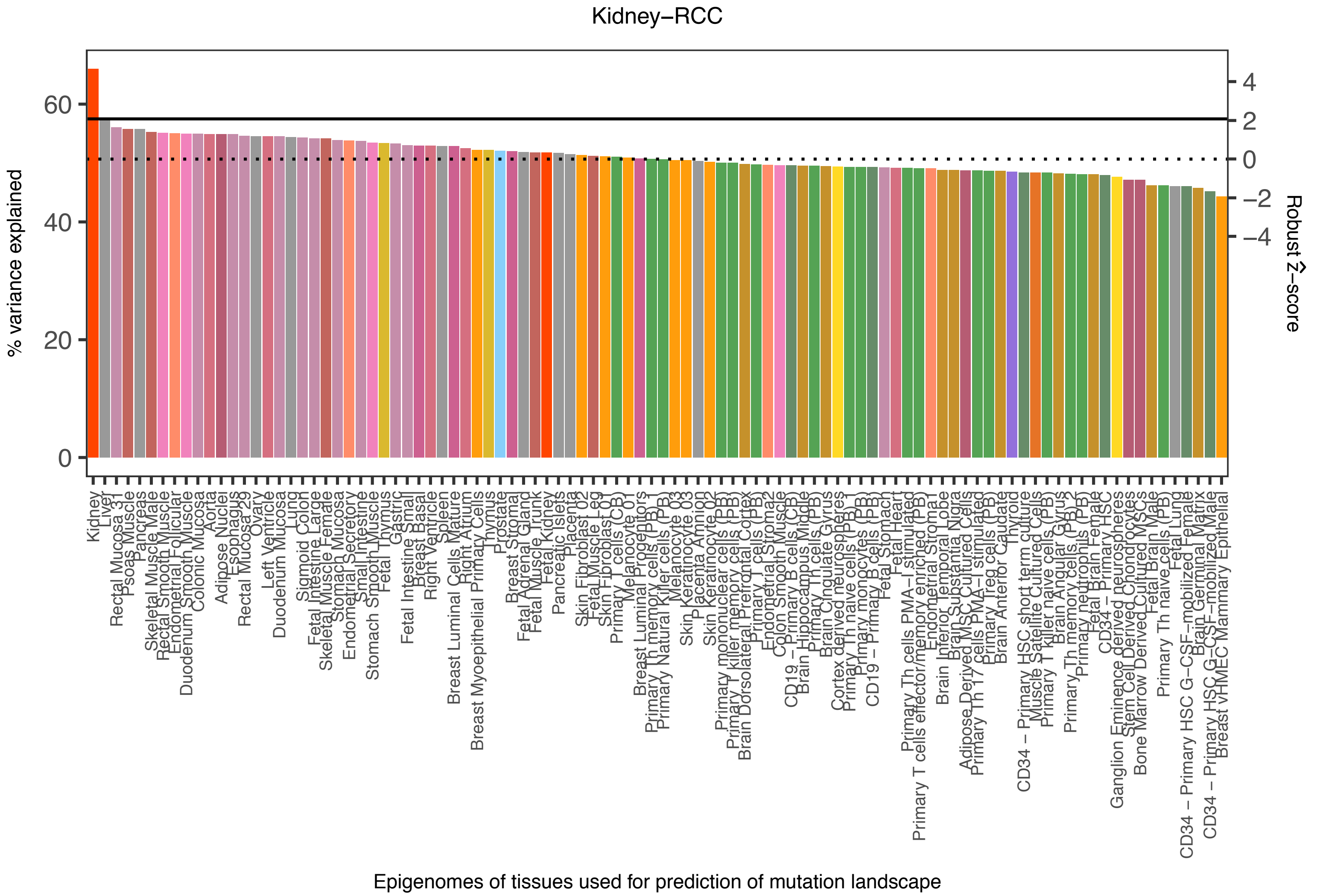

Kidney−RCC
% variance explained
0
20
40
60
Robust Ẑ−score
4
2
0
−2
−4
Kidney
Liver
Rectal Mucosa 31
Psoas Muscle
Pancreas
Skeletal Muscle Male
Rectal Smooth Muscle
Endometrial Follicular
Duodenum Smooth Muscle
Colonic Mucosa
Aorta
Adipose Nuclei
Esophagus
Rectal Mucosa 29
Ovary
Left Ventricle
Duodenum Mucosa
Lung
Sigmoid Colon
Fetal Intestine Large
Skeletal Muscle Female
Stomach Mucosa
Endometrial Secretory
Small Intestine
Stomach Smooth Muscle
Fetal Thymus
Gastric
Fetal Intestine Small
Breast Basal
Right Ventricle
Spleen
Breast Luminal Cells Mature
Right Atrium
Breast Myoepithelial Primary Cells
Thymus
Prostate
Breast Stromal
Fetal Adrenal Gland
Fetal Muscle Trunk
Fetal Kidney
Pancreatic Islets
Placenta
Skin Fibroblast 02
Fetal Muscle Leg
Skin Fibroblast 01
Primary T cells (CB)
Melanocyte 01
Breast Luminal Progenitors
Primary Th memory cells (PB)
Primary Natural Killer cells (PB)
Melanocyte 03
Skin Keratinocyte 03
Placenta Amnion
Skin Keratinocyte 02
Primary mononuclear cells (PB)
Primary T killer memory cells (PB)
Brain Dorsolateral Prefrontal Cortex
Primary T cells (PB)
Endometrial Stromal 2
Colon Smooth Muscle
CD19 − Primary B cells (CB)
Brain Hippocampus Middle
Primary Th cells (PB)
Brain Cingulate Gyrus
Cortex derived neurospheres
Primary Th naive cells (PB)
Primary monocytes (PB)
CD19 − Primary B cells (PB)
Fetal Stomach
Fetal Heart
Primary Th cells PMA−I stimulated
Primary T cells effector/memory enriched (PB)
Endometrial Stromal 1
Brain Inferior Temporal Lobe
Brain Substantia Nigra
Adipose Derived MSC Cultured Cells
Primary Th 17 cells PMA−I stimulated
Primary Treg cells (PB)
Brain Anterior Caudate
Thyroid
CD34 − Primary HSC short term culture
Muscle Satellite Cultured Cells
Primary T killer naive cells (PB)
Brain Angular Gyrus
Primary Th memory cells (PB)
Primary neutrophils (PB)
Fetal Brain Female
CD34 − Primary HSC
Ganglion Eminence derived neurospheres
Stem Cell Derived Chondrocytes
Bone Marrow Derived Cultured MSCs
Fetal Brain Male
Primary Th naive cells (PB)
Fetal Lung
CD34 − Primary HSC G−CSF−mobilized Female
Brain Germinal Matrix
CD34 − Primary HSC G−CSF−mobilized Male
Breast vHMEC Mammary Epithelial
Epigenomes of tissues used for prediction of mutation landscape

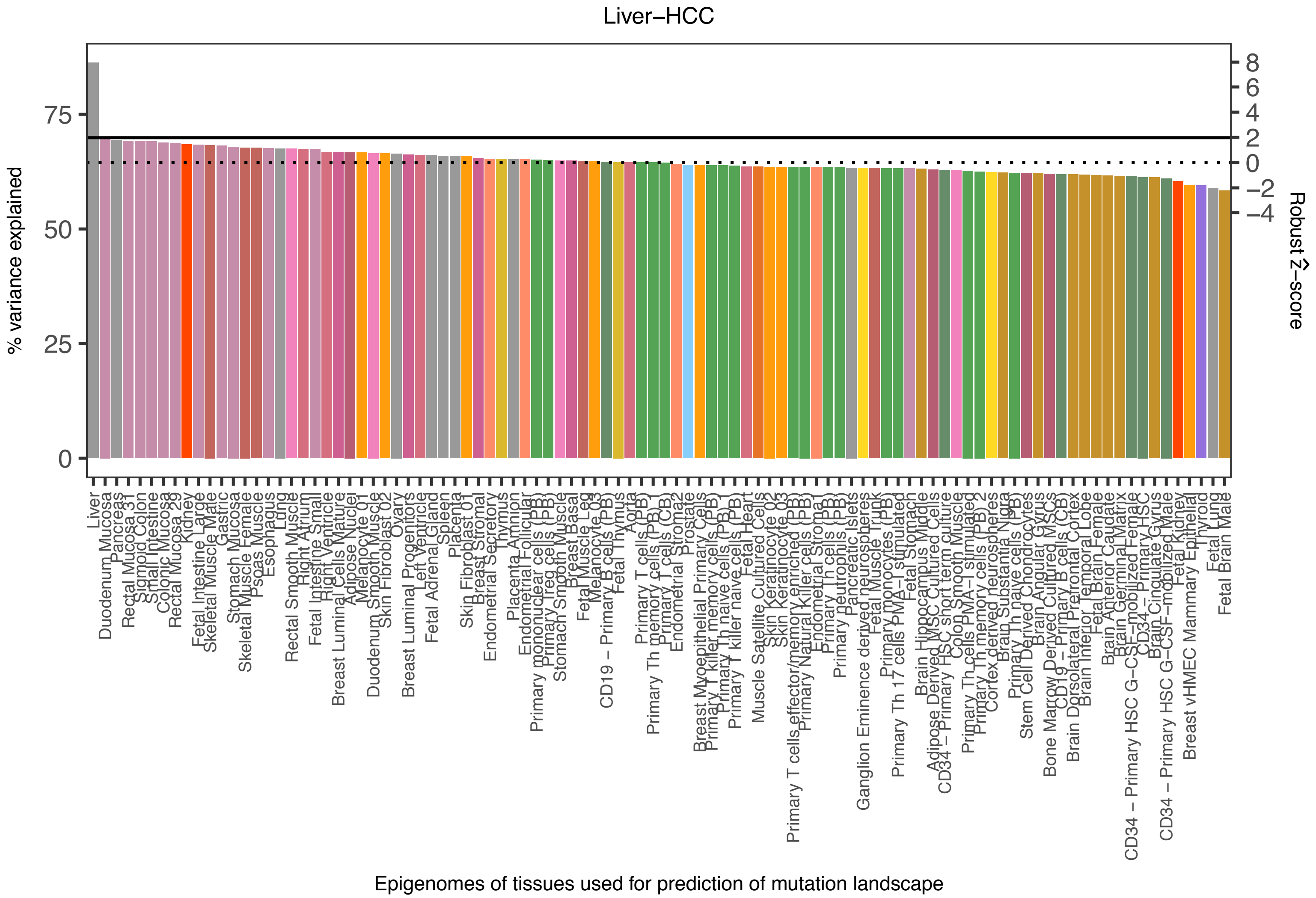

Liver−HCC
% variance explained
75
50
25
0
8
6
4
2
0
−2
−4
Robust Ẑ−score
Epigenomes of tissues used for prediction of mutation landscape

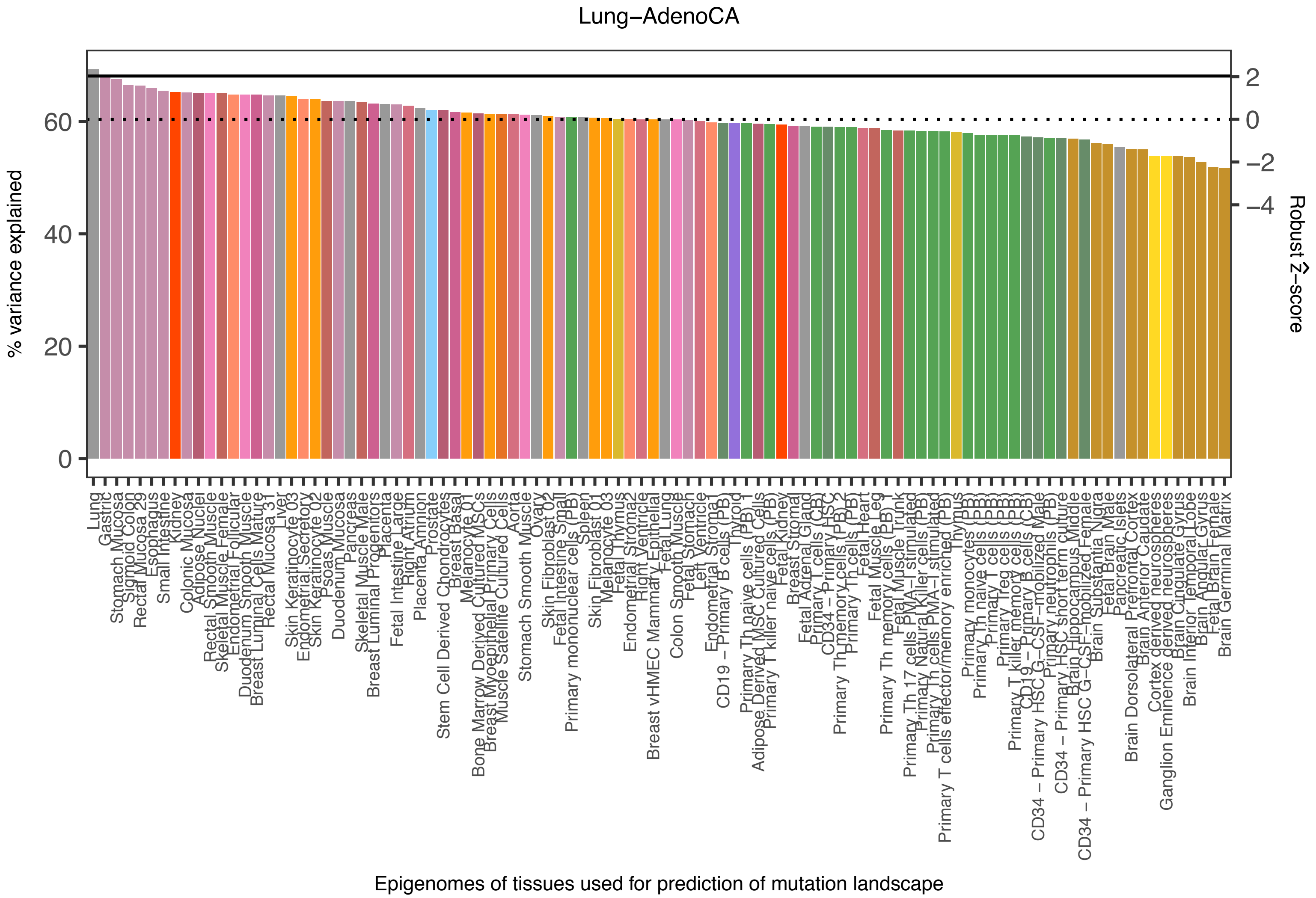

Lung−AdenoCA
% variance explained
0
20
40
60
Robust Ẑ−score
2
0
−2
−4
Lung
Gastric
Stomach Mucosa
Sigmoid Colon
Rectal Mucosa 29
Esophagus
Small Intestine
Kidney
Colonic Mucosa
Adipose Nuclei
Rectal Smooth Muscle
Skeletal Muscle Female
Endometrial Follicular
Duodenum Smooth Muscle
Breast Luminal Cells Mature
Rectal Mucosa 31
Liver
Skin Keratinocyte 03
Endometrial Secretory
Skin Keratinocyte 02
Psoas Muscle
Duodenum Mucosa
Pancreas
Skeletal Muscle Male
Breast Luminal Progenitors
Placenta
Fetal Intestine Large
Right Atrium
Placenta Amnion
Prostate
Stem Cell Derived Chondrocytes
Breast Basal
Melanocyte 01
Bone Marrow Derived Cultured MSCs
Breast Myoepithelial Primary Cells
Muscle Satellite Cultured Cells
Aorta
Stomach Smooth Muscle
Ovary
Skin Fibroblast 02
Fetal Intestine Small
Primary mononuclear cells (PB)
Spleen
Skin Fibroblast 01
Melanocyte 03
Fetal Thymus
Endometrial Stromal 2
Right Ventricle
Breast vHMEC Mammary Epithelial
Fetal Lung
Colon Smooth Muscle
Fetal Stomach
Left Ventricle
Endometrial Stromal 1
CD19 − Primary B cells (PB)
Thyroid
Primary Th naive cells (PB)
Adipose Derived MSC Cultured Cells
Primary T killer naive cells (PB)
Fetal Kidney
Breast Stromal
Fetal Adrenal Gland
Primary T cells (CB)
CD34 − Primary HSC
Primary Th memory cells (PB)
Primary Th cells (PB)
Fetal Heart
Fetal Muscle Leg
Primary Th memory cells (PB) 1
Fetal Muscle Trunk
Primary Th 17 cells PMA−I stimulated
Primary Natural Killer cells (PB)
Primary Th cells PMA−I stimulated
Primary T cells effector/memory enriched (PB)
Thymus
Primary monocytes (PB)
Primary Th naive cells (PB)
Primary T cells (PB)
Primary Treg cells (PB)
Primary T killer memory cells (PB)
CD19 − Primary B cells (CB)
CD34 − Primary HSC G−CSF−mobilized Male
Primary neutrophils (PB)
CD34 − Primary HSC short term culture
Brain Hippocampus Middle
CD34 − Primary HSC G−CSF−mobilized Female
Brain Substantia Nigra
Fetal Brain Male
Pancreatic Islets
Brain Dorsolateral Prefrontal Cortex
Brain Anterior Caudate
Cortex derived neurospheres
Ganglion Eminence derived neurospheres
Brain Cingulate Gyrus
Brain Inferior Temporal Lobe
Brain Angular Gyrus
Fetal Brain Female
Brain Germinal Matrix
Epigenomes of tissues used for prediction of mutation landscape

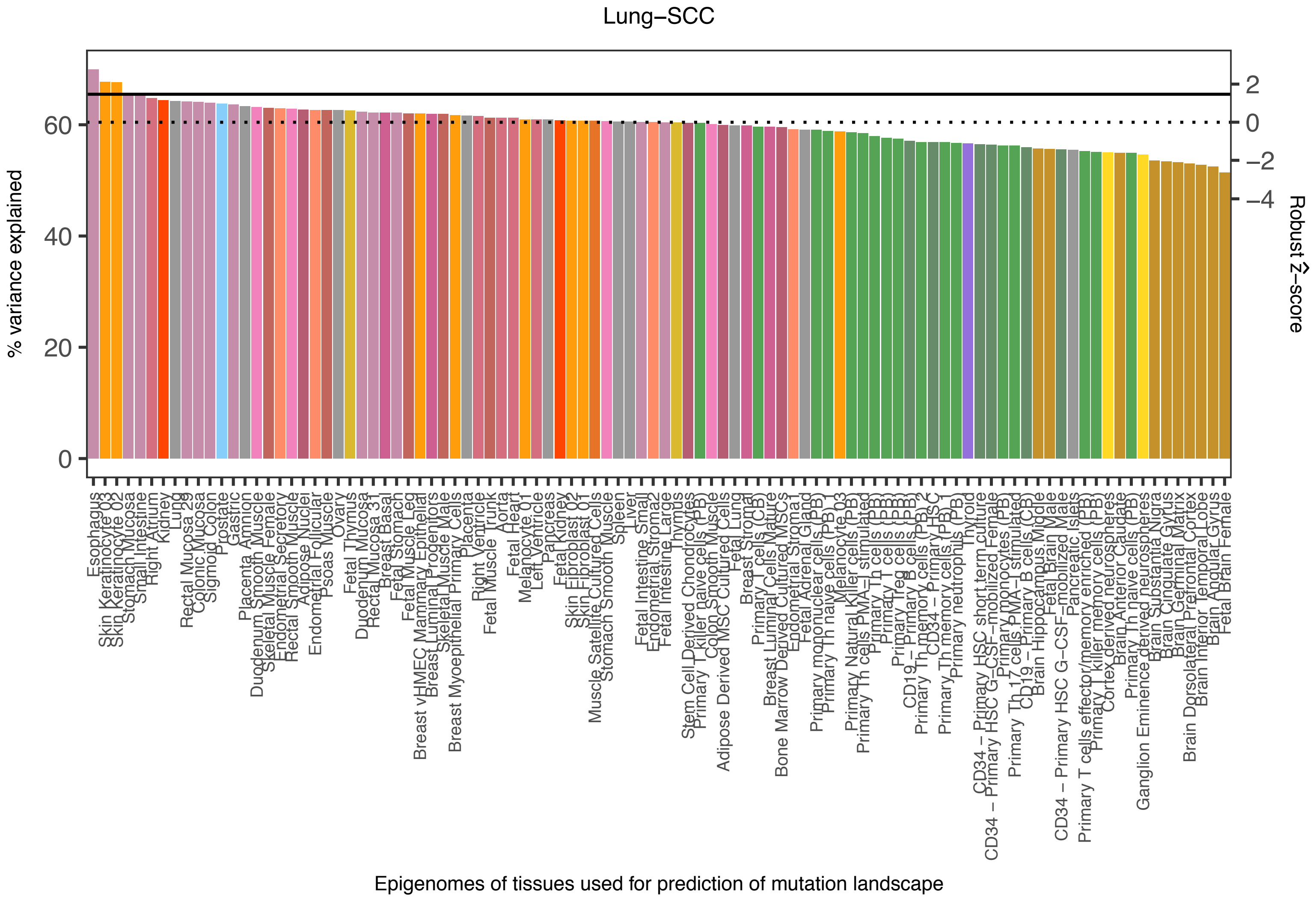

Lung−SCC
% variance explained
Robust Ẑ−score
0
20
40
60
2
0
−2
−4
Esophagus
Skin Keratinocyte 03
Skin Keratinocyte 02
Stomach Mucosa
Small Intestine
Right Atrium
Kidney
Lung
Rectal Mucosa 29
Colonic Mucosa
Sigmoid Colon
Prostate
Gastric
Placenta Amnion
Duodenum Smooth Muscle
Skeletal Muscle Female
Endometrial Secretory
Rectal Smooth Muscle
Adipose Nuclei
Endometrial Follicular
Psoas Muscle
Ovary
Fetal Thymus
Duodenum Mucosa
Rectal Mucosa 31
Breast Basal
Fetal Stomach
Fetal Muscle Leg
Breast vHMEC Mammary Epithelial
Breast Luminal Progenitors
Skeletal Muscle Male
Breast Myoepithelial Primary Cells
Placenta
Right Ventricle
Fetal Muscle Trunk
Aorta
Fetal Heart
Melanocyte 01
Left Ventricle
Pancreas
Fetal Kidney
Skin Fibroblast 02
Skin Fibroblast 01
Muscle Satellite Cultured Cells
Stomach Smooth Muscle
Spleen
Liver
Fetal Intestine Small
Endometrial Stromal
Fetal Intestine Large
Thymus
Stem Cell Derived Chondrocytes
Primary T killer naive cells (PB)
Colon Smooth Muscle
Adipose Derived MSC Cultured Cells
Fetal Lung
Breast Stromal
Primary T cells (CB)
Breast Luminal Cells Mature
Bone Marrow Derived Cultured MSCs
Endometrial Stromal
Fetal Adrenal Gland
Primary mononuclear cells (PB)
Primary Th naive cells (PB)
Melanocyte 03
Primary Natural Killer cells (PB)
Primary Th cells PMA−I stimulated
Primary Th cells (PB)
Primary T cells (PB)
Primary Treg cells (PB)
CD19 − Primary B cells (PB)
Primary Th memory cells (PB)
CD34 − Primary HSC
Primary Th memory cells (PB)
Primary neutrophils (PB)
Thyroid
CD34 − Primary HSC short term culture
CD34 − Primary HSC G−CSF−mobilized Female
Primary monocytes (PB)
Primary Th17 cells PMA−I stimulated
CD19 − Primary B cells (CB)
Brain Hippocampus Middle
Fetal Brain Male
CD34 − Primary HSC G−CSF−mobilized Male
Pancreatic Islets
Primary T cells effector/memory enriched (PB)
Primary T killer memory cells (PB)
Cortex derived neurospheres
Brain Anterior Caudate
Primary Th naive cells (PB)
Ganglion Eminence derived neurospheres
Brain Substantia Nigra
Brain Cingulate Gyrus
Brain Germinal Matrix
Brain Dorsolateral Prefrontal Cortex
Brain Inferior Temporal Lobe
Brain Angular Gyrus
Fetal Brain Female
Epigenomes of tissues used for prediction of mutation landscape

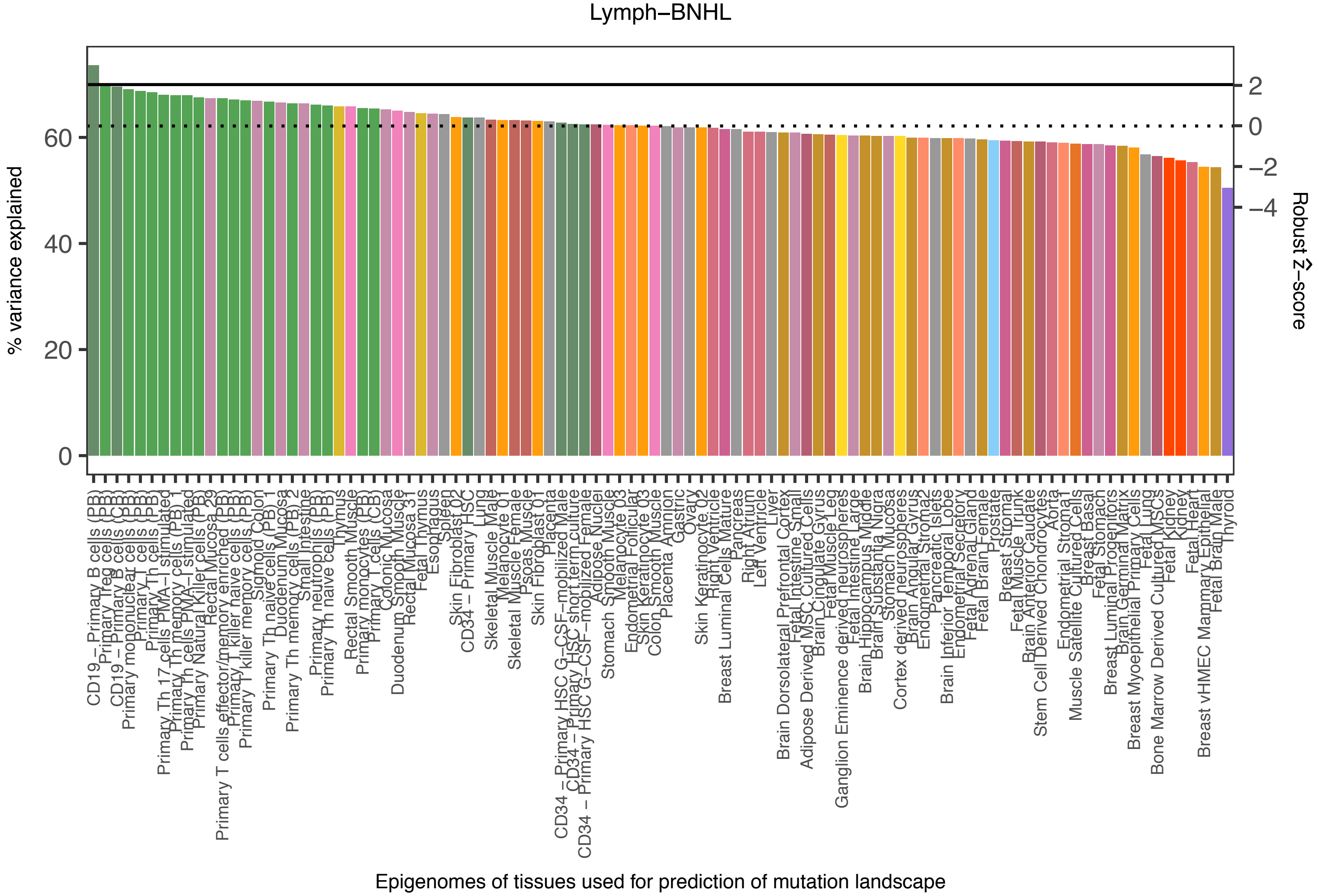
Lymph−BNHL
% variance explained
0
20
40
60
Robust Ẑ−score
2
0
−2
−4
Epigenomes of tissues used for prediction of mutation landscape

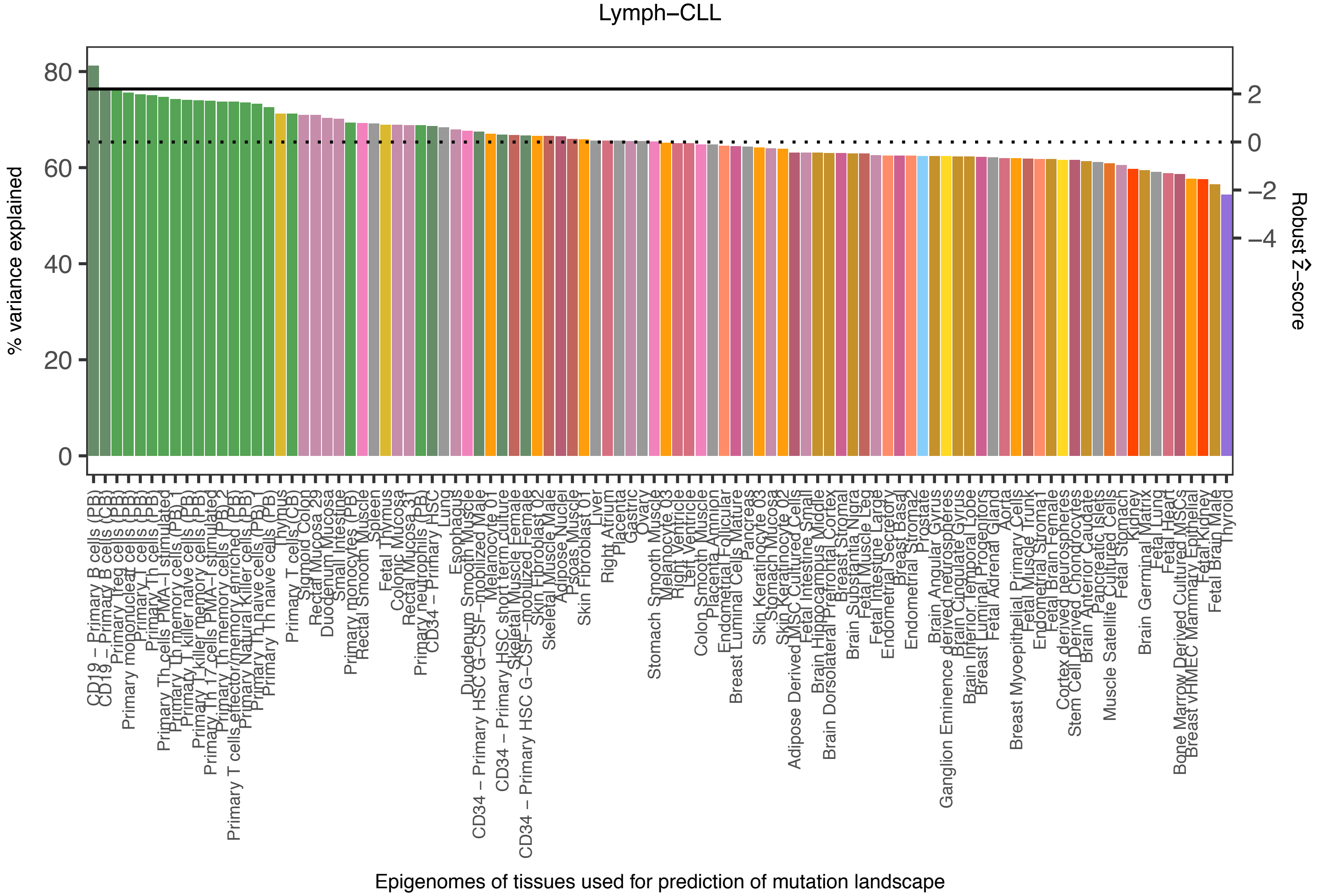
Lymph−CLL
% variance explained
0
20
40
60
80
Robust Ẑ−score
2
0
−2
−4
Epigenomes of tissues used for prediction of mutation landscape
CD19 − Primary B cells (PB)
CD19 − Primary B cells (CB)
Primary Treg cells (PB)
Primary mononuclear cells (PB)
Primary T cells (PB)
Primary Th cells (PB)
Primary Th cells PMA−I stimulated
Primary Th memory cells (PB) 1
Primary T killer naive cells (PB)
Primary T killer memory cells (PB)
Primary Th 17 cells PMA−I stimulated
Primary Th memory cells (PB) 2
Primary T cells effector/memory enriched (PB)
Primary Natural Killer cells (PB)
Primary Th naive cells (PB)
Primary Th naive cells (PB)
Thymus
Primary T cells (CB)
Sigmoid Colon
Rectal Mucosa 29
Duodenum Mucosa
Small Intestine
Primary monocytes (PB)
Rectal Smooth Muscle
Spleen
Fetal Thymus
Colonic Mucosa
Rectal Mucosa 31
Primary neutrophils (PB)
CD34 − Primary HSC
Lung
Esophagus
Duodenum Smooth Muscle
CD34 − Primary HSC G−CSF−mobilized Male
Melanocyte 01
CD34 − Primary HSC short term culture
Skeletal Muscle Female
CD34 − Primary HSC G−CSF−mobilized Female
Skin Fibroblast 02
Skeletal Muscle Male
Adipose Nuclei
Psoas Muscle
Skin Fibroblast 01
Liver
Right Atrium
Placenta
Gastric
Ovary
Stomach Smooth Muscle
Melanocyte 03
Right Ventricle
Left Ventricle
Colon Smooth Muscle
Placenta Amnion
Endometrial Follicular
Breast Luminal Cells Mature
Pancreas
Skin Keratinocyte 03
Stomach Mucosa
Skin Keratinocyte 02
Adipose Derived MSC Cultured Cells
Fetal Intestine Small
Brain Hippocampus Middle
Brain Dorsolateral Prefrontal Cortex
Breast Stromal
Brain Substantia Nigra
Fetal Muscle Leg
Fetal Intestine Large
Endometrial Secretory
Breast Basal
Endometrial Stroma2
Prostate
Brain Angular Gyrus
Ganglion Eminence derived neurospheres
Brain Cingulate Gyrus
Brain Inferior Temporal Lobe
Breast Luminal Progenitors
Fetal Adrenal Gland
Aorta
Breast Myoepithelial Primary Cells
Fetal Muscle Trunk
Endometrial Stroma1
Fetal Brain Female
Cortex derived neurospheres
Stem Cell Derived Chondrocytes
Brain Anterior Caudate
Pancreatic Islets
Muscle Satellite Cultured Cells
Fetal Stomach
Kidney
Brain Germinal Matrix
Fetal Lung
Fetal Heart
Bone Marrow Derived Cultured MSCs
Breast vHMEC Mammary Epithelial
Fetal Kidney
Fetal Brain Male
Thyroid

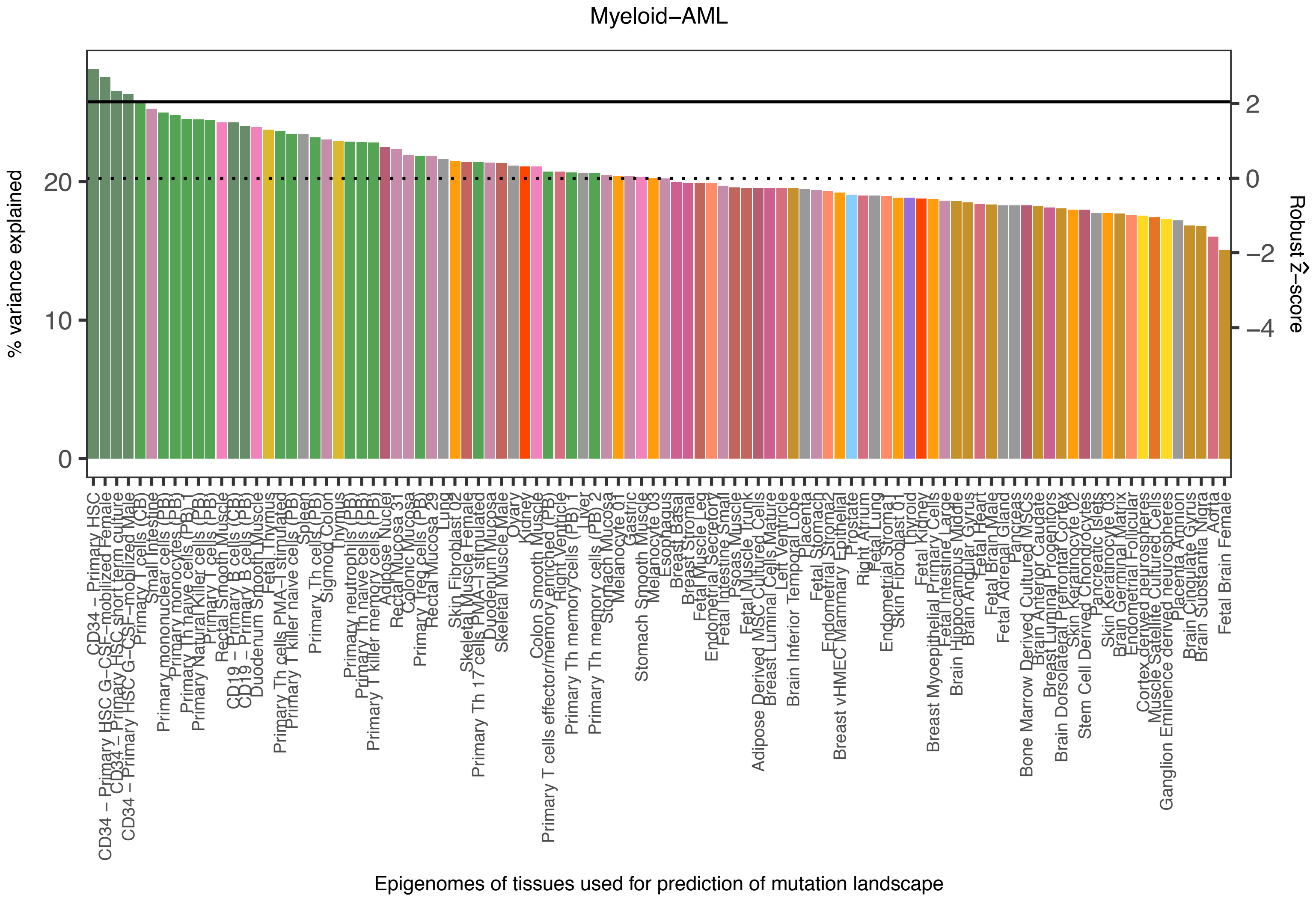
Myeloid−AML
% variance explained
0
10
20
Robust Ẑ−score
2
0
−2
−4
CD34 − Primary HSC
CD34 − Primary HSC G−CSF−mobilized Female
CD34 − Primary HSC short term culture
CD34 − Primary HSC G−CSF−mobilized Male
Primary T cells (CB)
Small Intestine
Primary mononuclear cells (PB)
Primary monocytes (PB)
Primary Th naive cells (PB)
Primary Natural Killer cells (PB)
Primary T cells (PB)
Rectal Smooth Muscle
CD19 − Primary B cells (CB)
CD19 − Primary B cells (PB)
Duodenum Smooth Muscle
Fetal Thymus
Primary Th cells PMA−I stimulated
Primary T killer naive cells (PB)
Spleen
Primary Th cells (PB)
Sigmoid Colon
Thymus
Primary neutrophils (PB)
Primary Th naive cells (PB)
Primary T killer memory cells (PB)
Adipose Nuclei
Rectal Mucosa 31
Colonic Mucosa
Primary Treg cells (PB)
Rectal Mucosa 29
Lung
Skin Fibroblast 02
Skeletal Muscle Female
Primary Th 17 cells PMA−I stimulated
Duodenum Mucosa
Skeletal Muscle Male
Ovary
Kidney
Colon Smooth Muscle
Primary T cells effector/memory enriched (PB)
Right Ventricle
Primary Th memory cells (PB) 1
Liver
Primary Th memory cells (PB) 2
Stomach Mucosa
Melanocyte 01
Gastric
Stomach Smooth Muscle
Melanocyte 03
Esophagus
Breast Basal
Breast Stromal
Fetal Muscle Leg
Endometrial Secretory
Fetal Intestine Small
Psoas Muscle
Fetal Muscle Trunk
Adipose Derived MSC Cultured Cells
Breast Luminal Cells Mature
Left Ventricle
Brain Inferior Temporal Lobe
Placenta
Fetal Stomach
Endometrial Stromal2
Breast vHMEC Mammary Epithelial
Prostate
Right Atrium
Fetal Lung
Endometrial Stromal
Skin Fibroblast 01
Thyroid
Fetal Kidney
Breast Myoepithelial Primary Cells
Fetal Intestine Large
Brain Hippocampus Middle
Brain Angular Gyrus
Fetal Heart
Fetal Brain Male
Fetal Adrenal Gland
Pancreas
Bone Marrow Derived Cultured MSCs
Brain Anterior Caudate
Breast Luminal Progenitors
Brain Dorsolateral Prefrontal Cortex
Skin Keratinocyte 02
Stem Cell Derived Chondrocytes
Pancreatic Islets
Skin Keratinocyte 03
Brain Germinal Matrix
Endometrial Follicular
Cortex derived neurospheres
Muscle Satellite Cultured Cells
Ganglion Eminence derived neurospheres
Placenta Amnion
Brain Cingulate Gyrus
Brain Substantia Nigra
Aorta
Fetal Brain Female
Epigenomes of tissues used for prediction of mutation landscape

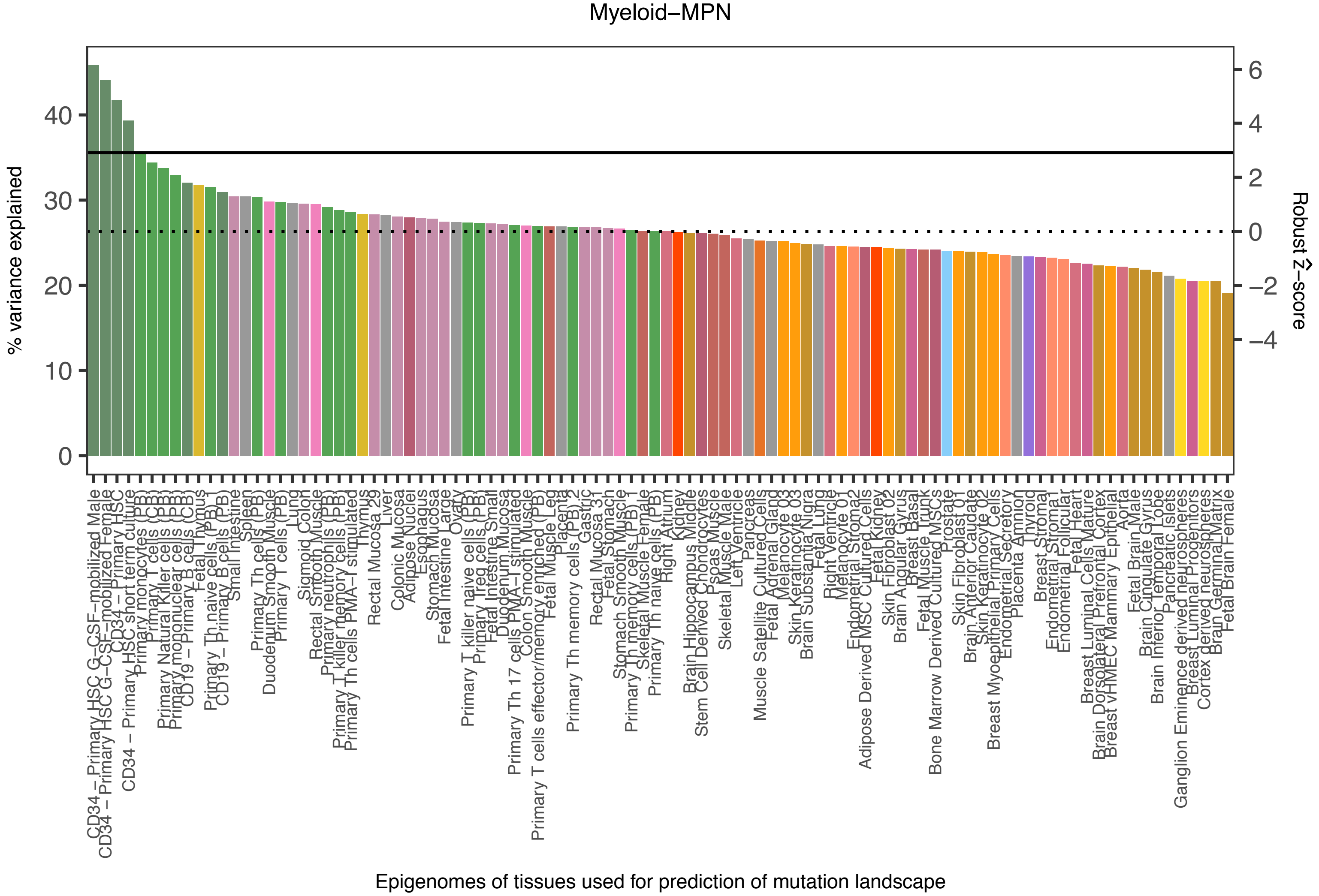

Myeloid–MPN
% variance explained
0
10
20
30
40
Robust Ẑ–score
6
4
2
0
−2
−4
CD34 – Primary HSC G–CSF–mobilized Male
CD34 – Primary HSC G–CSF–mobilized Female
CD34 – Primary HSC
CD34 – Primary HSC short term culture
Primary monocytes (PB)
Primary T cells (CB)
Primary Natural Killer cells (PB)
Primary mononuclear cells (PB)
CD19 – Primary B cells (CB)
Fetal Thymus
Primary Th naive cells (PB)
CD19 – Primary B cells (PB)
Small Intestine
Spleen
Primary Th cells (PB)
Duodenum Smooth Muscle
Primary T cells (PB)
Lung
Sigmoid Colon
Rectal Smooth Muscle
Primary neutrophils (PB)
Primary T killer memory cells (PB)
Primary Th cells PMA–I stimulated
Thymus
Rectal Mucosa 29
Liver
Colonic Mucosa
Adipose Nuclei
Esophagus
Stomach Mucosa
Fetal Intestine Large
Ovary
Primary T killer naive cells (PB)
Primary Treg cells (PB)
Fetal Intestine Small
Duodenum Mucosa
Primary Th 17 cells PMA–I stimulated
Colon Smooth Muscle
Primary T cells effector/memory enriched (PB)
Fetal Muscle Leg
Placenta
Primary Th memory cells (PB) 2
Gastric
Rectal Mucosa 31
Fetal Stomach
Stomach Smooth Muscle
Primary Th memory cells (PB) 1
Skeletal Muscle Female
Primary Th naive cells (PB)
Right Atrium
Kidney
Brain Hippocampus Middle
Stem Cell Derived Chondrocytes
Psoas Muscle
Skeletal Muscle Male
Left Ventricle
Pancreas
Muscle Satellite Cultured Cells
Fetal Adrenal Gland
Melanocyte 03
Skin Keratinocyte 03
Brain Substantia Nigra
Fetal Lung
Right Ventricle
Melanocyte 01
Endometrial Stroma2
Adipose Derived MSC Cultured Cells
Fetal Kidney
Skin Fibroblast 02
Brain Angular Gyrus
Breast Basal
Fetal Muscle Trunk
Bone Marrow Derived Cultured MSCs
Prostate
Skin Fibroblast 01
Brain Anterior Caudate
Skin Keratinocyte 02
Breast Myoepithelial Primary Cells
Endometrial Secretory
Placenta Amnion
Thyroid
Breast Stromal
Endometrial Stroma1
Endometrial Follicular
Fetal Heart
Breast Luminal Cells Mature
Brain Dorsolateral Prefrontal Cortex
Breast vHMEC Mammary Epithelial
Aorta
Fetal Brain Male
Brain Cingulate Gyrus
Brain Inferior Temporal Lobe
Pancreatic Islets
Ganglion Eminence derived neurospheres
Breast Luminal Progenitors
Cortex derived neurospheres
Brain Germinal Matrix
Fetal Brain Female
Epigenomes of tissues used for prediction of mutation landscape

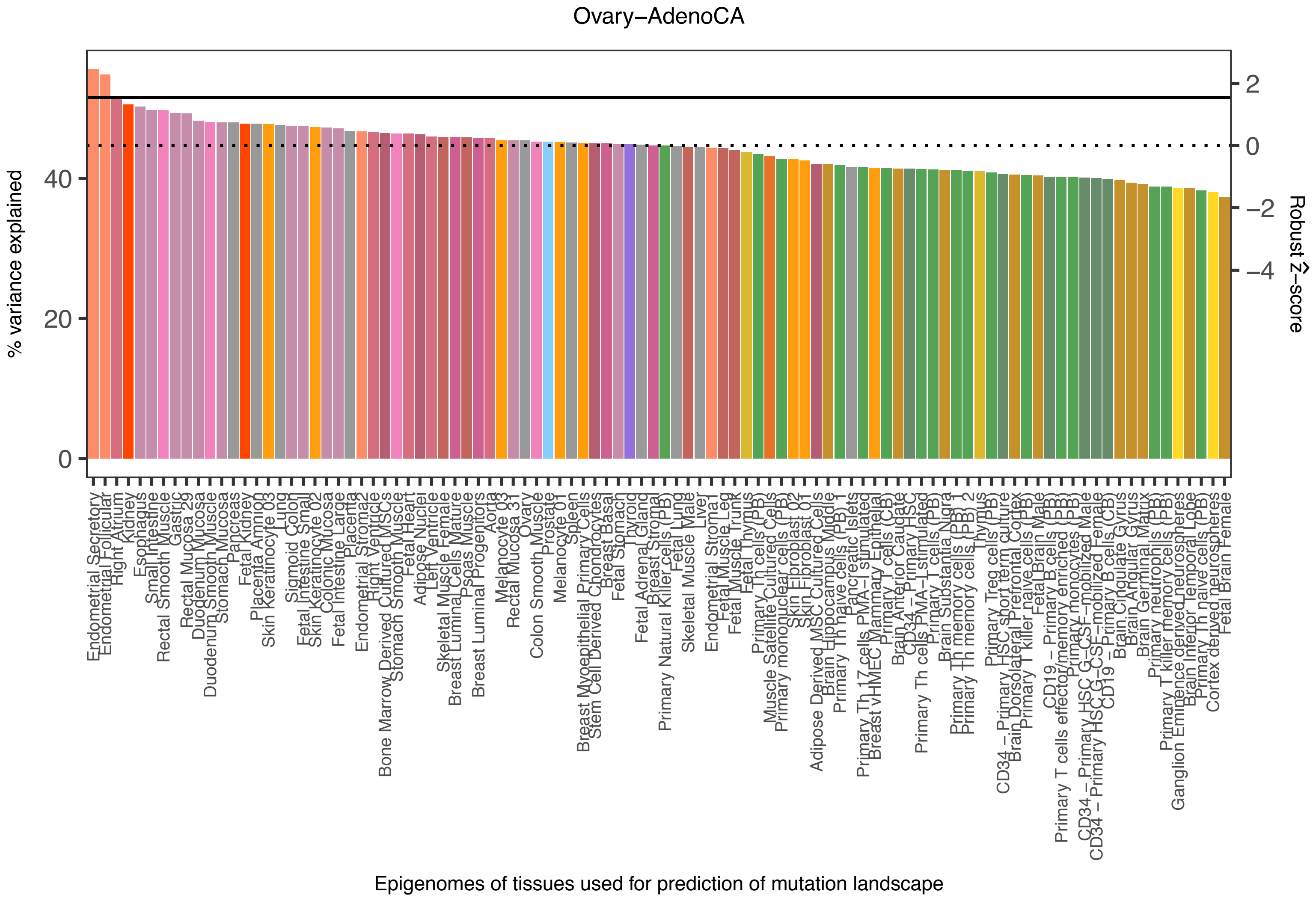
Ovary−AdenoCA
% variance explained
0
20
40
Robust Ẑ−score
2
0
−2
−4
Endometrial Secretory
Endometrial Follicular
Right Atrium
Kidney
Esophagus
Small Intestine
Rectal Smooth Muscle
Gastric
Rectal Mucosa 29
Duodenum Mucosa
Duodenum Smooth Muscle
Stomach Mucosa
Pancreas
Fetal Kidney
Placenta Amnion
Skin Keratinocyte 03
Lung
Sigmoid Colon
Fetal Intestine Small
Skin Keratinocyte 02
Colonic Mucosa
Fetal Intestine Large
Placenta
Endometrial Stromal
Right Ventricle
Bone Marrow Derived Cultured MSCs
Stomach Smooth Muscle
Fetal Heart
Adipose Nuclei
Left Ventricle
Skeletal Muscle Female
Breast Luminal Cells Mature
Psoas Muscle
Breast Luminal Progenitors
Aorta
Melanocyte 03
Rectal Mucosa 31
Ovary
Colon Smooth Muscle
Prostate
Melanocyte 01
Spleen
Breast Myoepithelial Primary Cells
Stem Cell Derived Chondrocytes
Breast Basal
Fetal Stomach
Thyroid
Fetal Adrenal Gland
Breast Stromal
Primary Natural Killer cells (PB)
Fetal Lung
Skeletal Muscle Male
Liver
Endometrial Stromal
Fetal Muscle Leg
Fetal Muscle Trunk
Fetal Thymus
Primary Th cells (PB)
Muscle Satellite Cultured Cells
Primary mononuclear cells (PB)
Skin Fibroblast 02
Skin Fibroblast 01
Adipose Derived MSC Cultured Cells
Brain Hippocampus Middle
Primary Th naive cells (PB)
Pancreatic Islets
Primary Th17 cells PMA−I stimulated
Breast vHMEC Mammary Epithelial
Primary T cells (CB)
Brain Anterior Caudate
CD34 − Primary HSC
Primary Th cells PMA−I stimulated
Primary T cells (PB)
Brain Substantia Nigra
Primary Th memory cells (PB) 1
Primary Th memory cells (PB) 2
Thymus
Primary Treg cells (PB)
CD34 − Primary HSC short term culture
Brain Dorsolateral Prefrontal Cortex
Primary T killer naive cells (PB)
Fetal Brain Male
CD19 − Primary B cells (PB)
Primary T cells effector/memory enriched (PB)
Primary monocytes (PB)
CD34 − Primary HSC G−CSF−mobilized Male
CD34 − Primary HSC G−CSF−mobilized Female
CD19 − Primary B cells (CB)
Brain Cingulate Gyrus
Brain Angular Gyrus
Brain Germinal Matrix
Primary neutrophils (PB)
Primary T killer memory cells (PB)
Ganglion Eminence derived neurospheres
Brain Inferior Temporal Lobe
Primary Th naive cells (PB)
Cortex derived neurospheres
Fetal Brain Female
Epigenomes of tissues used for prediction of mutation landscape

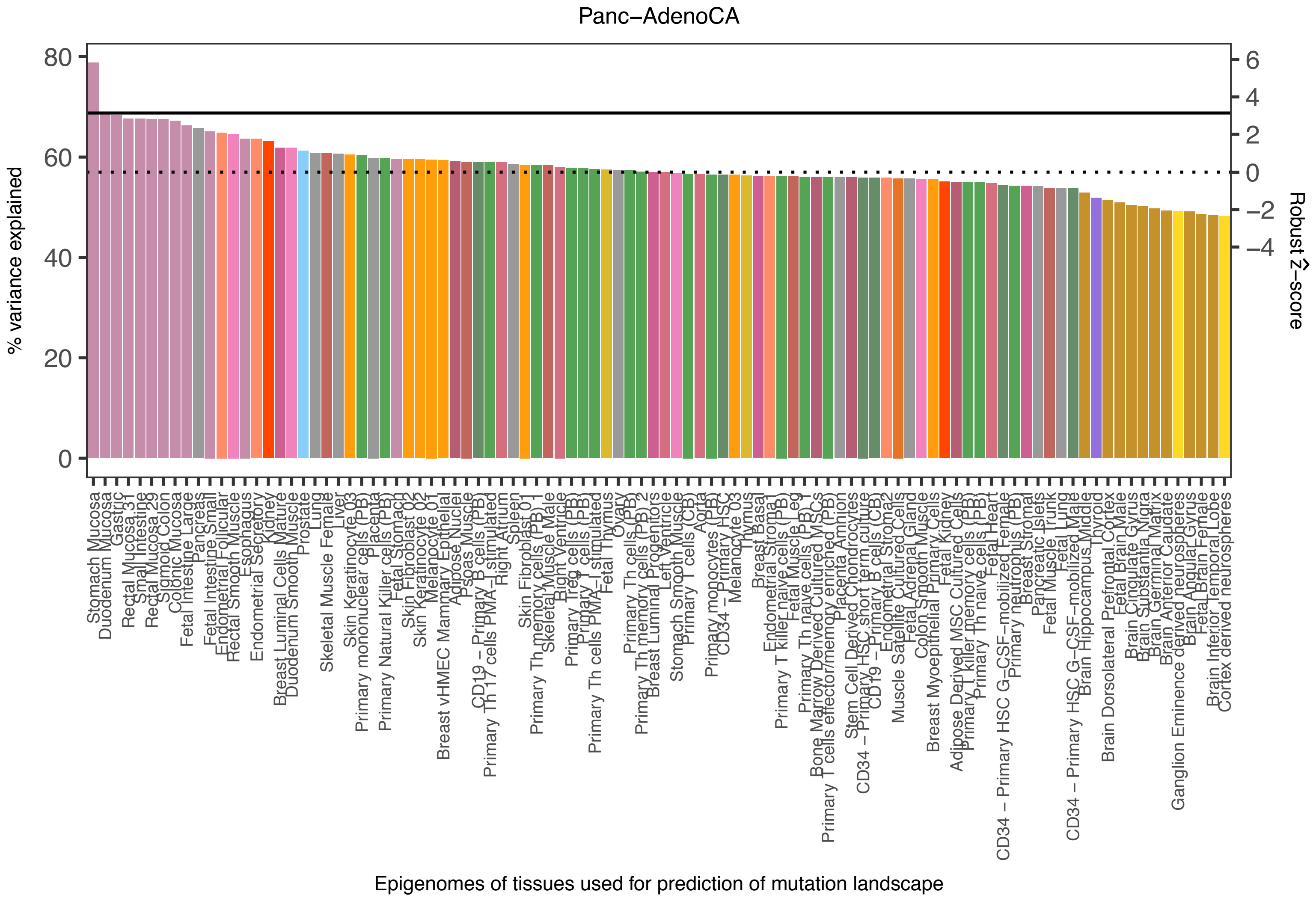

Panc−AdenoCA
% variance explained
0
20
40
60
80
Robust Ẑ−score
6
4
2
0
−2
−4
Epigenomes of tissues used for prediction of mutation landscape
Stomach Mucosa
Duodenum Mucosa
Gastric
Rectal Mucosa 31
Small Intestine
Rectal Mucosa 29
Sigmoid Colon
Colonic Mucosa
Fetal Intestine Large
Pancreas
Fetal Intestine Small
Endometrial Follicular
Rectal Smooth Muscle
Esophagus
Endometrial Secretory
Kidney
Breast Luminal Cells Mature
Duodenum Smooth Muscle
Prostate
Lung
Skeletal Muscle Female
Liver
Skin Keratinocyte 03
Primary mononuclear cells (PB)
Placenta
Primary Natural Killer cells (PB)
Fetal Stomach
Skin Fibroblast 02
Skin Keratinocyte 02
Melanocyte 01
Breast vHMEC Mammary Epithelial
Adipose Nuclei
Psoas Muscle
CD19 − Primary B cells (PB)
Primary Th17 cells PMA−I stimulated
Right Atrium
Spleen
Skin Fibroblast 01
Primary Th memory cells (PB) 1
Skeletal Muscle Male
Right Ventricle
Primary Treg cells (PB)
Primary T cells (PB)
Primary Th cells PMA−I stimulated
Fetal Thymus
Ovary
Primary Th cells (PB)
Primary Th memory cells (PB) 2
Breast Luminal Progenitors
Left Ventricle
Stomach Smooth Muscle
Primary T cells (CB)
Aorta
Primary monocytes (PB)
CD34 − Primary HSC
Melanocyte 03
Thymus
Breast Basal
Endometrial Stromal
Primary T killer naive cells (PB)
Fetal Muscle Leg
Primary Th naive cells (PB) 1
Bone Marrow Derived Cultured MSCs
Primary T cells effector/memory enriched (PB)
Placenta Amnion
Stem Cell Derived Chondrocytes
CD34 − Primary HSC short term culture
CD19 − Primary B cells (CB)
Endometrial Stroma2
Muscle Satellite Cultured Cells
Fetal Adrenal Gland
Colon Smooth Muscle
Breast Myoepithelial Primary Cells
Fetal Kidney
Adipose Derived MSC Cultured Cells
Primary T killer memory cells (PB)
Primary Th naive cells (PB) 2
Fetal Heart
CD34 − Primary HSC G−CSF−mobilized Female
Primary neutrophils (PB)
Breast Stromal
Pancreatic Islets
Fetal Muscle Trunk
Fetal Lung
CD34 − Primary HSC G−CSF−mobilized Male
Brain Hippocampus Middle
Thyroid
Brain Dorsolateral Prefrontal Cortex
Fetal Brain Male
Brain Cingulate Gyrus
Brain Substantia Nigra
Brain Germinal Matrix
Brain Anterior Caudate
Ganglion Eminence derived neurospheres
Brain Angular Gyrus
Fetal Brain Female
Brain Inferior Temporal Lobe
Cortex derived neurospheres

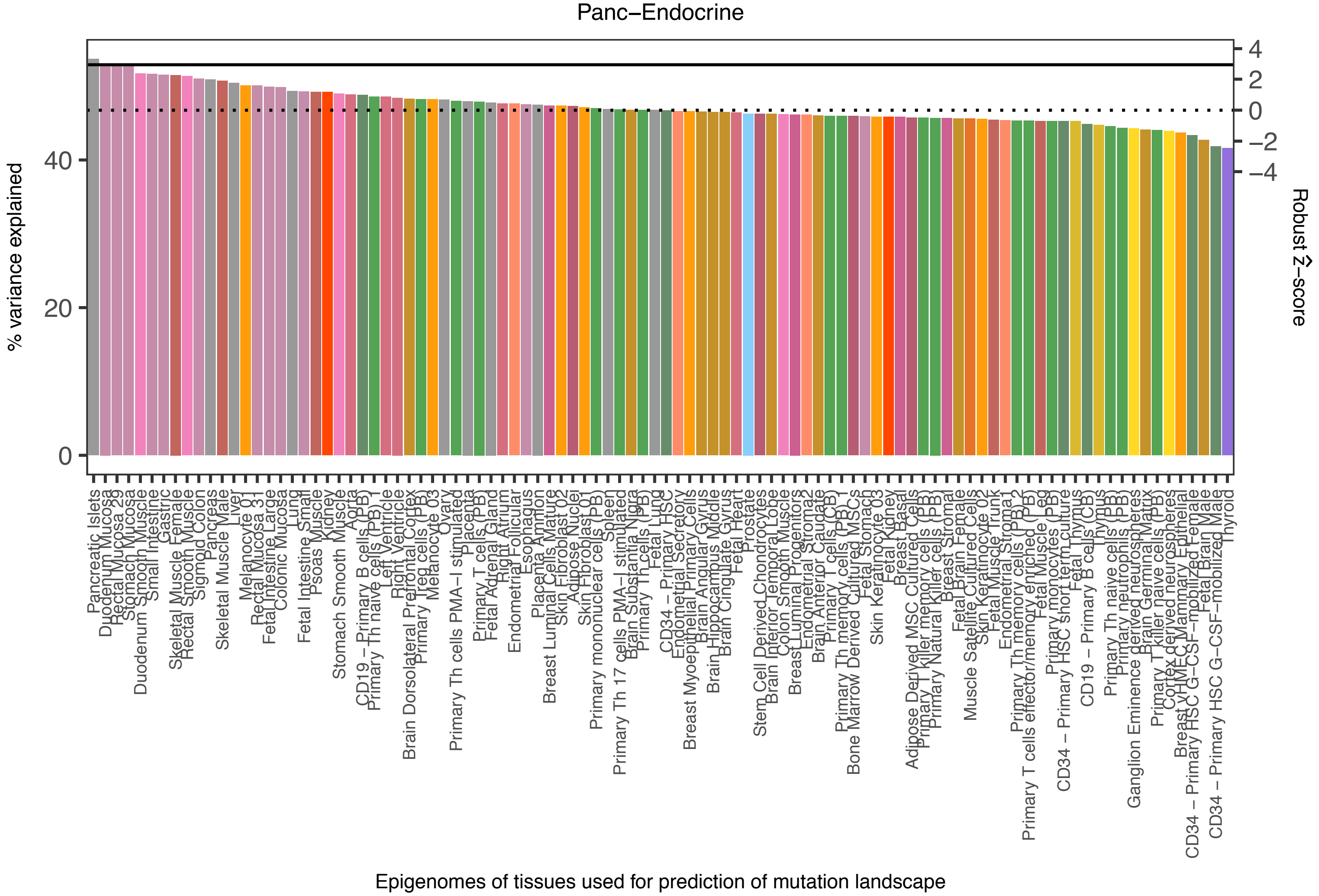

Panc−Endocrine
% variance explained
Robust Ẑ−score
Epigenomes of tissues used for prediction of mutation landscape
Pancreatic Islets
Thyroid

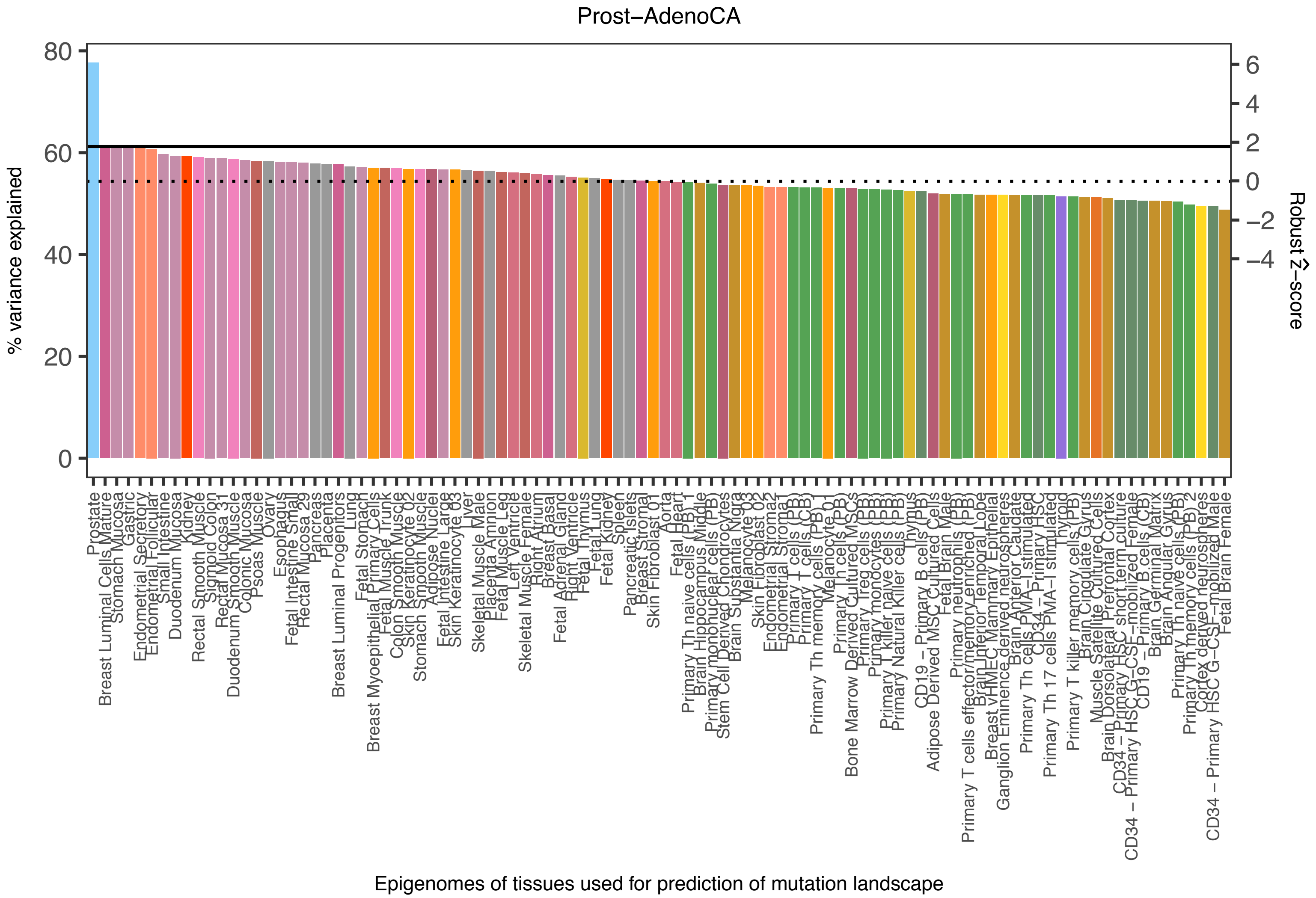

Prost−AdenoCA
% variance explained
0
20
40
60
80
Robust Ẑ−score
−4
−2
0
2
4
6
Prostate
Breast Luminal Cells Mature
Stomach Mucosa
Gastric
Endometrial Secretory
Endometrial Follicular
Small Intestine
Duodenum Mucosa
Kidney
Rectal Smooth Muscle
Sigmoid Colon
Rectal Mucosa 31
Duodenum Smooth Muscle
Colonic Mucosa
Psoas Muscle
Ovary
Esophagus
Fetal Intestine Small
Rectal Mucosa 29
Pancreas
Placenta
Breast Luminal Progenitors
Lung
Fetal Stomach
Breast Myoepithelial Primary Cells
Fetal Muscle Trunk
Colon Smooth Muscle
Skin Keratinocyte 02
Stomach Smooth Muscle
Adipose Nuclei
Fetal Intestine Large
Skin Keratinocyte 03
Liver
Skeletal Muscle Male
Placenta Amnion
Fetal Muscle Leg
Left Ventricle
Skeletal Muscle Female
Right Atrium
Breast Basal
Fetal Adrenal Gland
Right Ventricle
Fetal Thymus
Fetal Lung
Fetal Kidney
Spleen
Pancreatic Islets
Breast Stromal
Skin Fibroblast 01
Aorta
Fetal Heart
Primary Th naive cells (PB) 1
Brain Hippocampus Middle
Primary mononuclear cells (PB)
Stem Cell Derived Chondrocytes
Brain Substantia Nigra
Melanocyte 03
Skin Fibroblast 02
Endometrial Stroma2
Endometrial Stroma1
Primary T cells (CB)
Primary T cells (PB)
Primary Th memory cells (PB) 1
Melanocyte 01
Primary Th cells (PB)
Bone Marrow Derived Cultured MSCs
Primary Treg cells (PB)
Primary monocytes (PB)
Primary T killer naive cells (PB)
Primary Natural Killer cells (PB)
Thymus
CD19 − Primary B cells (PB)
Adipose Derived MSC Cultured Cells
Fetal Brain Male
Primary neutrophils (PB)
Primary T cells effector/memory enriched (PB)
Brain Inferior Temporal Lobe
Breast vHMEC Mammary Epithelial
Ganglion Eminence derived neurospheres
Brain Anterior Caudate
Primary Th cells PMA−I stimulated
CD34 − Primary HSC
Primary Th 17 cells PMA−I stimulated
Thyroid
Primary T killer memory cells (PB)
Brain Cingulate Gyrus
Muscle Satellite Cultured Cells
Brain Dorsolateral Prefrontal Cortex
CD34 − Primary HSC short term culture
CD34 − Primary HSC G−CSF−mobilized Female
CD19 − Primary B cells (CB)
Brain Germinal Matrix
Brain Angular Gyrus
Primary Th naive cells (PB) 2
Primary Th memory cells (PB) 2
Cortex derived neurospheres
CD34 − Primary HSC G−CSF−mobilized Male
Fetal Brain Female
Epigenomes of tissues used for prediction of mutation landscape

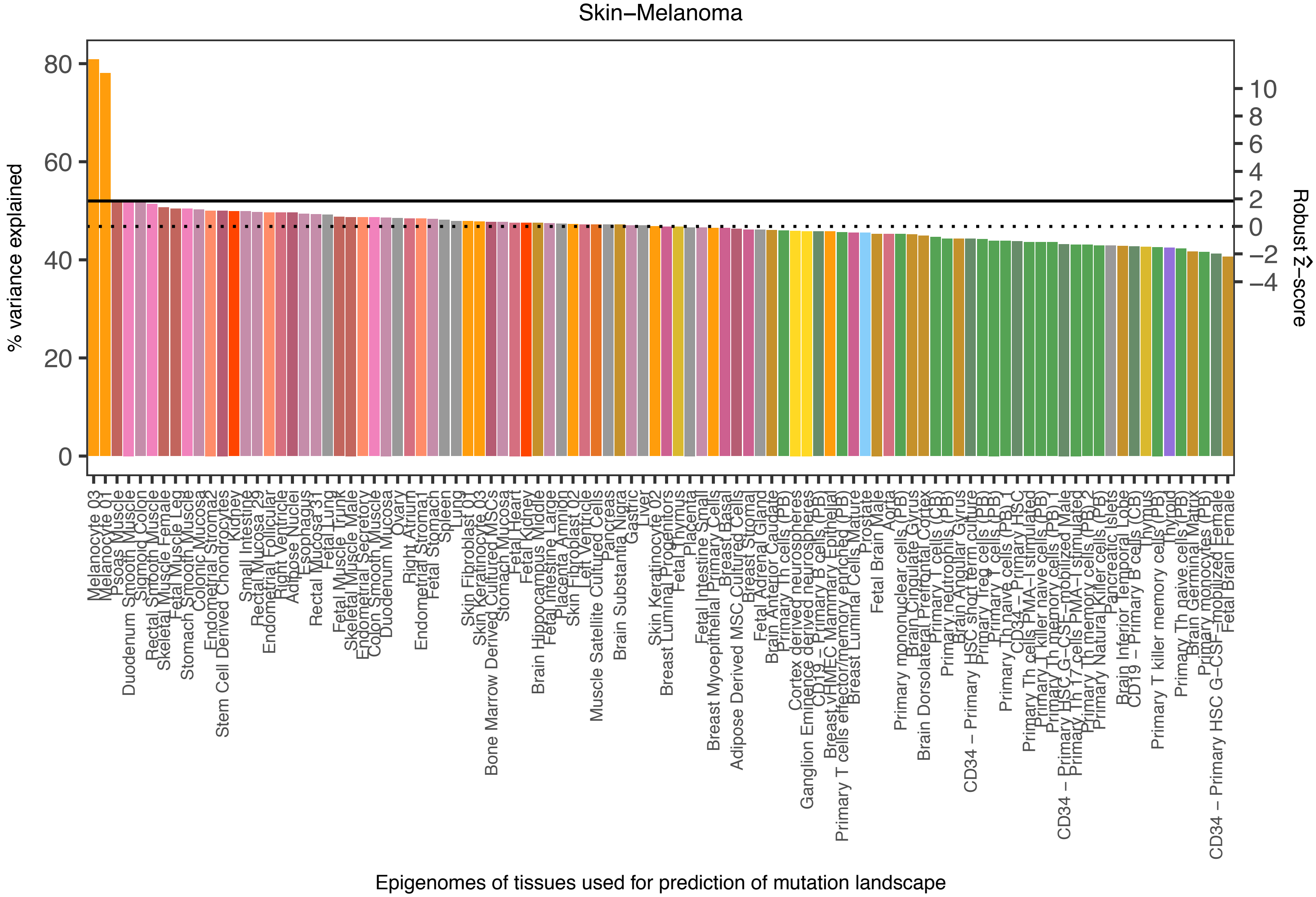
Skin−Melanoma
% variance explained
Robust Z-score
Epigenomes of tissues used for prediction of mutation landscape

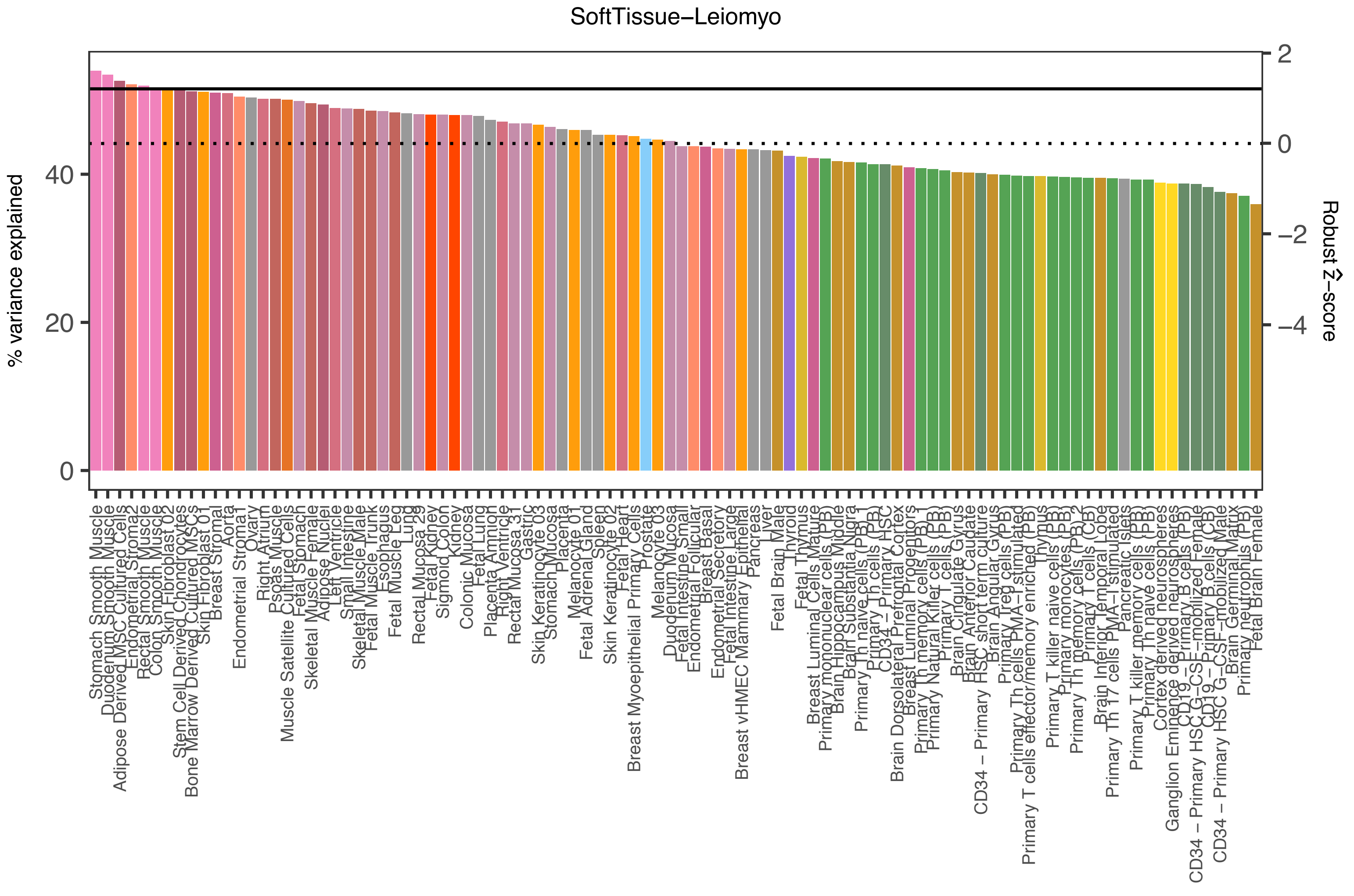

SoftTissue−Leiomyo
% variance explained
0
20
40
Robust Ẑ−score
2
0
−2
−4
Stomach Smooth Muscle
Duodenum Smooth Muscle
Adipose Derived MSC Cultured Cells
Endometrial Stroma2
Rectal Smooth Muscle
Colon Smooth Muscle
Skin Fibroblast 02
Stem Cell Derived Chondrocytes
Bone Marrow Derived Cultured MSCs
Skin Fibroblast 01
Breast Stromal
Aorta
Endometrial Stroma1
Ovary
Right Atrium
Psoas Muscle
Muscle Satellite Cultured Cells
Fetal Stomach
Skeletal Muscle Female
Adipose Nuclei
Left Ventricle
Small Intestine
Skeletal Muscle Male
Fetal Muscle Trunk
Esophagus
Fetal Muscle Leg
Lung
Rectal Mucosa 29
Fetal Kidney
Sigmoid Colon
Kidney
Colonic Mucosa
Fetal Lung
Placenta Amnion
Right Ventricle
Rectal Mucosa 31
Gastric
Skin Keratinocyte 03
Stomach Mucosa
Placenta
Melanocyte 01
Fetal Adrenal Gland
Spleen
Skin Keratinocyte 02
Fetal Heart
Breast Myoepithelial Primary Cells
Prostate
Melanocyte 03
Duodenum Mucosa
Fetal Intestine Small
Endometrial Glandular
Breast Basal
Endometrial Secretory
Fetal Intestine Large
Breast vHMEC Mammary Epithelial
Pancreas
Liver
Fetal Brain Male
Thyroid
Fetal Thymus
Breast Luminal Cells Mature
Primary mononuclear cells (PB)
Brain Hippocampus Middle
Brain Substantia Nigra
Primary Th naive cells (PB)
Primary Th cells (PB)
CD34 − Primary HSC
Brain Dorsolateral Prefrontal Cortex
Breast Luminal Progenitors
Primary Th memory cells (PB) 1
Primary Natural Killer cells (PB)
Primary T cells (PB)
Brain Cingulate Gyrus
Brain Anterior Caudate
CD34 − Primary HSC short term culture
Brain Angular Gyrus
Primary Treg cells (PB)
Primary Th cells PMA−I stimulated
Primary T cells effector/memory enriched (PB)
Thymus
Primary T killer naive cells (PB)
Primary monocytes (PB)
Primary Th memory cells (PB) 2
Primary T cells (CB)
Brain Inferior Temporal Lobe
Primary Th 17 cells PMA−I stimulated
Pancreatic Islets
Primary T killer memory cells (PB)
Primary Th naive cells (PB)
Cortex derived neurospheres
Ganglion Eminence derived neurospheres
CD19 − Primary B cells (PB)
CD34 − Primary HSC G−CSF−mobilized Female
CD19 − Primary B cells (CB)
CD34 − Primary HSC G−CSF−mobilized Male
Brain Germinal Matrix
Primary neutrophils (PB)
Fetal Brain Female
Epigenomes of tissues used for prediction of mutation landscape

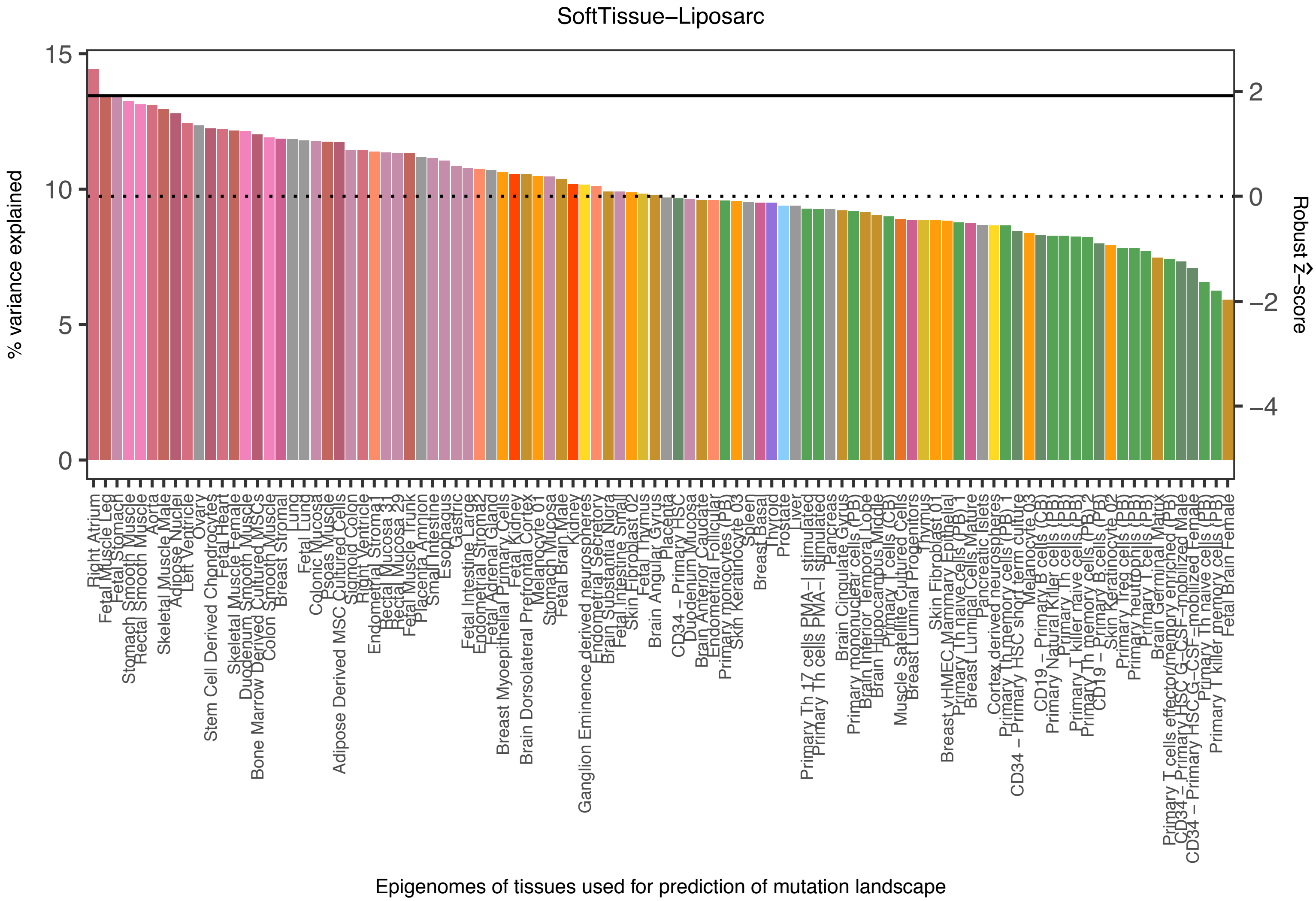
SoftTissue−Liposarc
% variance explained
Robust Ẑ−score
Epigenomes of tissues used for prediction of mutation landscape
15
10
5
0
2
0
−2
−4

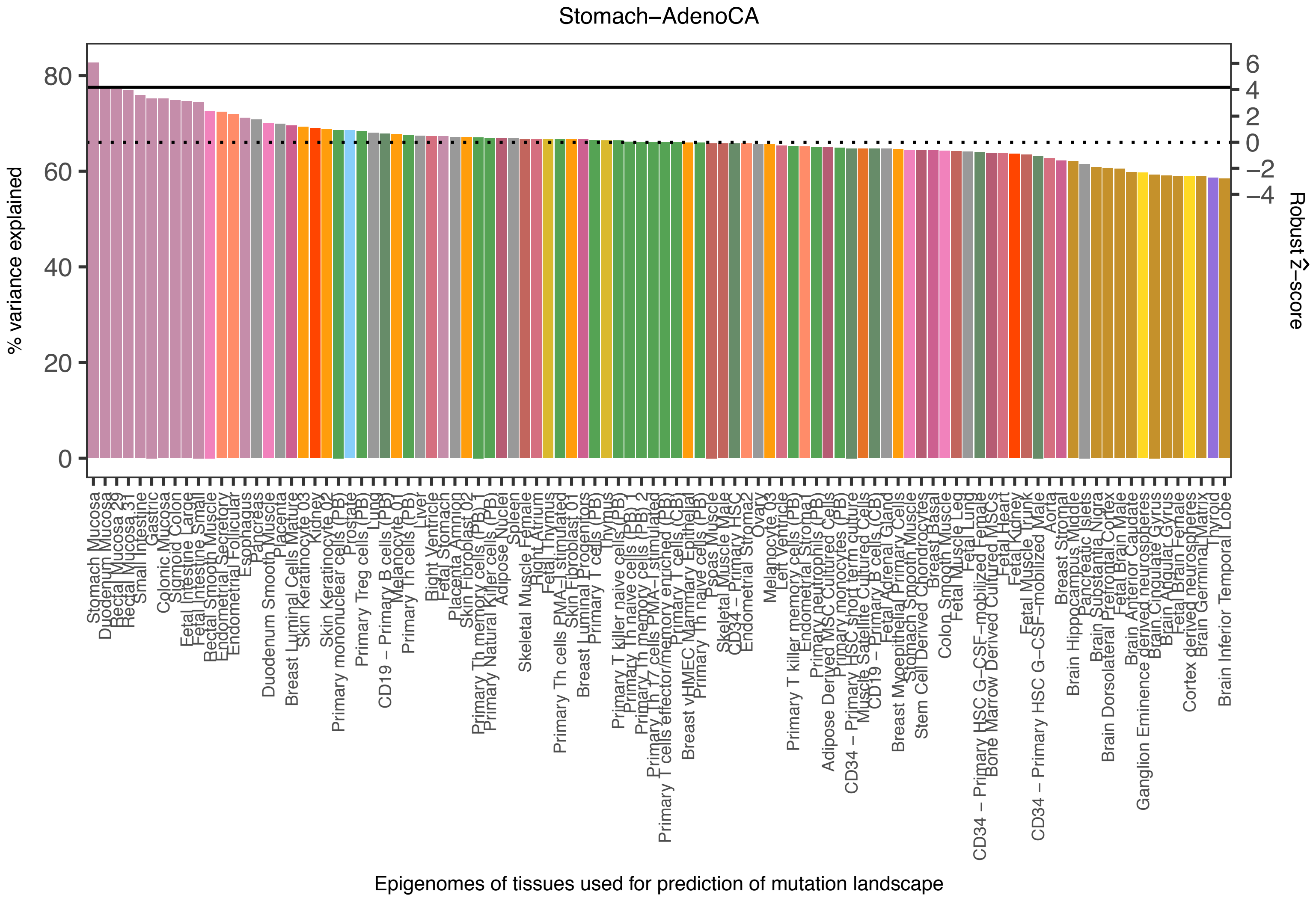
Stomach−AdenoCA
% variance explained
0
20
40
60
80
Robust Ẑ−score
6
4
2
0
−2
−4
Epigenomes of tissues used for prediction of mutation landscape

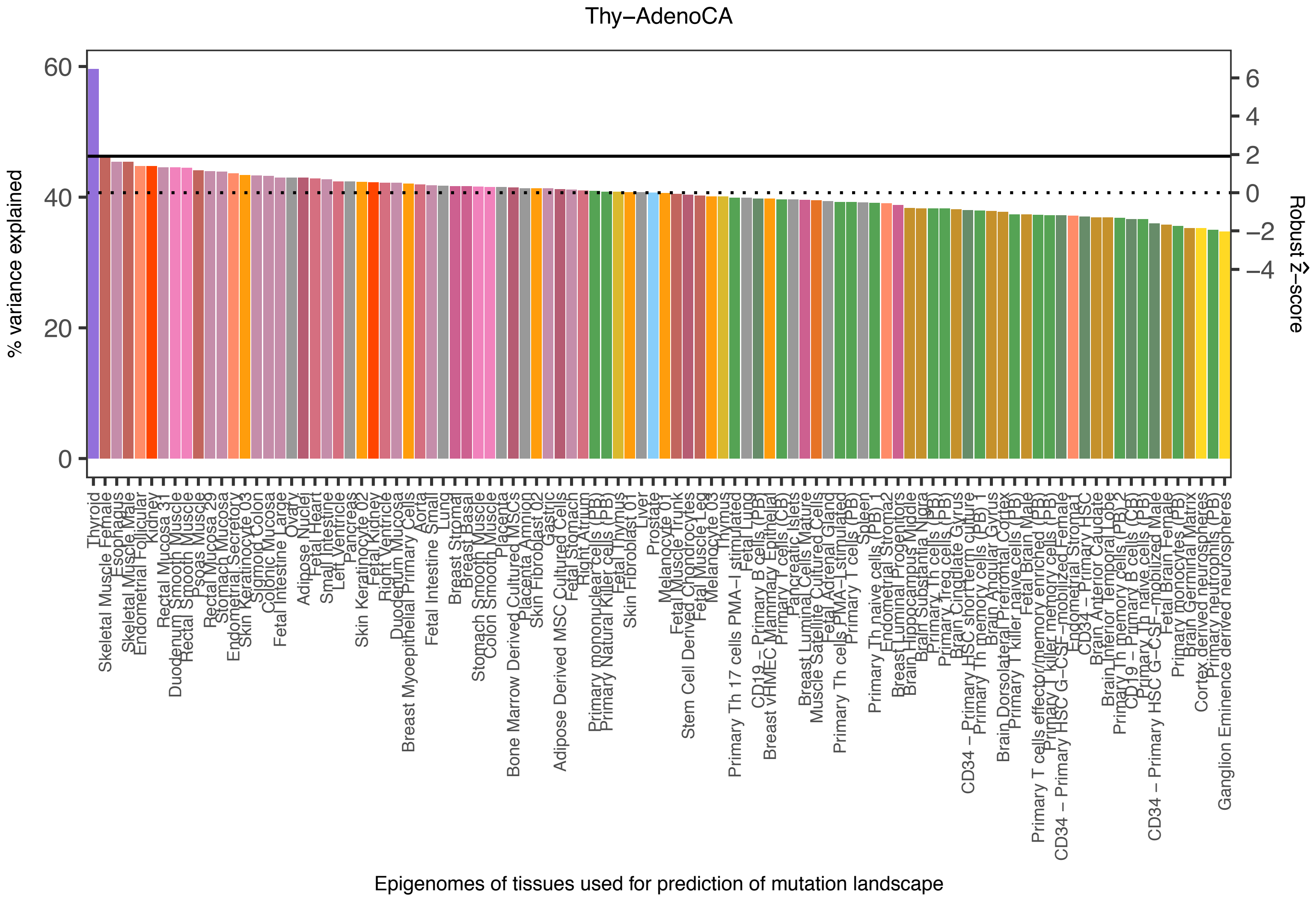

Thy−AdenoCA
% variance explained
Robust Ẑ−score
60
40
20
0
6
4
2
0
−2
−4
Epigenomes of tissues used for prediction of mutation landscape
Thyroid
Skeletal Muscle Female
Esophagus
Skeletal Muscle Male
Endometrial Follicular
Kidney
Rectal Mucosa 31
Duodenum Smooth Muscle
Rectal Smooth Muscle
Psoas Muscle
Rectal Mucosa 29
Stomach Mucosa
Endometrial Secretory
Skin Keratinocyte 03
Sigmoid Colon
Colonic Mucosa
Fetal Intestine Large
Ovary
Adipose Nuclei
Fetal Heart
Small Intestine
Left Ventricle
Pancreas
Skin Keratinocyte 02
Fetal Kidney
Right Ventricle
Duodenum Mucosa
Breast Myoepithelial Primary Cells
Aorta
Fetal Intestine Small
Lung
Breast Stromal
Breast Basal
Stomach Smooth Muscle
Colon Smooth Muscle
Placenta
Bone Marrow Derived Cultured MSCs
Placenta Amnion
Skin Fibroblast 02
Gastric
Adipose Derived MSC Cultured Cells
Fetal Stomach
Right Atrium
Primary mononuclear cells (PB)
Primary Natural Killer cells (PB)
Fetal Thymus
Skin Fibroblast 01
Liver
Prostate
Melanocyte 01
Fetal Muscle Trunk
Stem Cell Derived Chondrocytes
Fetal Muscle Leg
Melanocyte 03
Thymus
Primary Th 17 cells PMA−I stimulated
Fetal Lung
CD19 − Primary B cells (PB)
Breast vHMEC Mammary Epithelial
Primary T cells (CB)
Pancreatic Islets
Breast Luminal Cells Mature
Muscle Satellite Cultured Cells
Fetal Adrenal Gland
Primary Th cells PMA−I stimulated
Primary T cells (PB)
Spleen
Primary Th naive cells (PB) 1
Endometrial Stroma2
Breast Luminal Progenitors
Brain Hippocampus Middle
Brain Substantia Nigra
Primary Th cells (PB)
Primary Treg cells (PB)
Brain Cingulate Gyrus
CD34 − Primary HSC short term culture
Primary Th memory cells (PB) 1
Brain Angular Gyrus
Brain Dorsolateral Prefrontal Cortex
Primary T killer naive cells (PB)
Fetal Brain Male
Primary T cells effector/memory enriched (PB)
Primary T killer memory cells (PB)
CD34 − Primary HSC G−CSF−mobilized Female
Endometrial Stroma1
CD34 − Primary HSC
Brain Anterior Caudate
Brain Inferior Temporal Lobe
Primary Th memory cells (PB) 2
CD19 − Primary B cells (CB)
Primary Th naive cells (PB) 2
CD34 − Primary HSC G−CSF−mobilized Male
Fetal Brain Female
Primary monocytes (PB)
Brain Germinal Matrix
Cortex derived neurospheres
Primary neutrophils (PB)
Ganglion Eminence derived neurospheres

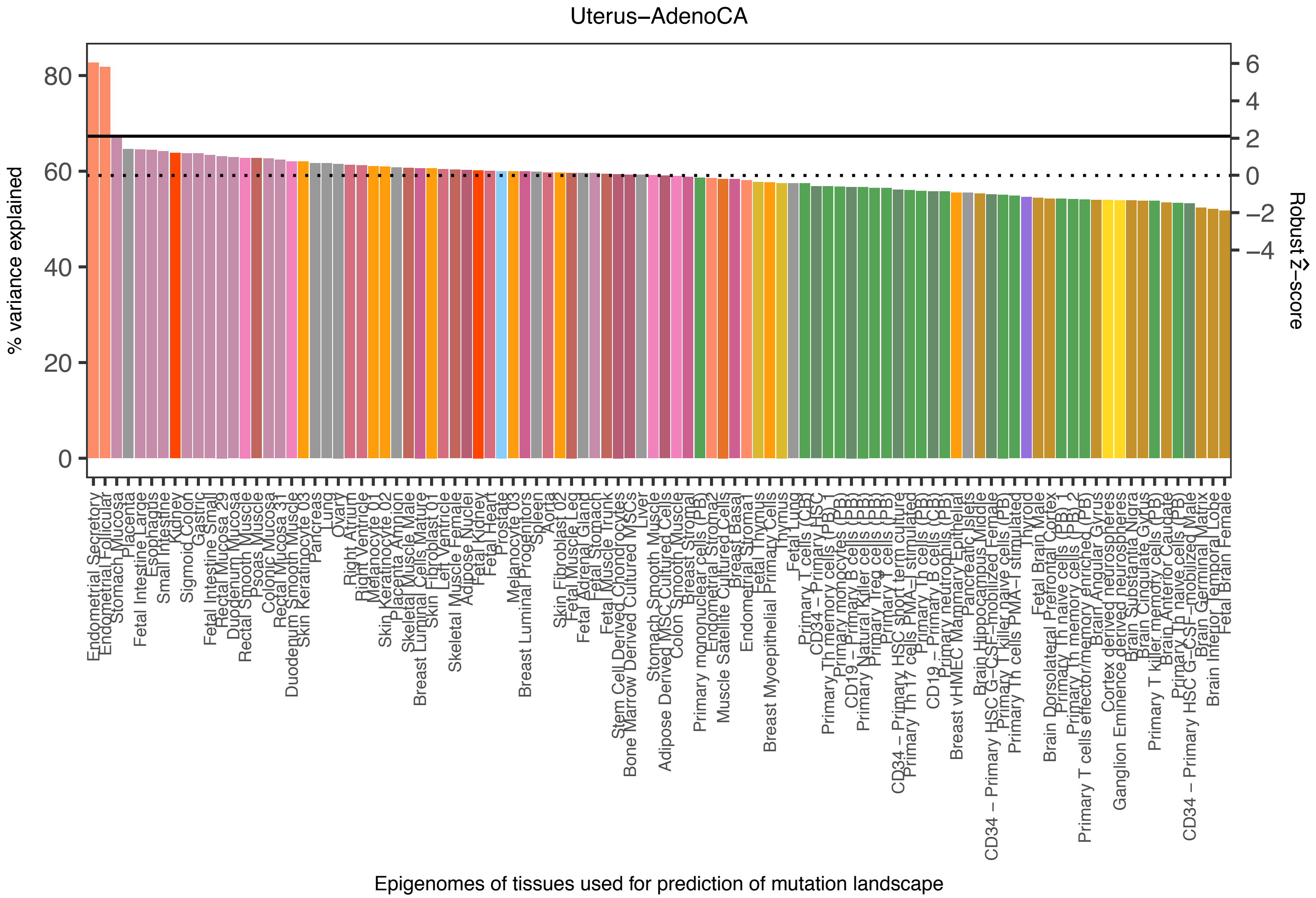
Uterus−AdenoCA
% variance explained
0
20
40
60
80
Robust Ẑ−score
6
4
2
0
−2
−4
Epigenomes of tissues used for prediction of mutation landscape

### Extended Data Figure 3

**a**

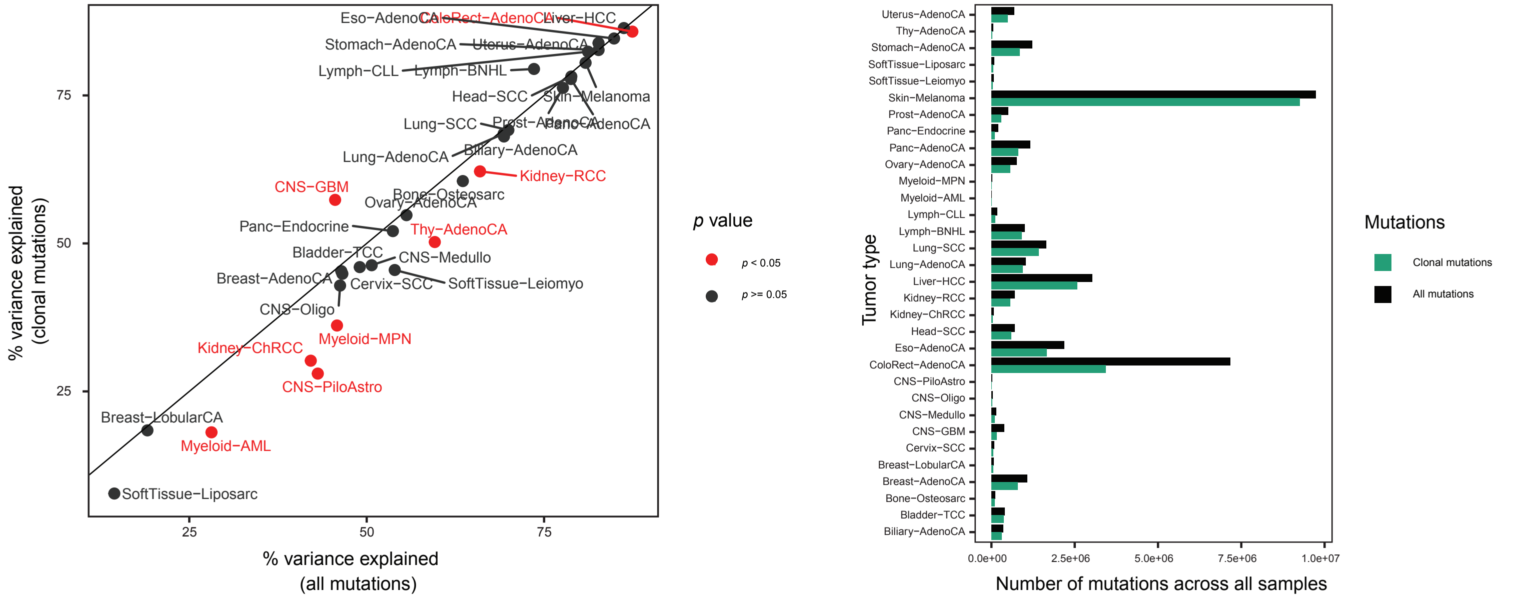


**b**

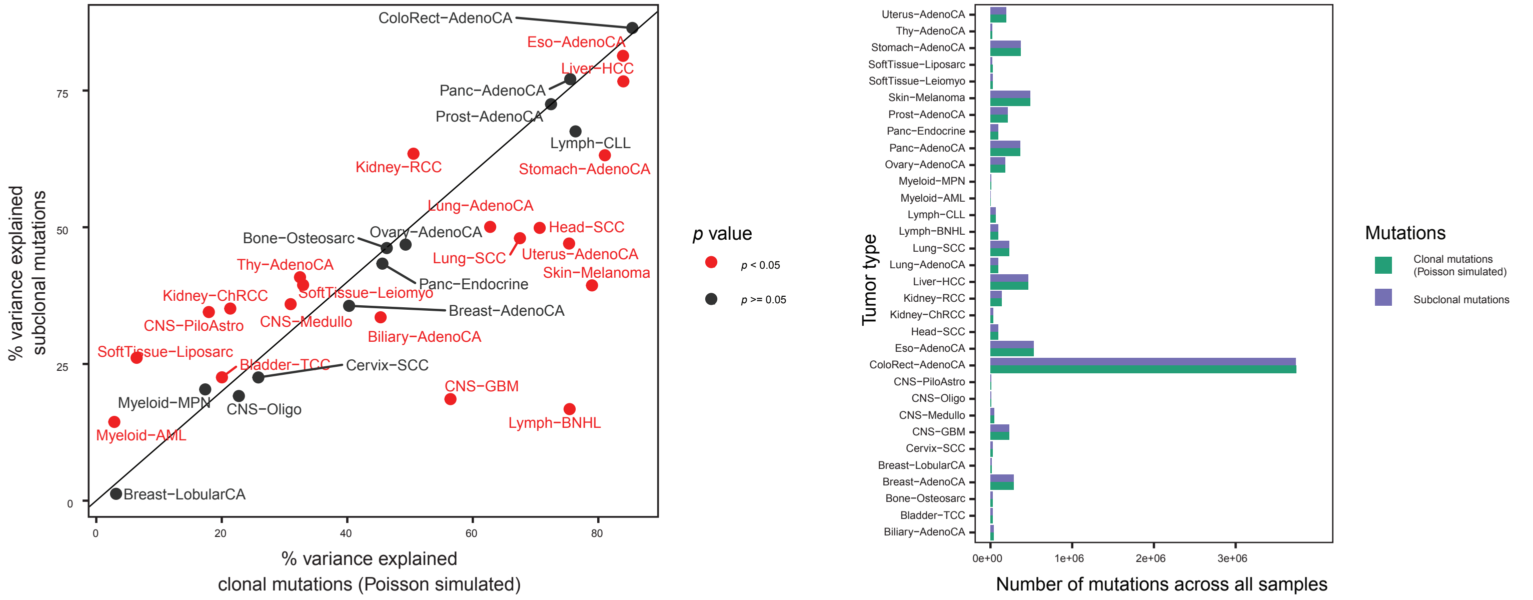

### Extended Data Figure 4

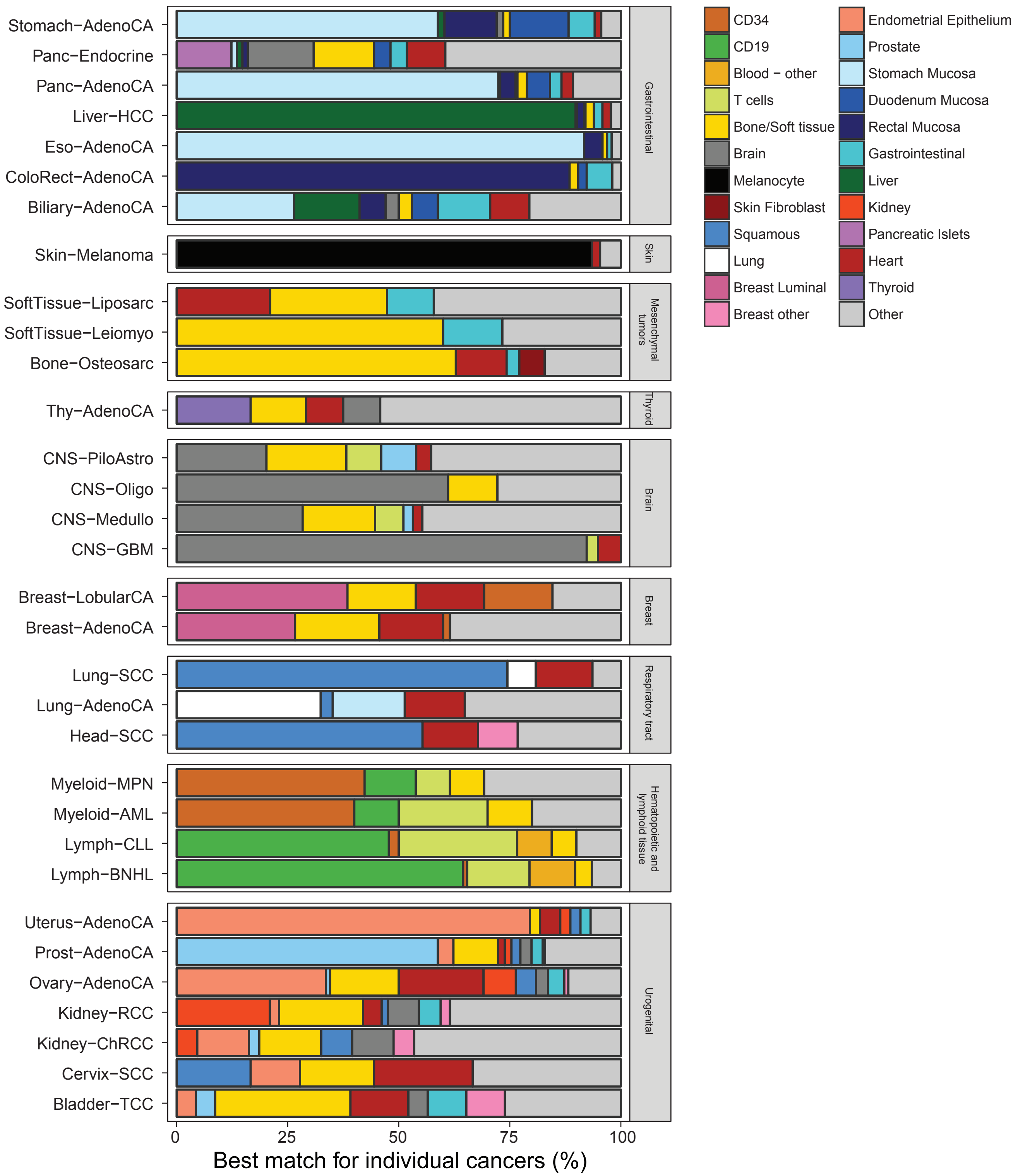

### Extended Data Figure 5

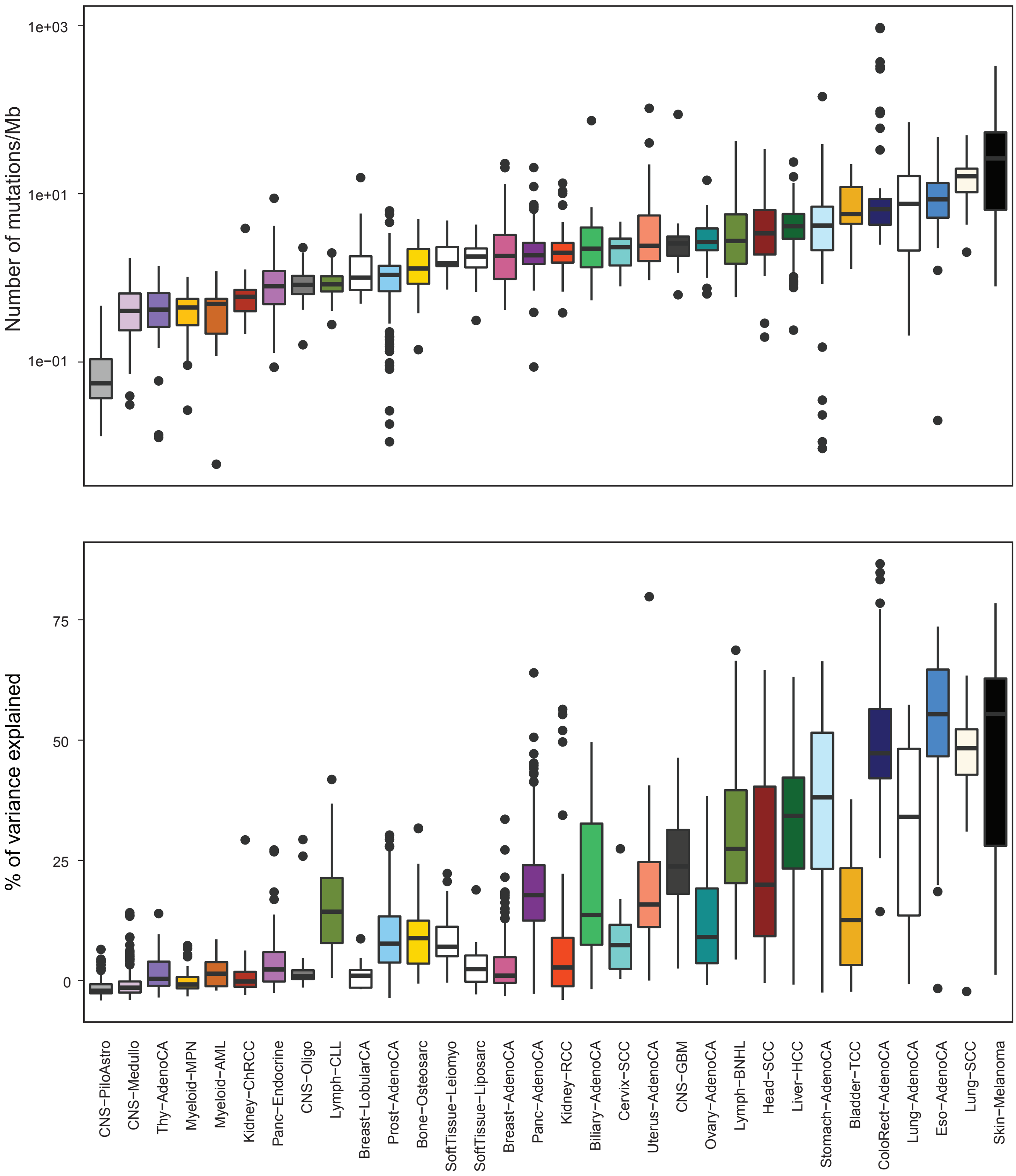

### Extended Data Figure 6

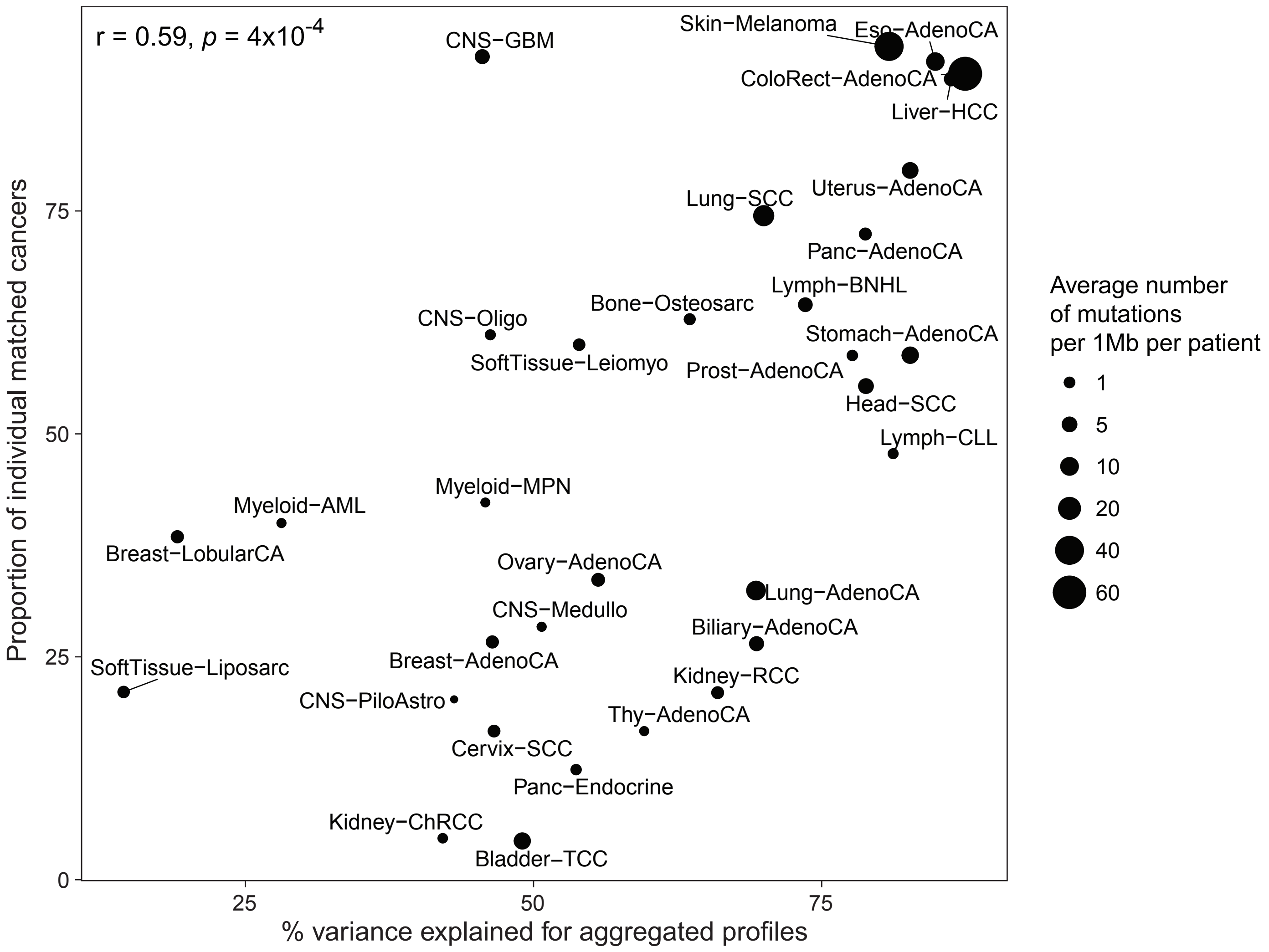

### Extended Data Figure 7

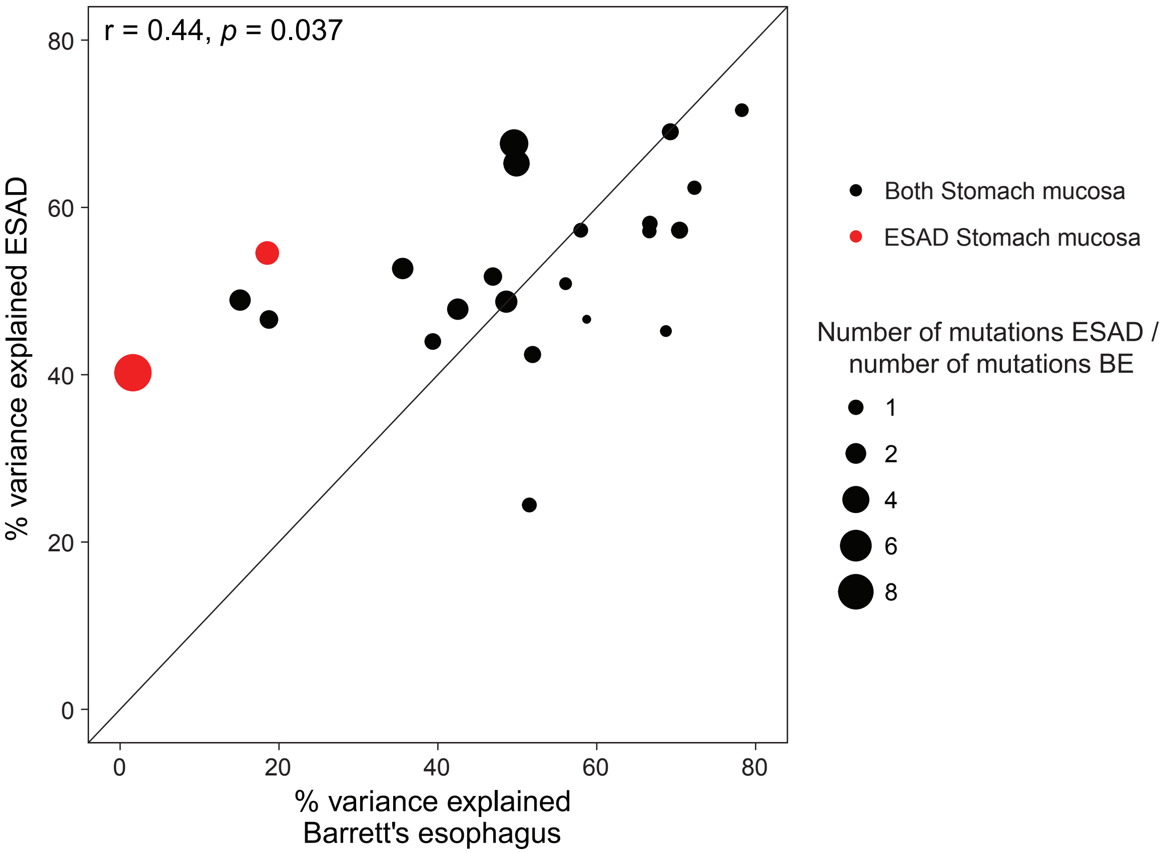

### Extended Data Figure 8

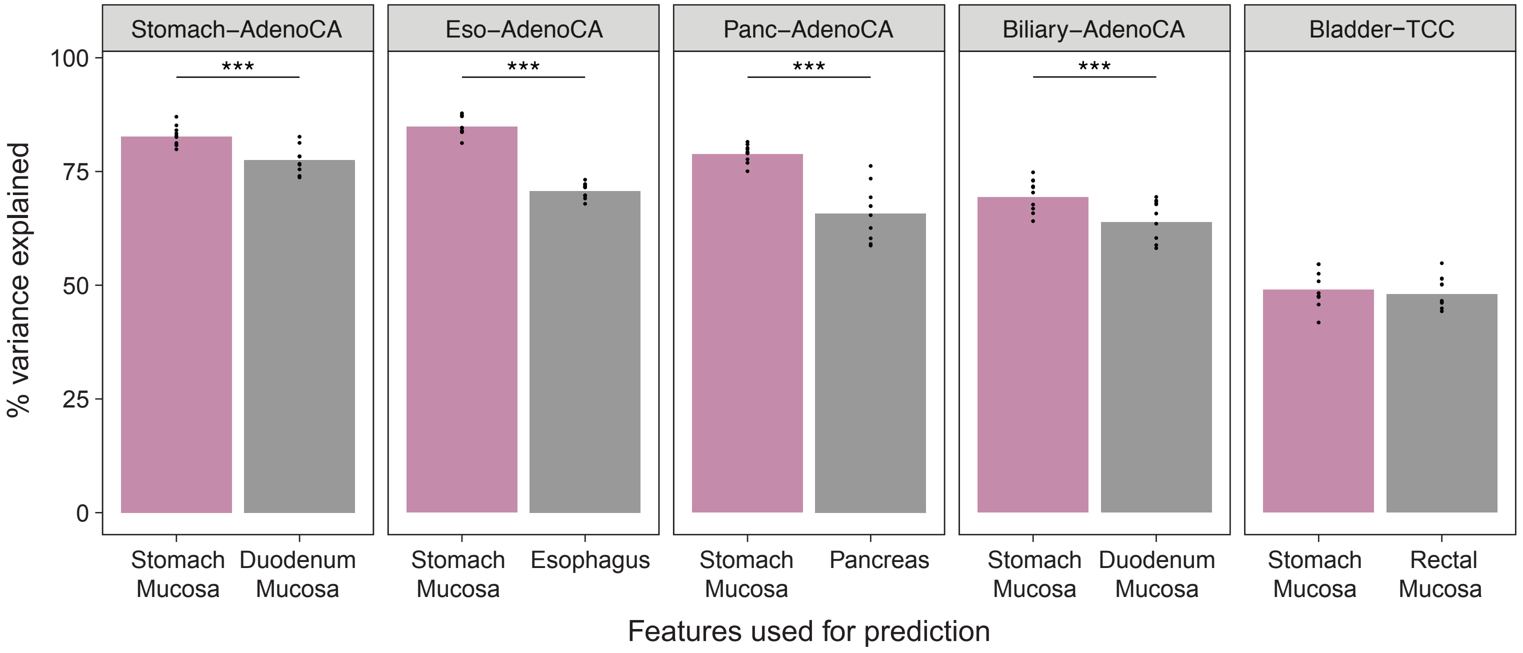

### Extended Data Figure 9

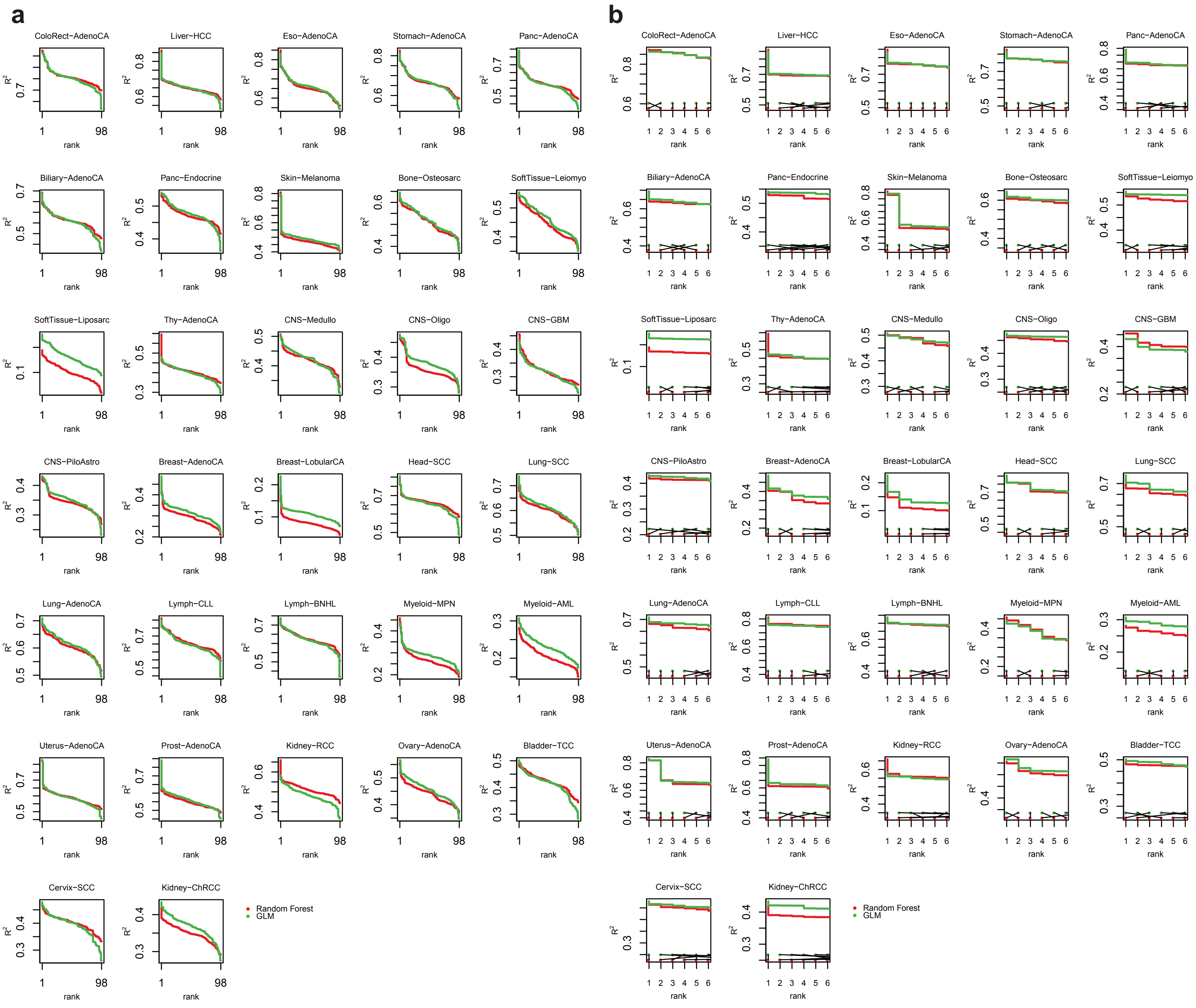

### Extended Data Figure 10

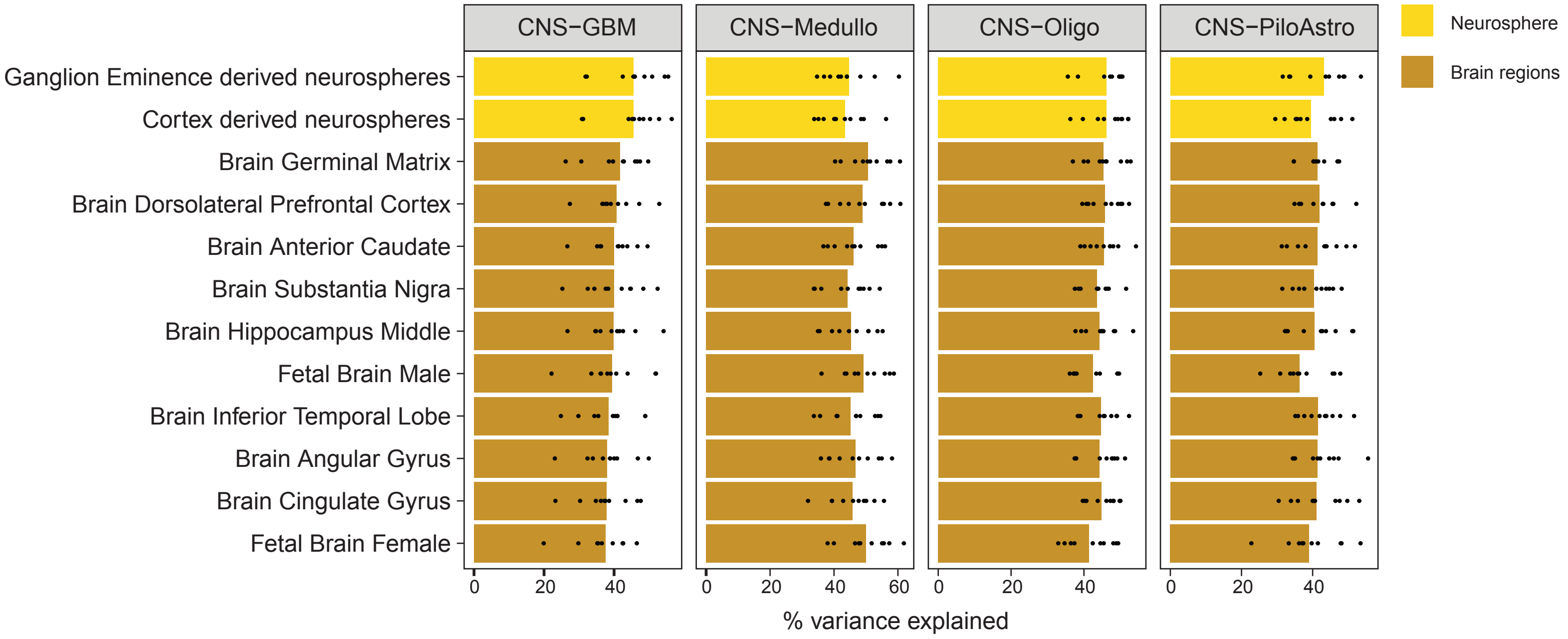

### Extended Data Figure 11

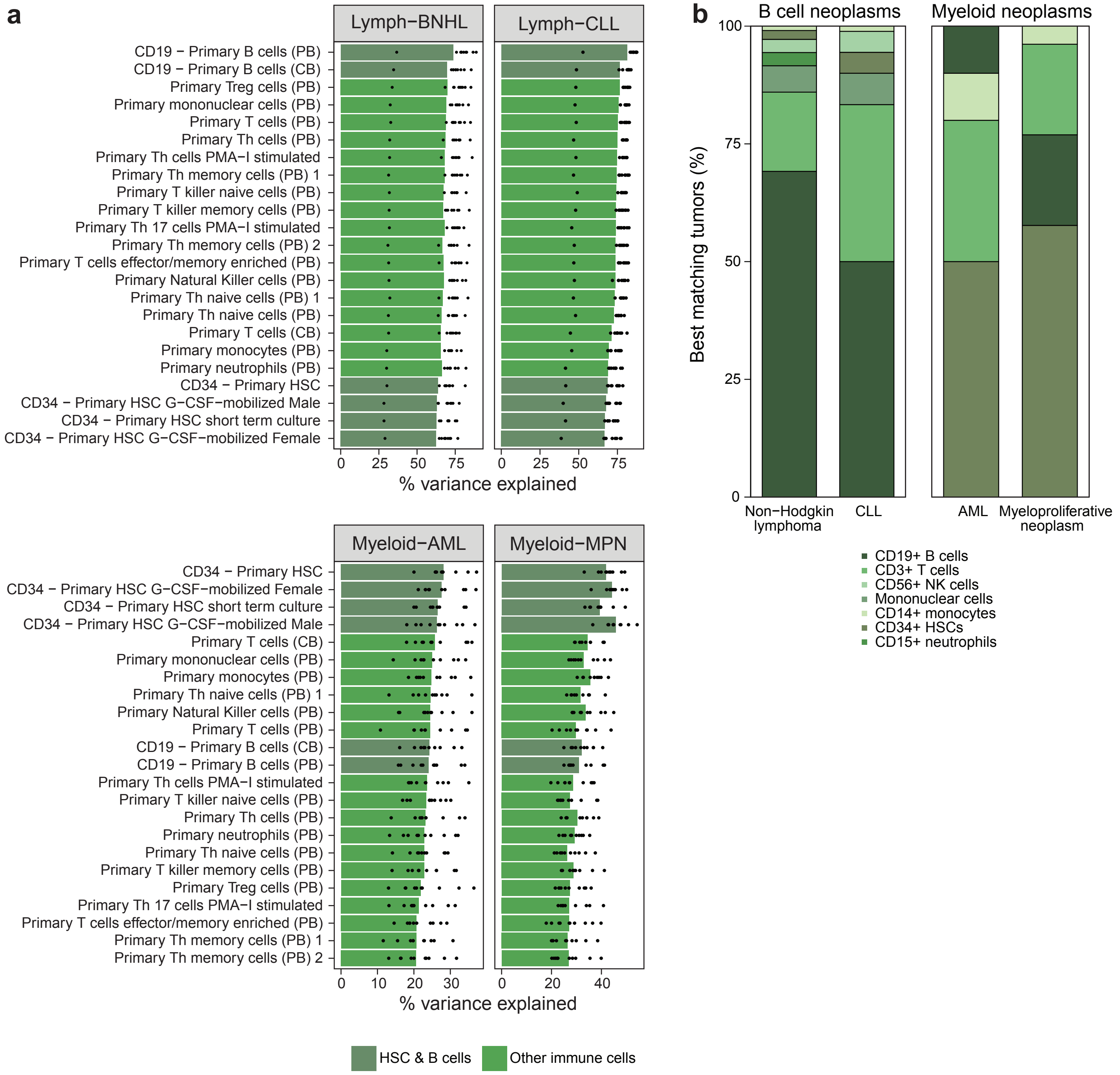

### Extended Data Figure 12

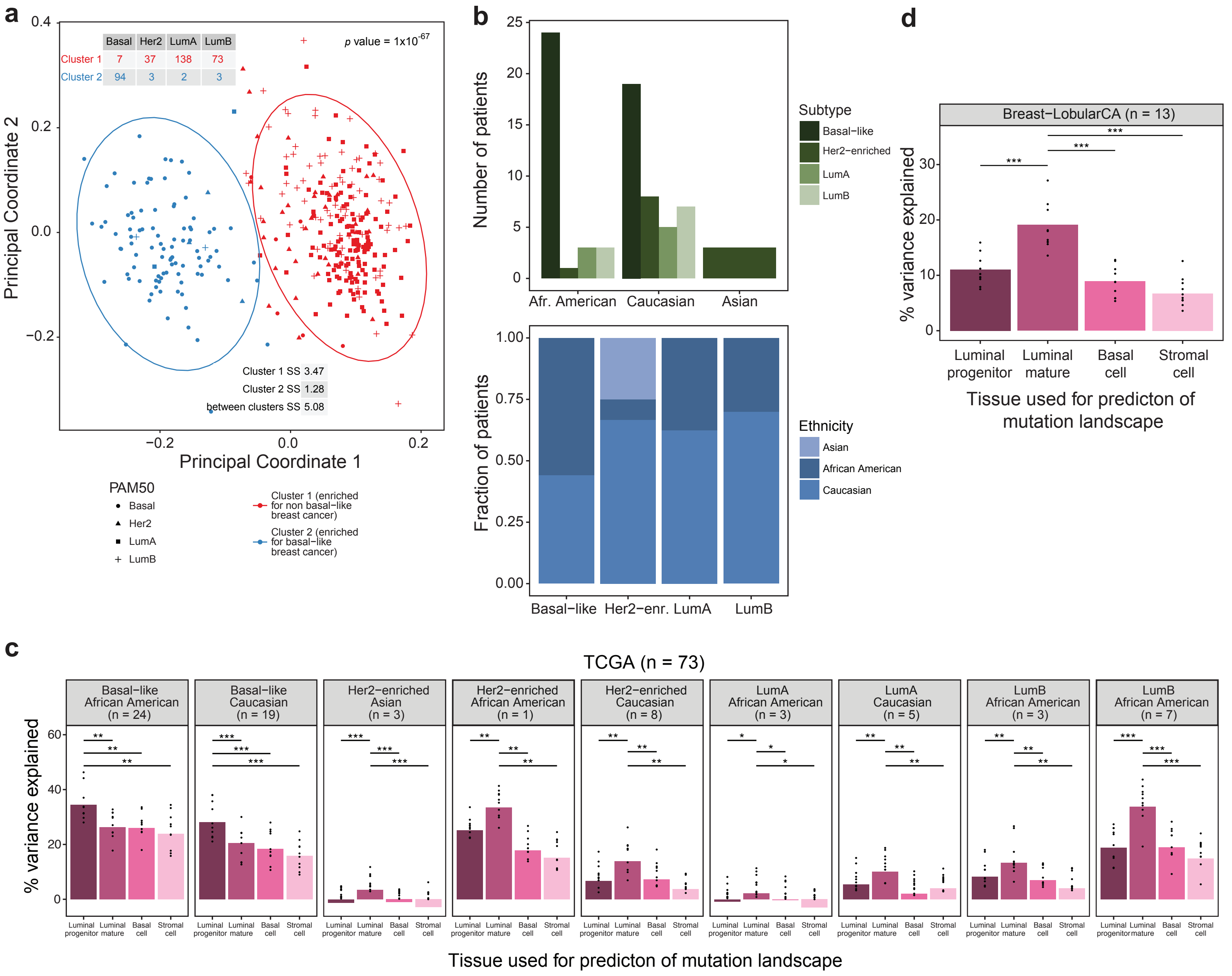

### Extended Data Figure 13

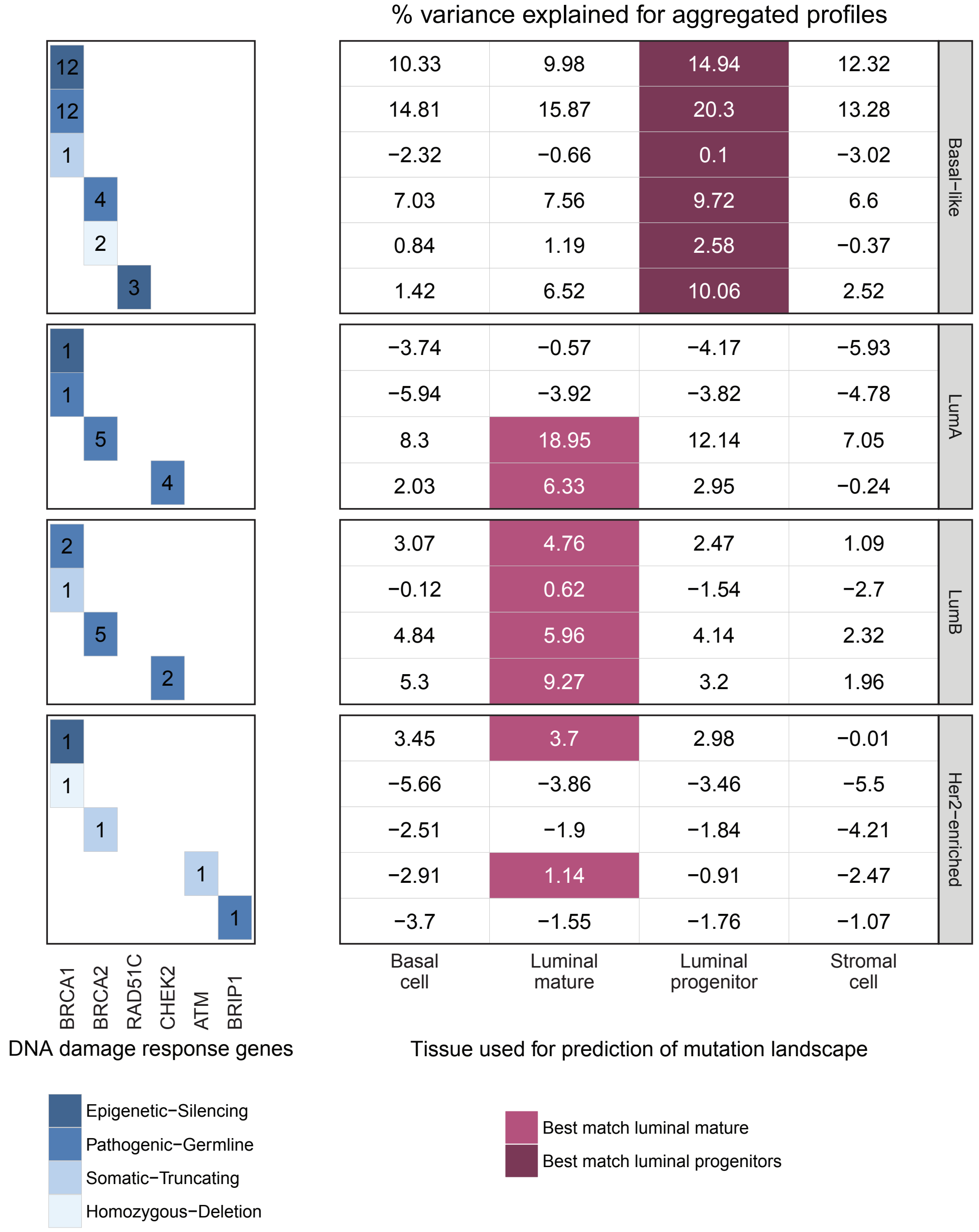

### Extended Data Figure 14

**a**

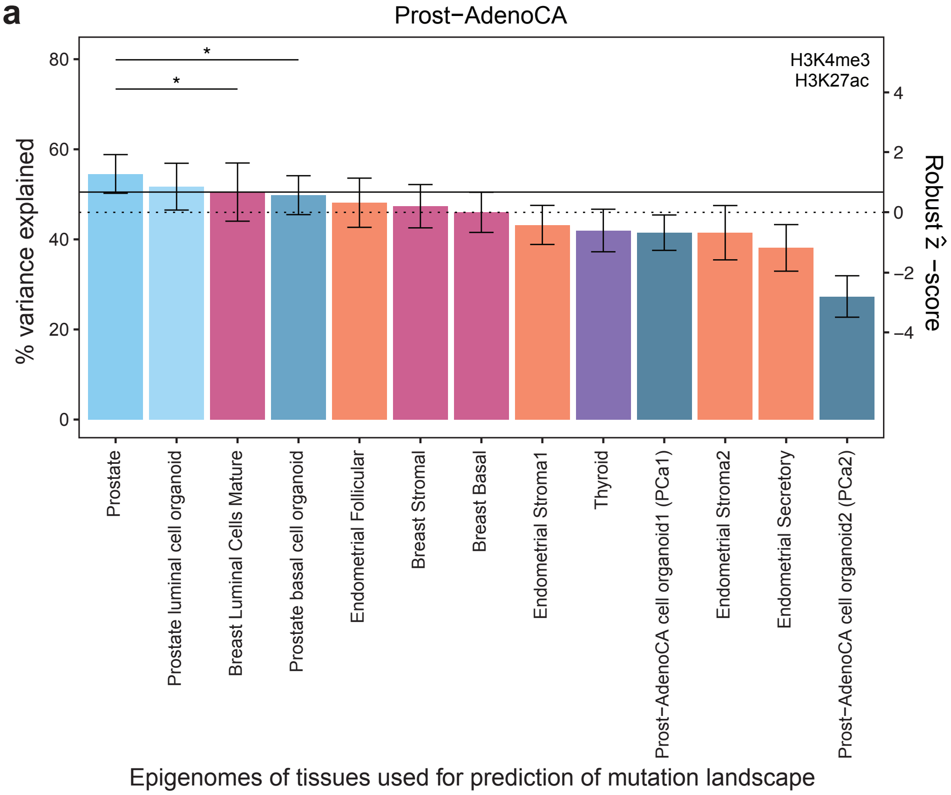


**b**

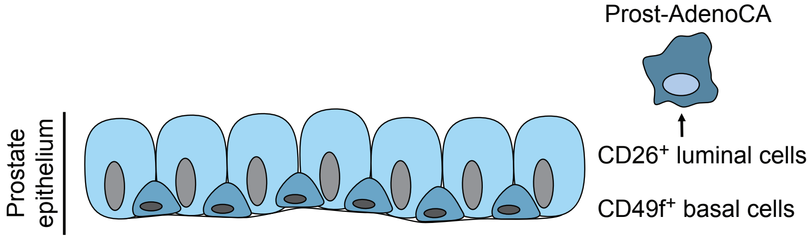

### Extended Data Figure 15

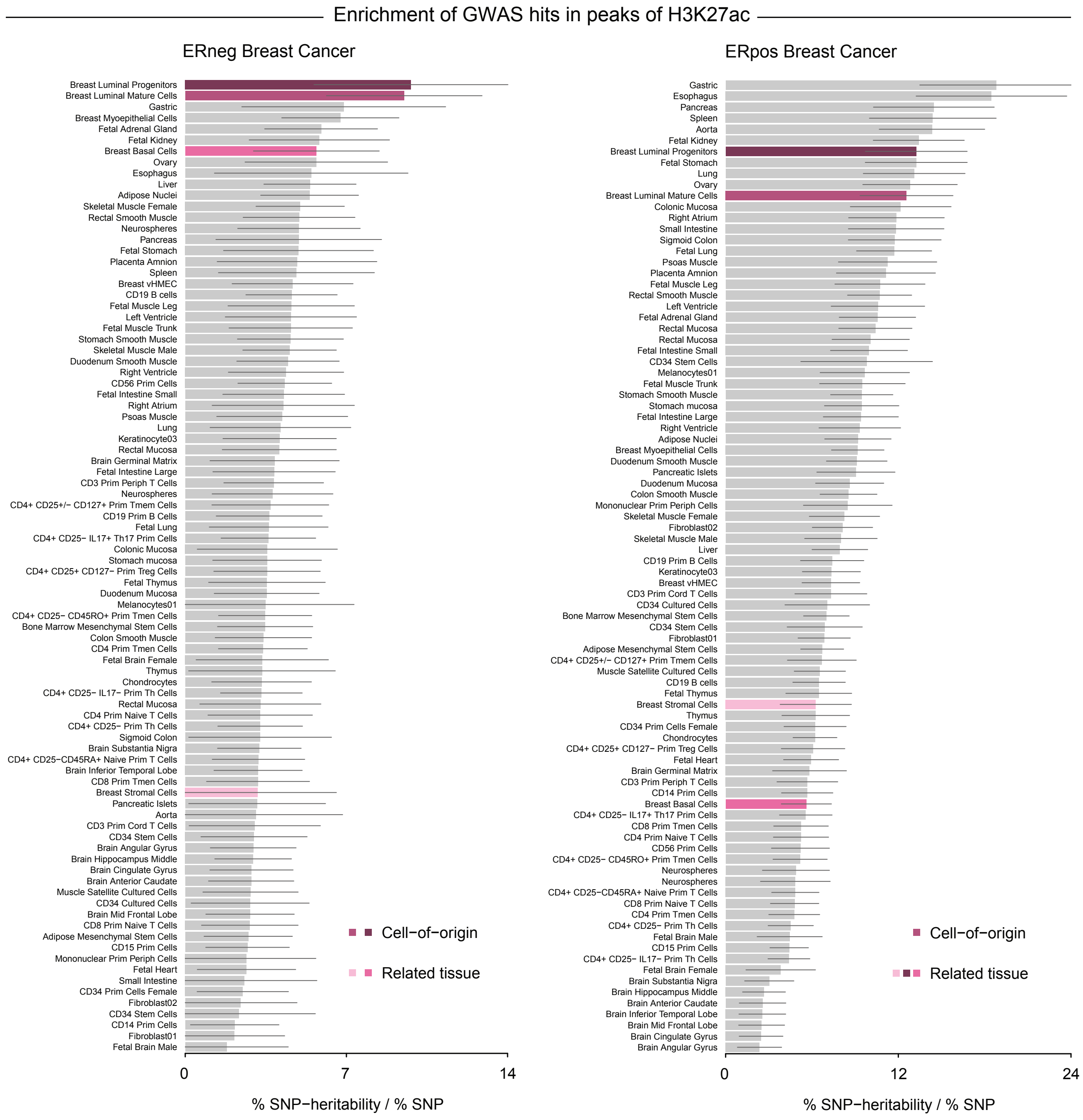

### Extended Data Figure 16

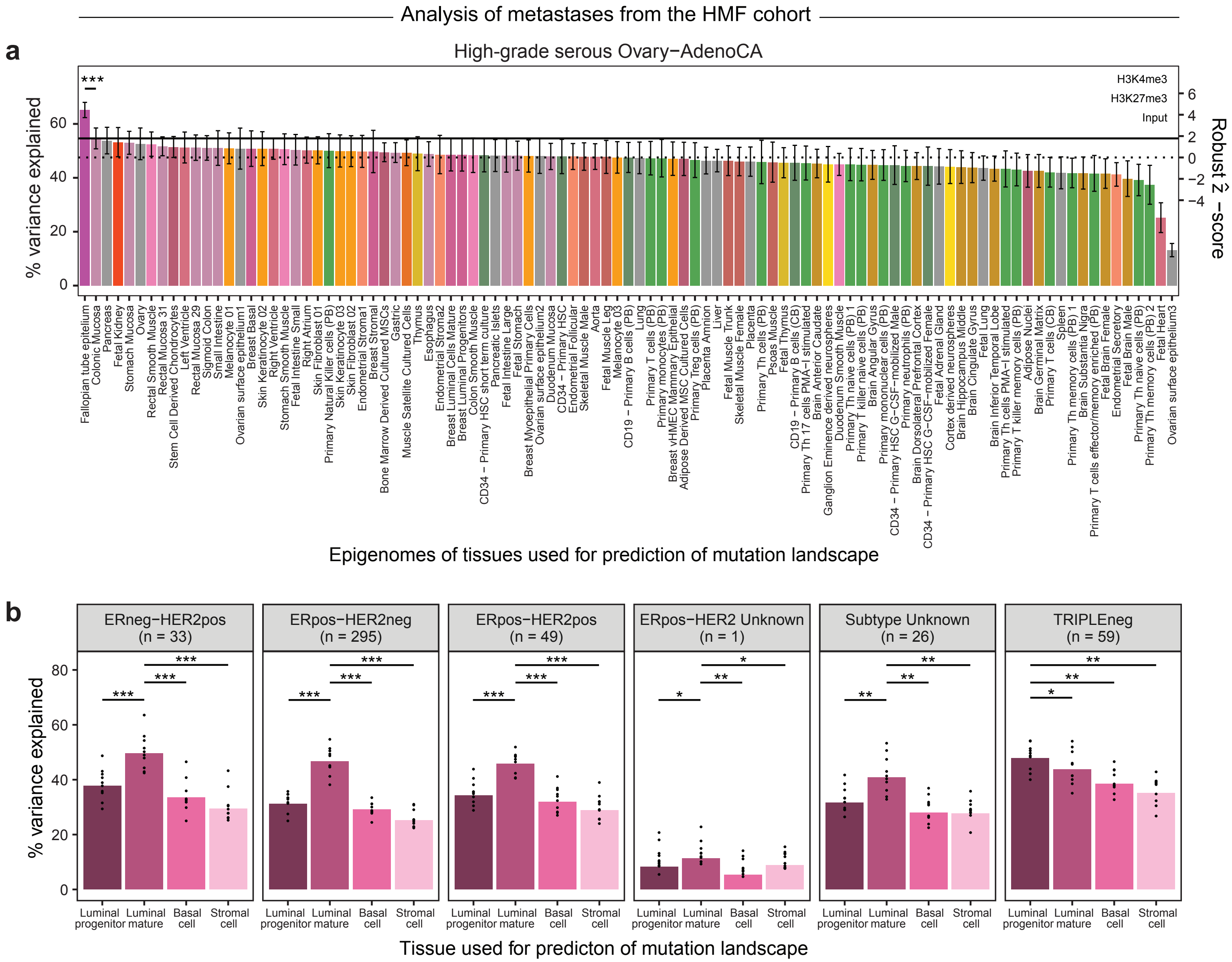

### Extended Data Figure 17

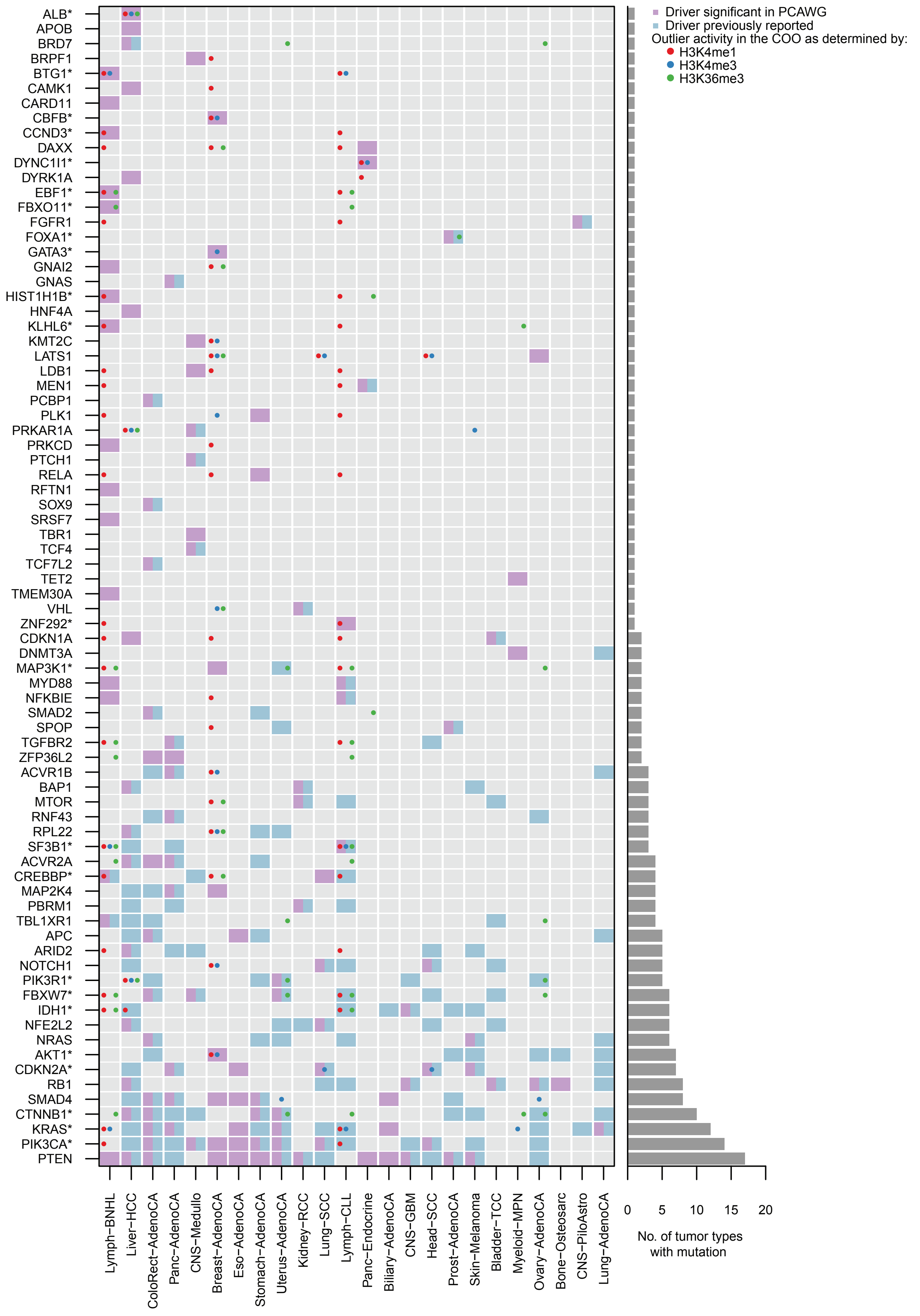

#### EXTENDED DATA INFORMATION

##### Alternative regression method gives similar results

We additionally applied a generalized linear model (GLM), which accounts for over-dispersed, Poisson-distributed data. The analysis was performed using the *caret* and *ranger* packages in 'R' and the same approach for aggregated tumor profiles as outlined for Random Forest regression (see **Methods** for details). Overall the performances of regression methods were comparable. In general, GLM performed marginally better (median delta variance explained, 0.44%) compared to Random Forest regression. However for a few tumor types Random Forest regression gave an advantage of up to 11.65% variance explained, indicating a slightly better performance (**Extended Data Fig. 9a,b**; **Extended Data Table 1**).

##### The cell-of-origin of breast cancers with homologous recombination deficiency

For analyzing the COO in breast cancer subtypes with homologous recombination deficiency (HRD), we used 60 cases from two independent datasets[1,2]. The HRD status was provided by these studies. In detail, mechanisms of inactivation included epigenetic silencing events in 17 tumors, homozygous deletion in 3 tumors, pathogenic germline variants in 36 tumors and somatic truncating mutations in 4 tumors (**Extended Data Fig. 13**). These events occurred in the following HR genes: *BRCA1* (32 tumors), *BRCA2* (17 tumors), *RAD51C* (3 tumors), *CHEK2* (6 tumors), *ATM* (1 tumor) and *BRIP1* (1 tumor). When analyzing molecular subtypes and their association with the lack of *BRCA1/2*, we discovered that tumors with a *BRCA1* mutational event were primarily basal-like (25 out of 32 tumors with *BRCA1* inactivation, 78%), while the majority of tumors with a *BRCA2* event were luminal A/B (10 out of 17 tumors with *BRCA1* inactivation, 59%), reflecting previous reports[3].

**Alternative approaches for defining outlier drivers confirm previous results**

As a robustness check, we used alternative definitions of outlier drivers and found similar results (data not shown). Drivers scoring in the top 1% top 2.5% and top 5% were deemed outliers. Again, we found a significant enrichment of driver genes in highly active chromatin regions in their COO (all *p*-values = 0.002; Fisher's exact test).

**Characterization of outlier driver genes**

To determine the biological significance of drivers with outlier activity, we searched the literature for their functional relevance and physiological role in their respective COO. Prior knowledge was available for 18 of the 29 (62%) genes.

Studies showed for 14 of the 18 (78%) annotated drivers, a crucial role in the development of their COO: (i) in breast tissue, outlier drivers have been described to be involved in the development of duct (*AKT1*[4]) and luminal cell lineages (*CBFB* [5], *GATA3* [6]); (ii) in blood cells, outlier drivers have been found to be master regulators of B cell development (*BTG1*[7], *CCND3*[8], *EBF1*[9], FBXO11[10]; *KLHL6*[11]) or hematopoietic differentiation (*IDH1*[12]; *SF3B1*[13]; *KRAS*[14], *PIK3CA*[15]); (iii) in liver tissue, *IDH1* was shown to be a key component of directing the development of hepatocytes from progenitors[16]; and (iv) in the prostate, *FOXA1*[17] was found to be required for epithelial differentiation.

In addition, a tissue-restricted expression pattern or a relevant role for tissue-specific homeostasis was reported for 4 of the 18 (22%) annotated drivers: (i) *KLHL6* was found to have a lymphoid-restricted expression across all stages of B cell development[11] and *CREBBP* to be crucial for B cell homeostasis[18]; (ii) albumin synthesis was shown to be restricted to the liver[19]; and (iii) an essential role of *CTNBB1*[20] for establishing endometrial homeostasis was suggested.

#### EXTENDED DATA FIGURE LEGENDS

**Extended Data Figure 1. Normal tissue types are depicted according to their histological relation.** Schematic illustration of anatomical sites for the 104 normal tissue types (left). The dendrogram (right) shows tissue types grouped according to their histological similarities with each group having a distinct color (see **Extended Data Table 2** for details). Cases with biological replicates are indicated by numbers; organs for which epigenetic features are available of the respective tumor type are marked with a hashtag. Abbreviations: gastroint, gastrointestinal group; urogenit, urogenital group; squam, squamous group; PB, peripheral blood; CB; cord blood; HSC, hematopoietic stem cells; MSC, mesenchymal stem cells; PMA, phorbol myristate acetate; Th cells, T helper cells; stim, stimulated; NK cells, natural killer cells; Treg cells, regulatory T cells.

**Extended Data Figure 2. The cell-of-origin can be predicted solely from DNA sequences in almost all cancer types.** Waterfall plots show the prediction accuracy of individual Random Forest regression models trained on epigenetic features from 98 normal cell types for the mutation density in aggregated profiles across 32 tumor types using windows of 1Mb. Models were trained separately for each tumor type. The prediction accuracy is measured as the percent of variance explained, calculated as the mean $R^2$ from ten-fold cross validation. Bars are colored according to **Extended Data Table 2**. Solid horizontal lines indicate performance of the second-best model based on chromatin marks from histologically unrelated cells; dashed horizontal lines depict the variance explained based on chromatin marks that represent the median tissue. The right-hand (secondary) axis shows the robust $\hat{z}$-score. Plots are ordered alphabetically by tumor type.

**Extended Data Figure 3. Clonal mutations explain more variance.** (**a**) Random Forest regression models were trained on epigenetic features from 98 cell types to predict the mutation density in 1Mb windows of aggregated tumor profiles for all mutations (x-axis) and clonal mutations (y-axis). The total number of all mutations (black) and clonal mutations (green) are shown (right). (**b**) Random Forest regression models were trained to predict the mutation density for sub-clonal mutations (y-axis) and subsampled clonal mutations (x-axis). The total number of sub-clonal mutations (violet) and subsampled clonal mutations (green) are shown (right). The paired Wilcoxon-Mann-Whitney test was used to obtain *p*-values for the comparison of $R^2$ values from the ten-fold cross validation. Data points depict the variance explained (in %) by the best performing models. Red colored dots indicate a significant change in prediction accuracy for the comparisons.

**Extended Data Figure 4. Most individual tumors best match their respective cell-of-origin or a histologically related tissue type.** Random Forest regression models were trained on epigenetic features from 98 cell types to predict the mutation density of individual samples. Bar plots show the distribution of the best-matched normal tissue types (i.e., the aggregation of all top three 'best fit' models) for individual tumors.

**Extended Data Figure 5. Prediction accuracy increases with the tumor mutation burden.** Distributions depicted include the number of mutations per 1Mb for individual tumors across cancer types (upper panel) and the maximum variance explained for individual tumors (lower panel). Tumor types are sorted according to the median count of mutations. Boxplots summarize the median, $25^{th}$ and $75^{th}$ percentiles; solid horizontal lines indicate the median; whiskers extend to the most extreme value still within 1.5*interquartile range from its nearest quartile. Points indicate outliers (defined as data points located more than 1.5*interquartile range above the upper quartile and below the lower quartile).

**Extended Data Figure 6. The best match based on aggregated tumor profiles correlates with the fraction of individual tumors matching this prediction**. The prediction accuracy (% of variance explained) of the best-performing model based on aggregated tumor profiles is shown on the x-axis. The fraction of individual tumors in which the prediction matched the prediction made from aggregated tumor profiles is shown on the y-axis. The size of the data points indicates the average number of mutations (per 1Mb) in each patient; Pearson correlation coefficient is shown.

**Extended Data Figure 7. Matched pairs of Eso-AdenoCA and Barrett's esophagus indicate a common ancestor.** Models were trained on chromatin marks to predict the mutation distribution in pairs of tumor and precursor. Prediction accuracies (% of variance explained) for Eso-AdenoCA (ESAD) and Barrett's esophagus (BE) are shown. The size of the data points indicates the number of shared mutations in Eso-AdenoCA and BE samples (per 1Mb). The color of the data points depicts whether stomach mucosa was the best match for both tumor and BE samples (black) or only for the tumor (red); Pearson correlation coefficient is shown.

**Extended Data Figure 8. Stomach mucosa is the best match for four gastrointestinal tumor types, suggesting a metaplastic switch.** Prediction accuracies (% of variance explained) of models based on stomach mucosa are shown alongside models trained on chromatin marks from (i) tissues of the same organ (Eso-AdenoCA, Panc-AdenoCA), or (ii) tissues that were second-best models (Stomach-AdenoCA, Biliary-AdenoCA, Bladder-TCC). The paired Wilcoxon-Mann-Whitney test was used to obtain *p*-values for the comparison of $R^2$ values from the ten-fold cross validation (***, $p < 0.001$; dots represent ten-fold cross-validation values).

**Extended Data Figure 9. Using another regression method yields similar results.** A generalized linear model (GLM, green) was used to prediction mutation densities for aggregated tumor profiles. Results are depicted alongside the findings from Random Forest regression (red). (**a**) The overall prediction accuracy ($R^2$) across all 98 tissue types is shown. Tumor types are ordered according to their anatomical location (as shown in **Fig. 3**). (**b**) The top 6 best matches are depicted for both regression methods across all tumor types. Any potential switch across these 6 best matches between both regression methods are depicted in the lower panels for each tumor type.

**Extended Data Figure 10. Analysis of discrete brain cells indicates a common pathway for brain tumor development.** Models were built on chromatin marks from multiple brain cell types to predict mutation density in 1Mb windows of aggregated cancer profiles of four distinct brain tumor types. Prediction accuracies (% of variance explained) are shown. Colors indicate chromatin marks derived from *in vitro* models (neurospheres, yellow) or discrete brain regions (brown). The dots represent ten-fold cross-validation values.

**Extended Data Figure 11. The analysis of discrete hematopoietic cells differentiates the COO of lymphoid and myeloid malignancies.** (**a**) Models were built on chromatin marks from hematopoietic cells to predict mutation density in 1Mb windows of aggregated cancer profiles of four distinct hematopoietic tumor types. Prediction accuracies (% of variance explained) are shown. Colors indicate chromatin marks derived from HSCs and B cells (dark green) versus all other immune cells (light green). The dots represent ten-fold cross-validation values. (**b**) Individual cancer samples were analyzed for their best matching normal cell using chromatin profiles from hematopoietic cells. Colors indicate the number of tumors that best match one of the normal cell subtypes. Color-coding is according to the distinct hematopoietic cells.

**Extended Data Figure 12. Mutation landscapes vary across breast cancer subtypes.** (**a**) Principal component analysis (PCA) was performed on mutation counts along 1Mb windows, and the resulting coordinates were clustered using k-means (k=2). The formed clusters differentiated between basal-like breast tumors (cluster 2, blue) and non-basal subtypes (i.e., luminal A, luminal B and HER2-enriched subtypes; cluster 1, red). SS depicts the sum of squared differences from the mean. Symbols denote breast cancer subtypes according to the RNA expression levels of 50 genes (PAM50, Prediction Analysis of Microarray 50) taken from TCGA[2] and Nik-Zainal et al.[1]. (**b**) Breast cancer subtypes are grouped by ethnicity (above); the fraction of ethnicities of women diagnosed with breast cancer grouped by subtype (below). Abbreviations: Afr. American, African American; Her2-enr., Her2-enriched. (**c**) Breast cancer patients were divided according to subtype and ethnicity. Models for the prediction of mutation distribution in 1Mb windows of aggregated profiles of subtypes were trained on epigenetic data from four mammary cell types. Prediction accuracies (% of variance explained) are shown. (**d**) Random Forest regression models for the prediction of mutation density of aggregated tumor profiles of Breast-LobularCA were trained on chromatin data from four normal mammary cell types. The paired Wilcoxon-Mann-Whitney test was used to obtain *p*-values for the comparison of $R^2$ values from the ten-fold cross validation (*, $p < 0.04$; **, $p < 0.007$; ***, $p < 0.001$; dots represent ten-fold cross-validation values).

**Extended Data Figure 13. Basal-like breast neoplasms are correlated with luminal progenitors, regardless of the inactivation mechanism.** The heatmap depicts the prediction accuracy (% of variance explained) across four breast cancer subtypes. Rows correspond to breast cancer subtypes stratified according to inactivation mechanisms in DNA-damage response genes (left panel with blue color scheme). Models for the prediction of mutation density in aggregated profiles were trained on chromatin marks from four normal mammary cell types (columns). The pink color scale reflects the best-matched normal cell in each category. Hierarchical clustering was used to sort both rows and columns.

**Extended Data Figure 14. Models based on luminal prostate cells perform better than those based on basal cells.** (**a**) In Prost-AdenoCA mutation density was predicted in 1Mb windows of aggregated cancer profiles using an additional set of normal tissue types restricted in the available ChIP-seq experiments (right upper corner). The paired Wilcoxon-Mann-Whitney test was used to obtain *p*-values for the comparison of $R^2$ values from the ten-fold cross validation (*, $p < 0.046$; the standard deviation across the ten-fold cross-validation values is depicted). (**b**) The schematic depicts luminal and basal prostate cells and the most likely cell-of-origin of Prost-AdenoCA.

**Extended Data Figure 15. The cell type of origin contributes to breast cancer risk.** GWAS summary statistics of ERneg and ERpos breast cancer were analyzed for the enrichment of GWAS heritability in peaks of H3K27ac. Bar plots depict the ratio of

GWAS heritability over all SNPs contained in these peaks across tissue types. Standard errors of the mean are shown. The cell subtype of origin and related breast cell types are colored; all other tissue types are marked in grey (vHMEC, variant human mammary epithelial cells; prim, primary; periph, peripheral; Tmem cells, T memory cells; Treg cells, regulatory T cells; Th cells, T helper cells).

**Extended Data Figure 16. The cell type of origin can be inferred from metastatic cancer samples.** (**a**) For fallopian tube and ovarian surface epithelium, only three ChIP-seq experiments were available (upper right corner). Accordingly, metastatic Ovary-AdenoCA models were trained on a slightly different set of 99 normal tissue types using aggregated tumor profiles. The paired Wilcoxon-Mann-Whitney test was used to obtain *p*-values for the comparison of $R^2$ values from the ten-fold cross validation (***, $p < 0.001$; the standard deviation across the ten-fold cross-validation values is depicted). (**b**) Metastatic breast cancer tumors were divided according to their hormone expression profiles. Random Forest regression models of aggregated tumor profiles of distinct were trained on chromatin data from four normal mammary cell types. For each subtype, the best match was determined, corresponding to the model with the highest prediction accuracy; *p*-values were obtained using the paired Wilcoxon-Mann-Whitney test for the comparison of $R^2$ values from the 10-fold cross validation (**, $p < 0.0$; **, $p < 0.003$ ***; $p < 0.001$; dots represent ten-fold cross-validation values).

**Extended Data Figure 17. A subset of driver genes resides in genomic regions that are uniquely active in their COOs.** Each column represents one tumor type; each row represents one coding driver gene. Colors indicate whether a gene was significantly mutated in a tumor type as identified by PCAWG (violet rectangle) or as reported previously (blue rectangle). Counts of H3K4me1 (red dots), H3K4me3 (blue dots) and H3K36me3 (green dots) peaks in the 1Mb region surrounding the driver genes were normalized as reads per million aligned (RPM). RPM values in the predicted COO were compared to the median RPM values across all tissues. Dots indicate outlier activity (i.e., 1.5*interquartile range above the upper quartile) of a driver gene in the COO of a particular tumor type. Stars are added to driver genes that provide outliers.
